# DREDge: robust motion correction for high-density extracellular recordings across species

**DOI:** 10.1101/2023.10.24.563768

**Authors:** Charlie Windolf, Han Yu, Angelique C. Paulk, Domokos Meszéna, William Muñoz, Julien Boussard, Richard Hardstone, Irene Caprara, Mohsen Jamali, Yoav Kfir, Duo Xu, Jason E. Chung, Kristin K. Sellers, Zhiwen Ye, Jordan Shaker, Anna Lebedeva, Manu Raghavan, Eric Trautmann, Max Melin, João Couto, Samuel Garcia, Brian Coughlin, Csaba Horváth, Richárd Fiáth, István Ulbert, J. Anthony Movshon, Michael N. Shadlen, Mark M. Churchland, Anne K. Churchland, Nicholas A. Steinmetz, Edward F. Chang, Jeffrey S. Schweitzer, Ziv M. Williams, Sydney S. Cash, Liam Paninski, Erdem Varol

## Abstract

High-density microelectrode arrays (MEAs) have opened new possibilities for systems neuroscience in human and non-human animals, but brain tissue motion relative to the array poses a challenge for downstream analyses, particularly in human recordings. We introduce DREDge (Decentralized Registration of Electrophysiology Data), a robust algorithm which is well suited for the registration of noisy, nonstationary extracellular electrophysiology recordings. In addition to estimating motion from spikes in the action potential (AP) frequency band, DREDge enables automated tracking of motion at high temporal resolution in the local field potential (LFP) frequency band. In human intraoperative recordings, which often feature fast (period <1s) motion, DREDge correction in the LFP band enabled reliable recovery of evoked potentials, and significantly reduced single-unit spike shape variability and spike sorting error. Applying DREDge to recordings made during deep probe insertions in nonhuman primates demonstrated the possibility of tracking probe motion of centimeters across several brain regions while simultaneously mapping single unit electrophysiological features. DREDge reliably delivered improved motion correction in acute mouse recordings, especially in those made with an recent ultra-high density probe. We also implemented a procedure for applying DREDge to recordings made across tens of days in chronic implantations in mice, reliably yielding stable motion tracking despite changes in neural activity across experimental sessions. Together, these advances enable automated, scalable registration of electrophysiological data across multiple species, probe types, and drift cases, providing a stable foundation for downstream scientific analyses of these rich datasets.

## 1 Introduction

High-density microelectrode arrays (MEAs), and in particular Neuropixels probes, have enabled simultaneous high quality recording from large populations (hundreds) of neurons with high resolution, both temporally (20-30kHz) and spatially (channels spaced by tens of microns or less)^1;2;3;4;5;6^. Since their introduction and ongoing development, high density MEAs have opened new possibilities for the study of neuronal populations via spiking activity and local field potentials, within and across brain regions. They have enabled testing a variety of novel hypotheses across species, including those related to electrophysiological^7^ and functional^8^ properties of cell types, neural correlates of consciousness^9^, population dynamics^10^, motor planning^11^, episodic memory^12^, visual decision making^13^, and skin patterning in dreaming octopi^14^. Further, Neuropixels probes have recently been employed for high-quality intraoperative recordings in humans^15;16^, both awake and under general anaesthesia while undergoing surgical interventions for their clinical care, enabling us to directly answer fundamental questions about human brain physiology with possible clinical implications.

However, several biological and physical sources of noise and variability can reduce the neural recording effectiveness of these probes^17^. In particular, in vivo recordings can be impacted by the motion of the brain relative to the recording probe, especially in recordings from human participants where brain motion effects may appear due to the heart rate, breathing, speaking, or movement of the patients^15^ and can be an order of magnitude larger than the brain movement observed in non-human animals such as mice. Such motion causes voltage measurements to drift across the recording electrodes, which can confound downstream tasks such as spike sorting and behavioral decoding. In the action potential (AP) frequency band (frequencies above ≈ 300Hz), the motion of a single well-isolated neuron relative to the probe can result in undersampling or false splits in its spiking activity if not properly motion corrected^18;19;6;20^; similarly, motion can make it difficult to identify and isolate events in the local field potential (LFP) frequency band (frequencies below ≈300Hz), leading previous studies to resort to manual or semi-automated tracking in some cases^15;16^. Further, these motion artifacts can lead to errors in downstream applications, reducing the power and accuracy of a given study’s scientific analyses and precluding full analysis of task-related activities that correlate with motion.

Estimating the motion of a sensor such as a high-density MEA from its data falls into the category of registration problems familiar from other domains, including biomedical image alignment^21;22;23;24^ and video stabilization^25;26^ among many other methods in a large and active field of research. In the context of extracellular neurophysiology recordings, registration methods need to be robust to both substantial measurement noise and the oscillations of the local field potentials and able to scale up to recordings on hundreds of channels with temporal resolution in the tens of kilohertz. Further, they must be flexible enough to model deformations of the brain tissue relative to the probe which change over time while also varying along the depth of the probe, as parts of the tissue may move differently; such spatially nonuniform motion estimation problems are referred to as “nonrigid” registration tasks, in contrast to rigid motions which do not vary along the probe depth.

Current methods rely on the motion tracking algorithm of Kilosort 2.5 (KS)^6;27^, which estimates drift from spiking activity in the action potential band using a template-based approach similar to that of the NormCoRRe (Non-Rigid Motion Correction) algorithm developed for calcium imaging data^28;26^. These methods first break the recording into independent spatial blocks (i.e., groups of channels) to account for nonrigidity and estimate motion within each block by computing a global template, which is a spatial summary of the neuronal activity computed by suitably aggregating statistics of individual spikes from across the recording into spatial bins. Next, these methods cross-correlate this global template with time-binned neuronal activity to estimate the displacement in each time bin relative to the template, leading to an estimated motion trace which can then be used to update the template in an iterative scheme. Although this method is effective in some real and simulated data^6;29^, its application is limited to datasets which can be aligned to such a global template, which excludes oscillating local field potentials and spiking data which is highly nonstationary or features drift which is large relative to the length of the probe or the spatial extent of the blocks used to account for nonrigidity. Further, KS’ motion estimate is limited in its temporal resolution by the noise characteristics of spiking data, leading to the development of algorithms to assist manual tracing at higher temporal resolution like MTracer^16^, which in addition to relying on manual annotations is also limited in its application to rigid drift (i.e., motion which does not vary along the depth of the probe).

In this work, we introduce DREDge (Decentralized Registration of Electrophysiology Data). In contrast to previous global template-based methods, DREDge starts from the decentralized framework of Varol et al. ^30^ ; Windolf et al. ^31^, which infers motion by modeling local relationships in the data, allowing for motion estimation from either time-binned spiking data or filtered local field potential recordings. This approach estimates the relative displacements of pairs of time bins via cross-correlation^28^, and models these local relationships as arising from a latent motion trace, which can then be inferred through optimization. DREDge extends this framework by posing a model which combines information from local displacements and correlations between pairs of time bins with a spatiotemporal smoothing prior, leading to a unified method which is able to produce stable motion estimates from both spikes and local field potentials. DREDge further extends this method through computational and algorithmic improvements which enable scaling to both longer and more rapidly sampled data, in particular by implementing an online algorithm that enables the inference of motion at hundreds of hertz from the local field potential band (Fig. 1). DREDge’s motion estimation runs in a small fraction of real time in the action potential band after spike detection and localization, and at around a quarter of real time when estimating nonrigid motion at high temporal resolution (∼250Hz) from local field potentials (Supp. Fig. 1).

**Figure 1.**
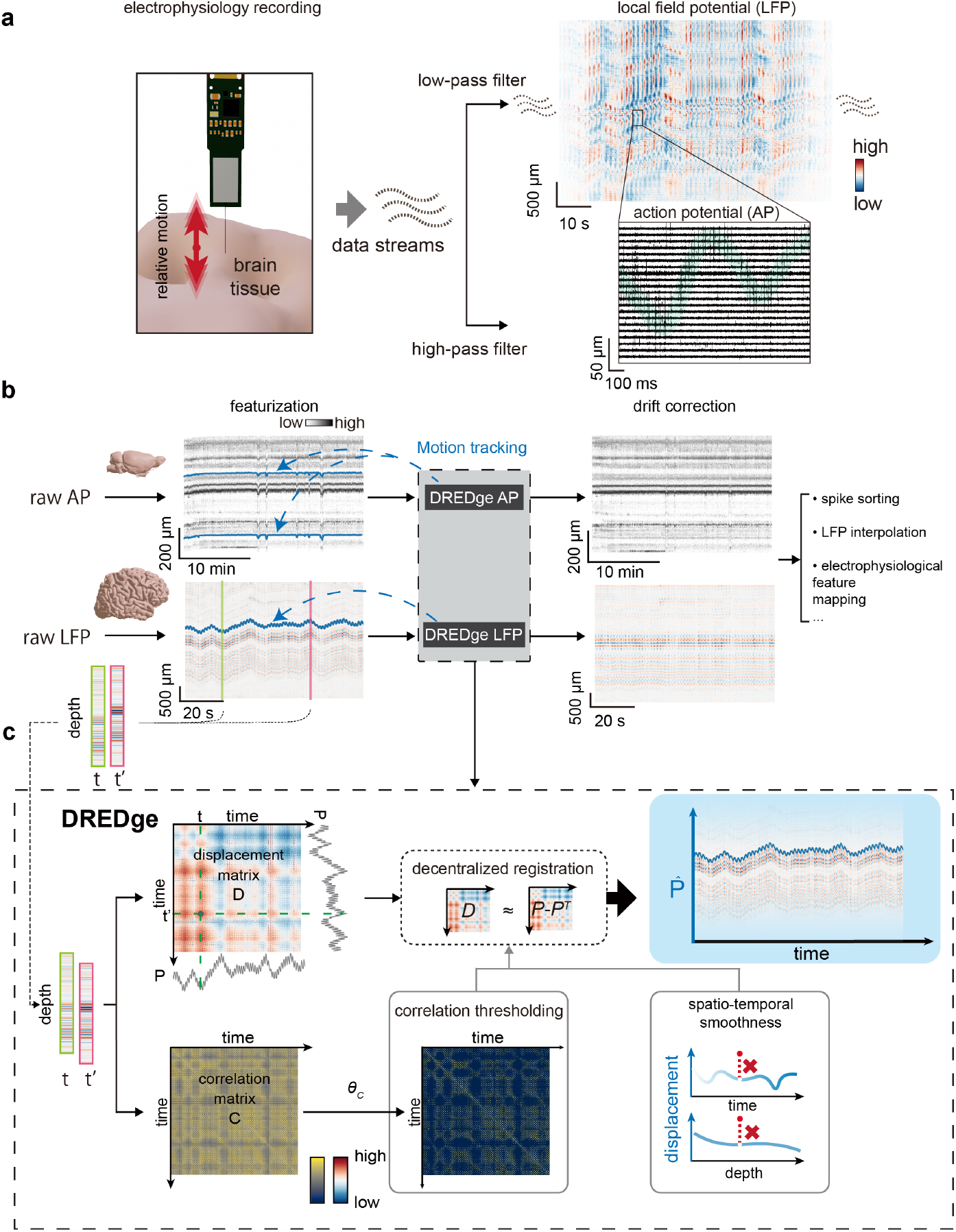
DREDge is a robust online motion drift estimating algorithm for electrophysiology recordings in both action potential (AP) and local field potential (LFP) bands. **a** Motion of the brain tissue relative to the probe causes signals to drift from channel to channel during extracellular recordings with high density multi-electrode arrays. This drift is visible in both the low-frequency local field potential (LFP; top right) band and the high-frequency action potential (AP; bottom right, green highlighting for visual emphasis) band. **b** The processing pipeline of DREDge motion estimation and analysis. Electrophysiology recordings are first preprocessed into spike rasters (here, extracted from a recording in mouse^32^; see see Section 4.1) or filtered LFP (here, from a human intraoperative recording^15^; see Section 4.2), which reveal changing structure along the long axis of the probe over time. DREDge takes in these preprocessed features and returns the drift estimate. The estimated drift is then used for drift correction that supports further analyses such as spike sorting, LFP event detection, and electrophysiological feature mapping. **c** Schematic of DREDge. Time bins of preprocessed data are cross-correlated with other time bins to generate a *T* × *T* matrix **D** of estimated optimal displacements along with the corresponding maximum cross-correlation matrix **C**. The displacement matrix **D** is filtered using a correlation cutoff, and the remaining terms are combined with a spatiotemporal smoothing prior in a bottom-up or decentralized fashion to determine drift estimates 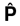 for each time bin (see Methods).

We applied DREDge to in vivo datasets from a variety of species and MEA types, including human Neuropixels recordings^15;16^, recordings in mice from the International Brain Lab’s large-scale reproducible electrophysiology experiment^32^, non-human primate recordings during probe insertion^33^, and mouse recordings using the experimental ultra high-density Neuropixels probe^34^, among others. Through these experiments, we demonstrate the usage and utility of DREDge along with some of the novel downstream analyses that it enables. These include extending motion tracking to smaller and denser probes, leveraging local field potential-based motion estimation to improve local field potential event tracking and spike sorting in human datasets, tracking electrophysiological properties of cells across tens of millimeters of brain tissue during a deep probe insertion in rhesus macaque, and enabling stable motion correction in chronic recordings with sessions separated by days or weeks. We also include detailed comparisons to current methods (i.e., Kilosort), introducing DREDge as a leading algorithm for this task.

## 2 Results

### A decentralized framework for motion estimation

DREDge is designed to estimate motion from both the action potential (AP) and local field potential (LFP) bands of extracellular recordings after suitably preprocessing them to reveal useful spatial features (Fig. 1.a and Methods). To preprocess the AP band for input into DREDge, unsorted spike events detected by existing pipelines (for example,^19;27;35^) are spatially localized relative to the probe using a model which predicts their locations from their waveforms, such as the point-source model of Boussard et al. ^36^ or alternative methods^6;37;19;38;39^. These spike positions are then combined with firing rate and amplitude information and binned in space and time to form a two-dimensional spike raster. LFP signals require less preprocessing, including spatial filtering and temporal downsampling to the target resolution for registration, along with standard filtering and artifact removal steps (Section 4.2).

After preprocessing reveals spatially localized features in the recording, our goal is to detect correlated spatial displacements of these features over time and then to use these displacements to estimate the underlying and possibly nonrigid relative motion of the probe and the brain tissue (Fig. 1.b). To that end, we began from the core operation of the decentralized framework of Varol et al. ^30^, which computes the offsets which maximize the cross-correlation between pairs of time bins of the preprocessed signal. In the decentralized framework, the motion is estimated in a bottom-up fashion from these pairwise estimates, rather than in a top-down or centralized fashion from a global template as in Kilosort’s algorithm. DREDge extends this framework, first by combining these estimates of the relative displacements between pairs of time bins with their corresponding correlations, which are used to increase the influence of pairs of time bins which contain more similar features. Displacement estimates between pairs of time bins are also excluded when the time bins lack significant signal (e.g., have very few spikes) or when the interval between time bins is large (to avoid the computational burden of cross-correlating all pairs of time bins). These observations are then placed into a Bayesian model with a spatiotemporal smoothing prior, leading to a robust and general framework which is able to estimate motion from both spikes and LFP (Fig. 1.c and Methods).

### DREDge rescues spike sorting and LFP features in human intraoperative patient brain activity

A major motivation for this work was the significant motion observed while recording human brain activity using Neuropixels probes (Figs. 2 and 3; Supplementary Video 1). As reported by two separate groups^15;16^, the brain movements during open craniotomy and deep brain stimulation surgeries are substantial, ranging up to millimeters (Fig. 2.a; Supplementary Video 1). Previous approaches to combat and correct for this motion signal primarily involved manual tracking in the local field potential^15^ or action potential bands^16^ or semi-automated tracking^16^ (MTracer, https://github.com/yaxigeigei/MTracer). In a collection of both openly shared deidentified data sets and newly collected data sets, we demonstrate the capability of DREDge to automatically track this movement within the neural signal both in the LFP band and the spiking activity or AP band (Fig. 2.a, right panel). In a subset of cases (*N* = 3), we compared DREDge’s LFP-based tracking to manual tracking using LFP signals (Supp. Fig. 2;^15^). We found a high correlation between manual tracking and DREDge motion tracking (Pt01, *r* = 0.98; Pt02, *r* = 0.99; Pt03, *r* = 0.85; Pearson’s *r* ; *p* < 0.000001 for all three instances, Supp. Fig. 2). Further, we found that the peaks in the power spectra for the manual and DREDge-tracked motion were in agreement. Finally, in an attempt to validate whether this movement tracked using neural signals corresponds to actual movement, we performed motion tracking of pixels in a video of the brain movement in an open craniotomy and found that the video-tracked movement and its spectral peaks were very similar to those of both the manual and DREDge motion tracked traces (*N* = 1; Supp. Fig. 2).

**Figure 2.**
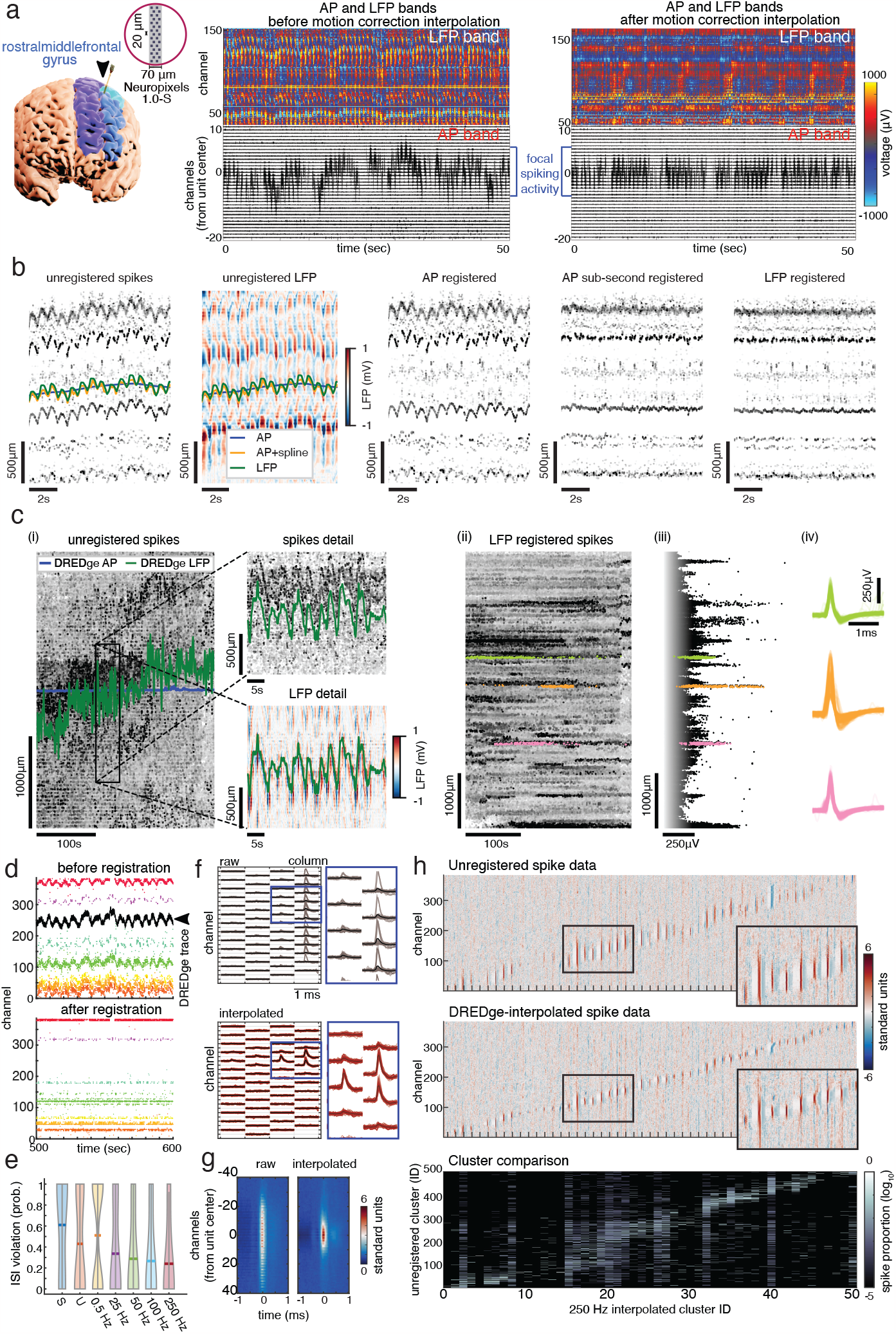
Correcting for motion in human spiking data. **a** In a recording^15^ in the rostral middle frontal gyrus (also the dorsolateral prefrontal cortex, shown in this participant in the brain reconstruction on the left), the neural signal before (middle) and after applying interpolation (right) to correct for the motion in the local field potential (LFP) and action potential (AP) bands based on DREDge’s motion tracking in the LFP band. Brain regions in figure on the left: rostral middle frontal gyrus, blue; caudal middle frontal gyrus, cyan; superior frontal gyrus, purple. Arrow indicates location of the Neuropixels probe. **b** DREDge’s LFP-based tracking accurately tracks motion which can be independently identified from spiking information alone. Fast breathing- and heartbeat-induced motion present in a human intraoperative recording is visible in spike and LFP rasters (i,ii). DREDge’s lower temporal resolution spike-based tracking finds and corrects the slow motion trend (i, blue; iii), while the LFP-based estimate (i and ii, green; v) tracks the fast oscillations. Sub-second correction on top of AP-based tracking based on clustering and splines matches well with the LFP-based method (i and ii, orange; iv; see Section 4.8). **c** Recovering units in noisy spiking data by motion estimation from the local field potential (LFP) band: although the large and rapid motion in this recording leads to a spike raster from which DREDge cannot extract a signal (i), using DREDge’s LFP-based non-rigid motion estimation to correct the positions of spikes reveals well-isolated single unit waveforms (iv) in groups of spikes collected by isolating clusters in plots of spike depths vs. time and amplitude (ii and iii). **d** A subset of spike detections and sorted units (with different single unit clusters color coded as dots) across channels before (top) and after (bottom) registration with a DREDge motion estimate (black line). Note the emergence of aligned spikes on the bottom panel. **e** Progressive decrease in inter-spike interval violation probability with increasing interpolation rate (0.5 - 250 Hz), as compared to unregistered data (U) and data interpolated using the motion-correction interpolated method based on a randomly permuted or “scrambled” DREDge motion estimate (S). Bar represents mean. **f** Representative unregistered (top) and 250 Hz-interpolated (bottom) unit (red dot on panel D), revealing a well-stereotyped multi-channel waveform after interpolation. Scale bar 1 ms. **g** Average spatial distribution of spike clusters when non-interpolated (left) and 250 Hz-interpolated (right); motion-correction interpolation concentrates spike power around a central channel. **h** Full probe spatial distribution of spikes in non-interpolated condition (top) and 250 Hz-interpolated clusters (middle). (Bottom) Comparison of 250 Hz interpolated spike assignments to unregistered clusters, showing over-splitting and cross-contamination of unregistered clusters.

To further validate this cross-band registration procedure, we examined another human recording^15^ with fast drift. In this dataset, DREDge’s AP-based motion estimation was able to capture the slow trend of the true motion, but not the faster motion due to heartbeats and other sub-second brain motions. Since this recording featured prominent and well-isolated spiking activity traces from probable single units, it was possible to estimate the trajectories of these point clouds in order to refine the motion estimate at higher temporal resolution. To do so, we used a rough clustering to isolate each of these units’ traces, and used the spike positions within these clusters to fit a spline (Fig. 2.b, Supp. Fig. 3; see also Section 4.8). This sub-second AP-based motion correction procedure was able to track the fast (< 1Hz period) heartbeat-induced motion visible in the modeled spike positions and LFP raster (Fig. 2.b, i and ii), leading to an apparent improvement of its registered spike raster (iv) over that of DREDge’s AP-based estimate (iii). Next, we applied DREDge’s LFP-based motion tracking to the same recording. We found that the motion traces estimated using the sub-second correction method and DREDge-LFP overlapped strongly (Fig. 2.b, i and ii) and that the LFP-registered spike raster (v) was visually aligned with the sub-second corrected raster (iv). This agreement reinforced the utility of applying LFP-based motion estimates to realign spike data while also validating the alternative spline-based method for estimating sub-second rigid motion from clustered spikes.

DREDge’s ability to track motion from both the AP and LFP bands allows users to choose the best signal source in each application. For instance, in some human recordings featuring large natural heartbeat- and breathing-induced motion which is fast relative to the timescale at which AP motion tracking is stable, which is typically around 1Hz due to the sparsity of spiking activity, motion tracking in the AP band can be unreliable or impossible, corresponding visually to a lack of structure in the spike raster plot (Fig. 2.c, i). However, we found that motion tracking in the spatiotemporally smooth LFP band was consistently reliable in such datasets, even when performing nonrigid registration at high temporal resolution (250Hz). In Fig. 2.c, we focused on a human Neuropixels 1 recording made with a long two-column channel configuration^16^, featuring thousands of microns of drift across the entire recording made up of fast motion oscillations of approximately 500μm around a long-term drift which extended over approximately 1mm. LFP-based nonrigid motion estimation visually appeared to track fast moving features present not only in the LFP band but also in scatter plots of spike positions (Fig. 2.c, i, detail plots). When visualizing the positions of spikes after correction using the nonrigid LFP-based motion estimate in scatter plots versus time (Fig. 2.c, ii) and spike amplitude (Fig. 2.c, iii), isolated clusters of these spike positions became apparent. Waveforms extracted from the detected events leading to the spikes visualized in these scatter plots had well-stereotyped shapes Fig. 2.c, iv), indicating that the LFP-based motion estimate was able to stabilize the positions of single units, validating the utility of cross-band registration in the estimation of extensive and fast drift in a dataset which would be challenging or impossible to process based on AP data alone. (Similar results are illustrated in Supp. Fig. 4.)

On the other hand, cases exist where the LFP band does not feature structures which can be used in motion tracking, or in which other signals dominate, making LFP-based motion tracking impossible. For instance, in recordings from ketamine/xylazine-anaesthetized rat^40^ (Supp. Fig. 5), the LFP band is dominated by slow-wave activity across the array that confounds DREDge’s LFP-based motion tracking, leading to an artifactually oscillating motion estimate which did not align with the very stable spike raster plot. However, when we applied spike-based motion tracking to these recordings, the estimated motion trace was very stable, in agreement with the apparent lack of drift in the spike rasters. The flexibility of the DREDge algorithm made it possible to switch between these modalities as required by each application.

As above, we found that in multiple other recordings (*N* > 20 in human cortex) the brain motion could be observed in both the changing voltages across the channels in the LFP and the identifiable single-unit waveforms moving up and down the channels in the recording (Fig. 2.d)^15^, and that tracking the motion in the LFP band using DREDge and then interpolating the voltage values in both the AP and LFP bands was able to compensate for this motion (Fig. 2.d, bottom panel). We hypothesized that this motion correction procedure would lead to marked improvements in the quality of single units isolated by spike sorters. Indeed, not only did this correction stabilize the location of detected spike waveforms, but the subsequent sorted single unit clusters were better isolated with decreased inter-spike-interval (ISI) violations (Fig. 2.e), more concentrated waveforms across channels per individual cluster (Fig. 2.f-g), and reduced oversplitting and contamination across clusters (Fig. 2.h). We found that the spatial spread of the voltages was concentrated in a smaller range with significantly higher amplitudes represented in a smaller spatial range following motion corrected interpolation compared to the raw data set (Supp. Fig. 6, two-sided two sample *t* -test at each distance from center, Bonferroni corrected with threshold *p* < 0.05). Importantly, the sorted clusters improved (had fewer ISI violations) as we increased the temporal resolution of LFP-based DREDge motion tracking from 1Hz to 250 Hz. To further demonstrate improved spike sorting results, we examined the relationship between sorted clusters before and after correcting for the tracked motion (Fig. 2.h). The number of sorted clusters (or single units) decreased from more than 500 to around 50. Visualizing the overlap between unregistered and registered units revealed that the unregistered clusters tended to comprise spikes from several of the registered clusters, indicating oversplitting relative to the improved clustering after registration.

LFP-based motion estimation and the following interpolation step can also be applied to correct for motion artifacts in the LFP band itself, leading to cleaner and more stable LFP signals (Fig. 3.a). However, even after this step, there was still a clearly visible heartbeat artifact in the signal, which is commonly observed in electrophysiological recordings (see, e.g., Tal and Abeles ^41^) and which manifested as large low-frequency peaks in the power spectrum. To remove this artifact from the traces after motion-correction interpolation, we applied Zapline-plus^42;43^, a generalized line-noise removal method which uses spectral and spatial filtering to effectively remove specified, narrow-band oscillatory components from the signals (see also Supp. Fig. 7). We targeted the low-frequency peaks in the signal, and in particular those which matched the spectral peaks in the DREDge motion trace. This additional step resulted in smoothed LFP signals similar to those which we observe in microscale laminar sampling of human cortical layers using other types of electrodes (Ulbert et al. ^44^ ; Csercsa et al. ^45^ ; Fig. 3.a). Further, when we examined the power spectra across channels, we found peaks before motion correction which disappeared after motion correction (Fig. 3.a).

**Figure 3.**
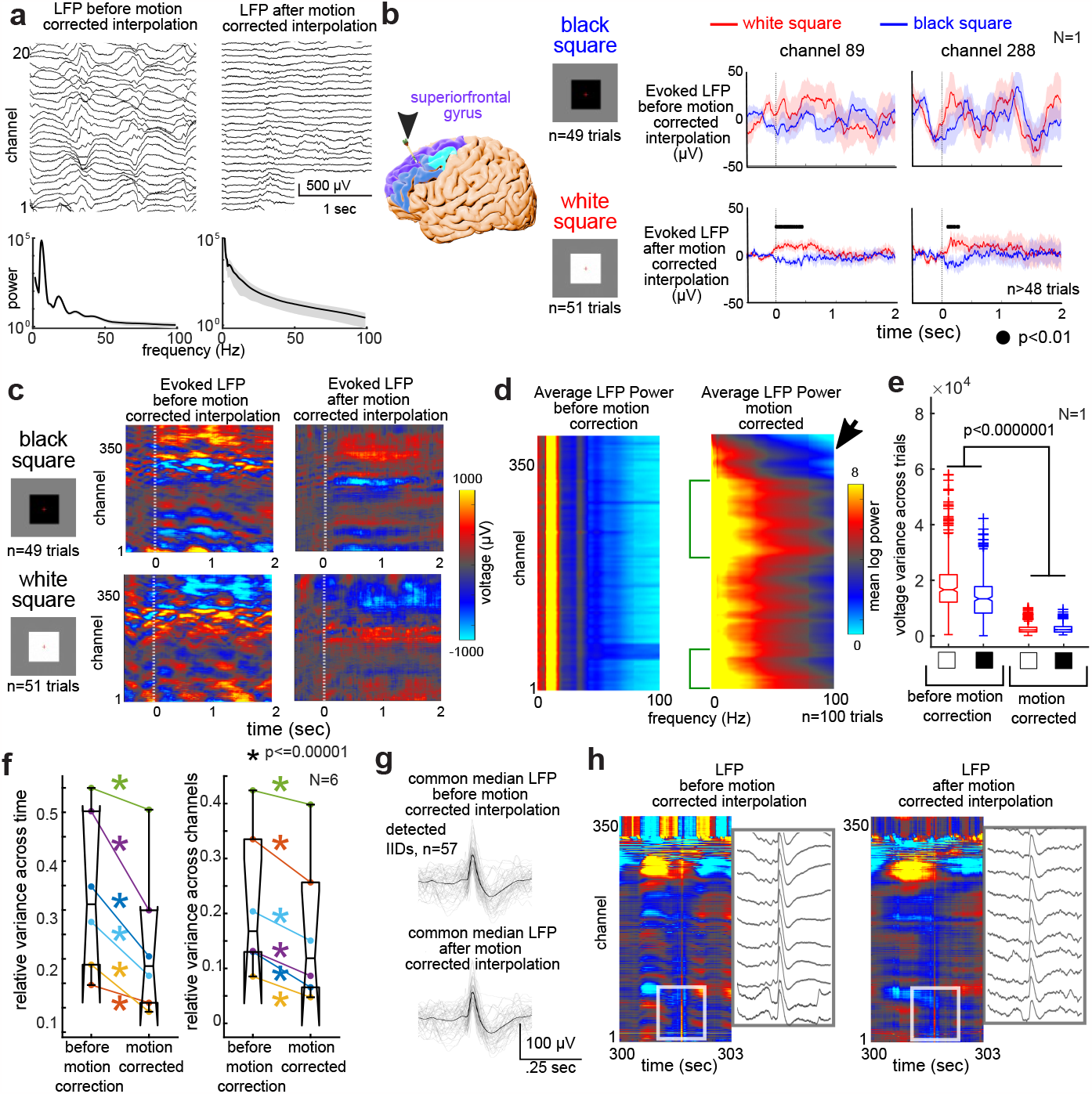
Correcting for motion in human local field potential data. **a** Top: Spontaneous LFP before and after motion-corrected interpolation and following Zapline-plus low-frequency peak removal. Bottom: Average power spectral curves before (left) and after (right) motion correction, averaged across channels. **b** In a recording in the superior frontal gyrus (also the dorsomedial prefrontal cortex), average visually evoked potentials can be observed in the LFP in a colormap to black versus white squares presented on a screen in front of the patient before and after motion-corrected inter-polation and Zapline-plus application. Brain regions in figure on the left: rostral middle frontal gyrus, blue; caudal middle frontal gyrus, cyan; superior frontal gyrus, purple. Arrow indicates location of the Neuropixels probe. **c** In the same recording in the superior frontal gyrus (also the dorsomedial prefrontal cortex), average visually evoked potentials across all channels can be observed in the LFP in a colormap before and after motion-corrected interpolation and Zapline-plus application. **d** Average log power spectral curves per channel (with the power represented as a color scale) before (left) and after (right) motion correction. Green brackets indicate ranges of channels with more power in the low and mid-frequencies across channel depths which are not evident before motion correction. Arrowhead indicates channels with lower power in the high frequencies in superficial channels. **e** Voltage variance across trials (first averaged across channels) before and after motion correction for the black and white visual stimuli. **f** Left: Relative variance averaged across 10 seconds of baseline activity per participant (different color dot lines and asterisks are different participants). Asterisks, *p* < 0.00001; pairwise Wilcoxon rank sum tests per participant. Right: Relative variance averaged across channels during baseline activity per participant (averaged across time, different color dot lines and asterisks are different participants). Asterisks, *p* < 0.00001; pairwise Wilcoxon rank sum tests per participant. **g** Common median LFP (across channels) of detected interictal epileptiform discharges (IIDs) before and after motion-correction interpolation, recorded in a patient with intractable epilepsy during an open craniotomy to remove epileptogenic tissue. **h** Spontaneous LFP per channel shown as a colormap and with a zoomed-in voltage trace of the same data for a detected IID before and after motion-corrected interpolation, showing that the IID survives the processing. The voltage and timing scale in **c** applies to the voltage traces here. The voltage colorbar in **c** applies to heatmaps here. Lower-indexed channels are deeper in the tissue.

This LFP-based motion correction was critical for identifying visual stimulus-induced evoked potentials in recordings in the dorsomedial prefrontal cortex (dmPFC, also the superiorfrontal gyrus). We presented a series of black and white squares to an awake participant undergoing DBS surgery and examined the LFP response in the dmPFC (Fig. 3.b; Supp. Fig. 8). As observed in other data sets^46^, the motion-corrected dmPFC LFP showed significantly different depth-specific average evoked potential responses to the visual stimuli on the per-channel level which differentiated between the black and white squares (*N* = 1; *p* < 0.01, Wilcoxon rank-sum test per time point, false discovery rate-corrected for multiple comparisons), whereas motion contamination had few to no image-onset induced differences between black and white square trials (*n* > 48 trials per condition (Fig. 3 and Supp. Fig. 8). Motion-correcting the LFP across channels further revealed depth-specific responses to the black versus white square stimuli that remained at the same depth throughout the averaged trial. Before motion correction, this voltage signal was highly variable vertically along the depth of the electrode (Fig. 3.c). As further validation that the motion correction could rescue physiologically relevant neural data which varies along the depth of the electrode in the cortex, we compared the power spectra across channels, averaged across trials. Before motion correction, we could not differentiate power spectral representations along the depth of the electrode. However, after motion correction, we found increased power in two different ranges of channels (green brackets in Fig. 3.d) and decreased high frequency power in the superficial layers (arrowhead; Fig. 3.d).

Even if the underlying neural response was present in the original LFP signal, the motion introduced not only large vertical movements but also significantly higher motion-induced voltage across trials with visual presentations (averaged across channels at 0.25 sec after image onset; *N* = 1; *p* < 0.000001, Kruskal-Wallis Test; Fig. 3.e). Taking baseline data without any stimuli across a total of 6 participants, we also found that motion correction along with Zapline-plus correction significantly decreased voltage variance on the per-participant level across time and across channels (*N* = 6; *p* < 0.000001, pairwise Wilcoxon rank-sum test per participant; Fig. 3.f).

To test whether these correction and interpolation steps either could rescue, or, alternately, remove neurally-induced LFP signals from contamination by motion and heartbeat artifacts, we next examined epileptiform interictal discharges (IIDs) before and after these preprocessing steps in Neuropixels recordings (Fig. 3; Supp. Fig. 9). We examined IID activity detected using automatic approaches and validated by an epileptologist (SSC) across the electrode depth in an open craniotomy case for the resection of anterior temporal lobe tissue in the treatment of epilepsy (*N* = 1; Fig. 3.g and Supp. Fig. 9; Paulk et al. ^15^). As the IIDs were large enough, we could detect them using the median of the LFP signal across channels both before and after motion-correction interpolation (Fig. 3.h). Importantly, in the raw traces as well as the IID-triggered average, we observed IID waveforms in the raw data which were not eliminated either after the motion-correction interpolation step or after Zapline-plus (Supp. Fig. 9). We found that the IIDs were larger on the probe contacts deeper in the tissue in this recording, which corresponded to the lower channel numbers in the figure, as has been observed in other cases of laminar recordings in the human cortex (*N* = 1, Pt03; Fig. 3 and Supp. Fig. 9; Fabo et al. ^47^). These results indicate that these processing steps can still result in an LFP signal that retains underlying neurophysiological signatures in the data set.

As DREDge’s motion tracking could be susceptible to signals which are widespread across the recording channels^48^, we next wanted to test whether general anesthesia-induced burst suppression activity could pose difficulties for DREDge, and whether the burst suppression signal could survive the interpolation step for motion correction of raw data (see Methods)^48;15^. On the contrary, following this motion-correction interpolation, the LFP still showed burst suppression voltage signatures which could be detected using automatic tools (Supp. Fig. 10; Westover et al. ^49^ ; Salami et al. ^50^). Indeed, we could detect bursts in the common median voltage traces at similar timings before and after motion-correction interpolation, with correlations between burst detections before and after motion-correction interpolation above 0.9 (Pt01, *r* = 0.93, *p* < 0.00001; Pt03, *r* = 0.95, *p* < 0.00001). Along with differentiating visual responses, the voltage variance, the power spectral differences, and IID detections, these results confirm that the processing steps to correct for the motion artifact detected by DREDge still allowed us to capture multi-channel dynamics related to neural processes which include differentiating sensory responses.

### Tracking long-range drift during probe insertion in non-human primates

A key advantage of the decentralized motion estimation framework is its ability to tolerate large nonstationarities in its input data, so that it does not require the same population of neurons to be present throughout an entire recording session. We thus hypothesized that DREDge would be able to track long-range drift surpassing the length of the probe, which would enable users to map the neural population recorded around the probe as it advances into the brain, in a manner similar to the previous tetrode study of Mechler et al. ^37^. To test this hypothesis, we implemented DREDge on long insertion datasets (*N* = 2, Fig. 4 and Supp. Fig. 11) recorded from rhesus macaque using Neuropixels 1.0-NHP probes^33^. The probe was inserted from the motor cortex targeting globus pallidus internus (GPi) in the basal ganglia using a commercial drive system (Fig. 4.a), with a target insertion depth of over 20 millimeters at a rate of 10μm/s (approximately 26 mm total estimated from drive motion, with an insertion speed of 10μm/s; recordings were cropped temporally to due to recording quality for input to DREDge).

**Figure 4.**
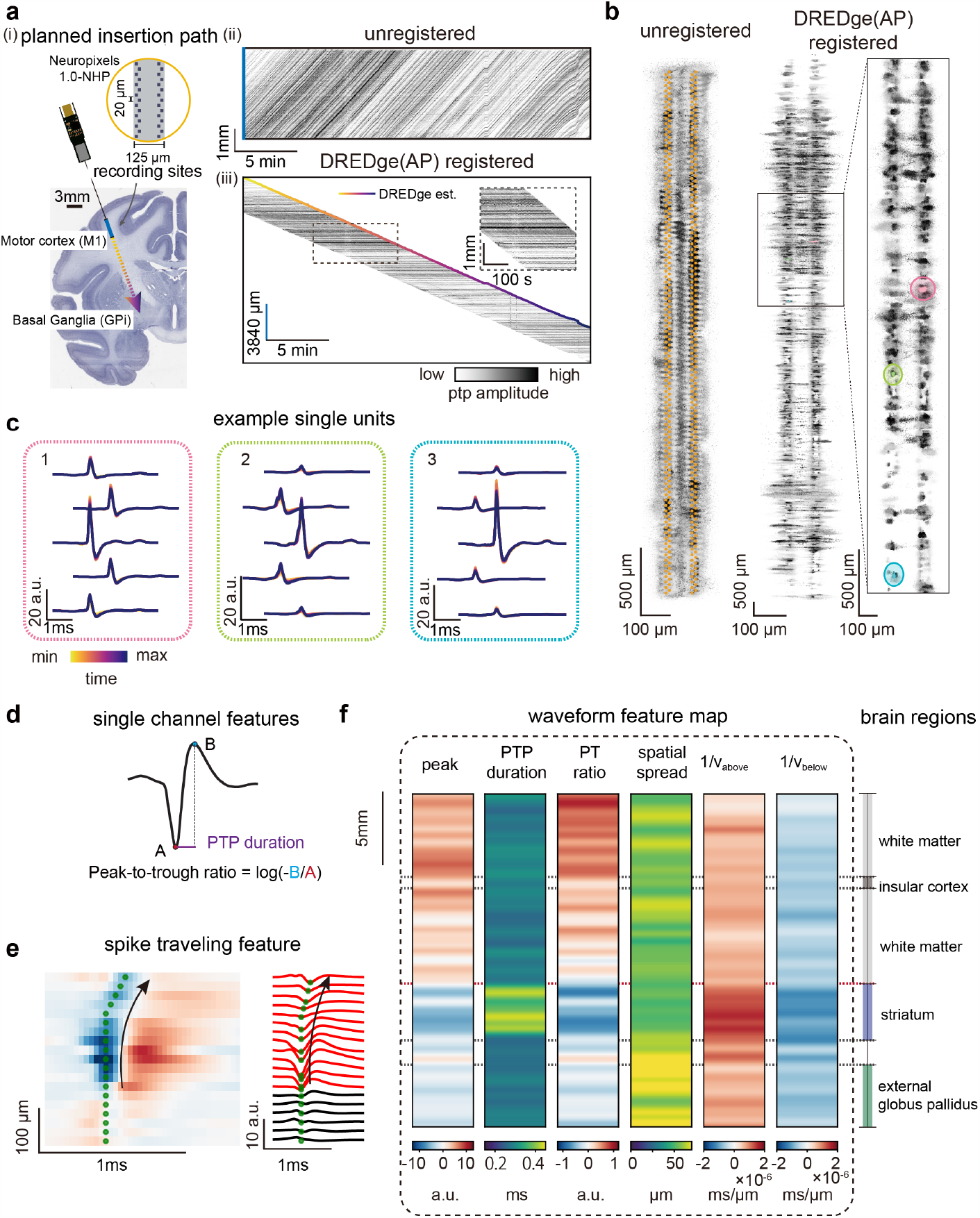
Monitoring long range drift during a deep probe insertion. **a** (i) The planned NP1.0-NHP probe insertion trajectory in the monkey brain (motor cortex to the internal globus pallidus). (ii) Spike raster before registration. (iii) Spike raster after registration, with DREDge estimated motion trace (scale bars: 3840μm vertical, 5min horizontal). **b** Localizations of detected spikes before (left) and after (right) drift correction according to the DREDge motion estimate. **c** Template waveforms for three example units estimated from time-binned (15s bins) spikes, clustered using location features stabilized using DREDge motion correction. Template waveforms are extracted on channel neighborhoods around the unit’s max amplitude channel in each time bin, and colored by time (color scale in **a**.i). The templates remain stable as the probe is inserted through its entire length. **d** Visual description of the three features extracted from spikes’ maximum amplitude channels and visualized in **f. e** Examples of traveling spike. The multiplicative inverses of the spike velocities below and above the channel with maximum peak-to-peak amplitude were shown as features in **f. f** Binned averaged spike features show consistent transitions across various depths, particularly near the putative striatal borders.

The large movement of the probe relative to the neuronal sources present during insertion was clearly visible in raster plots of spike depth positions over time (Fig. 4.a; Supp. Fig. 11.a). While KS’ template-based drift tracking failed in this case (Supp. Fig. 12), which we hypothesized was due to the difficulty of modeling several probe lengths’ of neuronal populations with a single template, DREDge was able to track motion across centimeters (Fig. 4.a).

To validate the motion estimate, we began by visualizing individual spikes’ vertical and horizontal locations in the plane of the probe, estimated using the point-source model of Boussard et al. ^36^ before and after motion correction (Fig. 4.b). While single unit clusters were completely obscured by the motion of the probe before motion correction (left), which is to be expected since each unit moved across the entire probe during the insertion, spike positions resolved into well-isolated clusters after registration (right). After manually isolating three clusters of spikes in the registered feature space, we separated their spike trains into 15-second temporal bins and computed average waveforms of the spikes in each bin. Plotting these waveforms on time-varying local channel neighborhoods extracted around their maximum amplitude channels revealed stable waveform shapes corresponding to single units as they traveled the length of the probe (Fig. 4.c).

In many experimental scenarios, the ability to accurately pinpoint the probe’s location within the target region’s anatomy is highly desirable. Experimenters identify the anatomical location of the probe during experiments by combining depth information from a drive system with observed changes in firing patterns along the insertion. However, this method can be subjective and prone to errors in depth estimates due to, e.g., tissue dimpling and deformation during insertion. Taking an alternative automated approach, we combined DREDge’s motion estimate with extracellular waveform features to determine the anatomical location of the probe. Waveform features were found to correlate with differences in cell type in previous studies^7;51^, so that collections of such features may also be informative in determining the brain region, thanks to the natural variability in cell type frequency across brain regions.

To correlate DREDge’s motion estimate with the approximately known anatomical trajectory of the probe in the NHP brain, which proceeded from motor cortex through white matter and striatum and finally to the internal globus pallidus (Fig. 4.d, right; Supp. Fig. 13), we collected waveform features from spikes observed across the insertion trajectory and studied their variation in relation to the motion-corrected spike depth. These features included the peak height, peak-to-peak duration, peak-to-trough ratio, spatial spread, and travel velocities of each spike (Fig. 4.d,e), and were computed after denoising each spike using the neural net denoiser of Lee et al. ^52^ ; more information on feature computation is included in Methods. Visualizing averages of these features as a function of motion-corrected spike depth revealed consistent variations which roughly aligned to expected anatomical boundaries along the insertion trajectory (Fig. 4.f). This experiment served both to validate DREDge’s long range motion tracking and to demonstrate the feasibility of simultaneous anatomical localization and electrophysiological feature mapping during probe insertion.

### Estimating motion in acute mouse recordings

Thus far, Neuropixels recordings have been made most frequently in mice. Since the mouse brain is much smaller than the primate brain, and since recordings made in mice may leverage experimental techniques such as head fixing which cannot be applied for instance in the human recordings discussed above, these recordings typically feature less extensive drift. Thus we were motivated to interrogate the extent to which DREDge could improve over Kilosort in mouse recordings. We began by comparing DREDge’s nonrigid spike-based motion estimation to that of KS on Neuropixels 1 and 2 datasets in which relatively small (∼50μm amplitude) vertical zig-zag probe was imposed via a micromanipulator (see Methods) where KS had previously been shown to perform well^6^. In these recordings, DREDge recapitulated the performance of KS (Fig. 5.a, i and ii). We compared DREDge to KS qualitatively in these datasets both by plotting the algorithms’ estimated nonrigid motion traces over a raster plot of spike positions over time (left), and by making raster plots of spike “registered positions” over time (i.e., positions offset inversely to the estimated motion; right, middle). In both NP1 and NP2, these algorithms’ estimated motion traces are similar and appear to qualitatively track the motion visible in the unregistered spike rasters, leading to well-stabilized registered raster plots.

**Figure 5.**
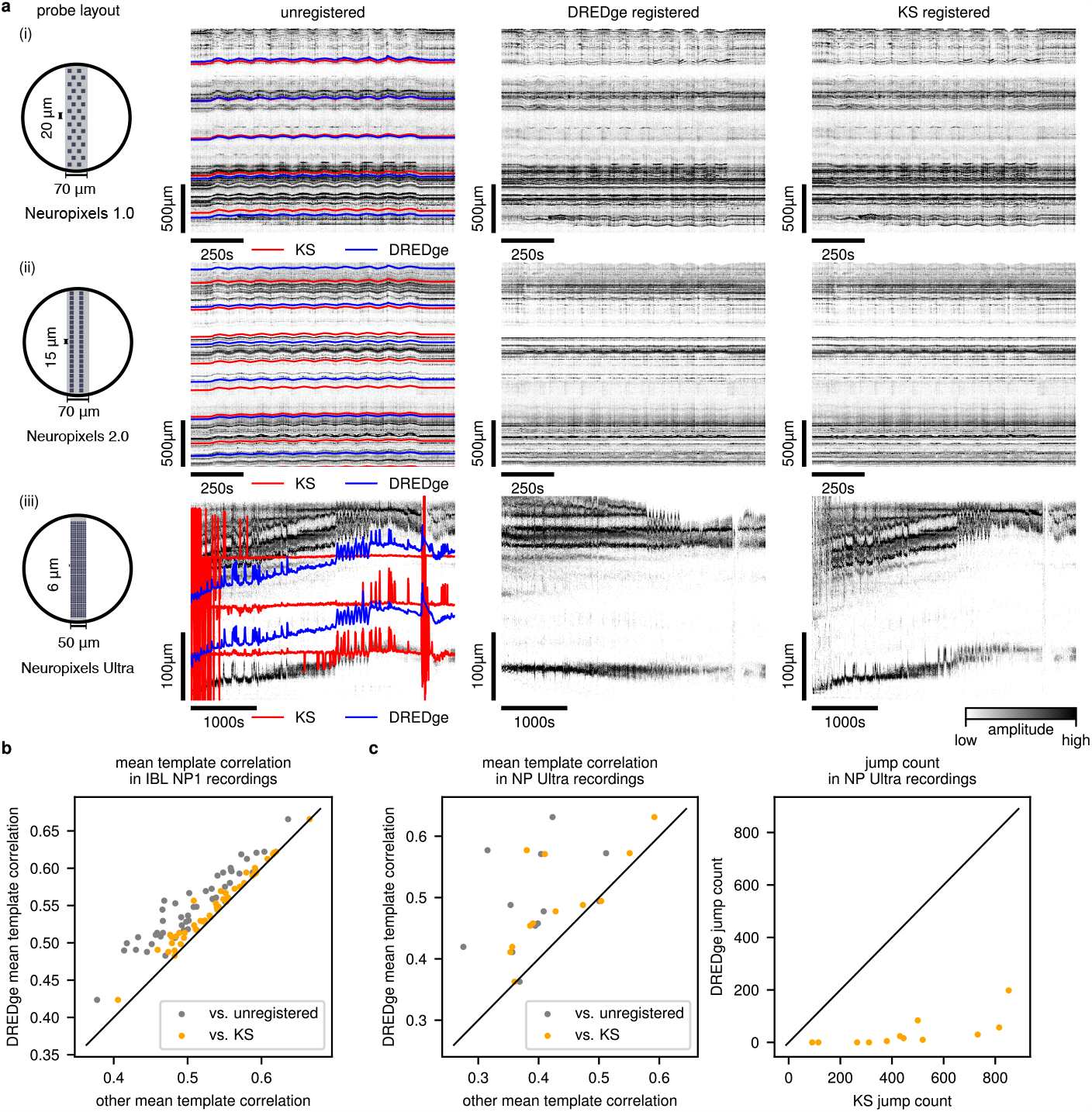
State of the art registration in acute mouse Neuropixels recordings. **a** Motion estimation from spikes detected in imposed motion datasets from Neuropixels 1, 2 (i, ii) and Ultra probes (iii)^6;34^. DREDge’s motion traces (left column, blue) and motion-corrected spike rasters (middle column) match the quality of Kilosort’s (left column, red; right column) on NP1 and NP2 data (i, ii). Unlike Kilosort (KS), DREDge also performs well when applied to the short and dense layout of the NP Ultra probe (iii). **b** In *n* = 47 datasets from the International Brain Lab’s repeated site experiment^32^, DREDge reliably outperforms KS on nonrigid spike-based registration according to a simple metric of registration quality (see Section 4.9). We computed this stability metric on unregistered, KS-registered, and DREDge-registered spike positions; here, the vertical position of a dot in the scatter shows the metric value after DREDge’s correction, and the horizontal position shows either the unregistered metric value (gray) or the value after KS’ correction (orange). **c**,**d** In *n* = 12 Neuropixels Ultra recordings with both natural and imposed zig-zag motion, DREDge reliably performs well relative to no correction and KS, leading to improvements in two metrics of stability. In **c**, we apply the metric study described in **b** to these NP Ultra datasets; colors and axes have the same meanings. In **d**, we plot the number of implausibly large jumps (motion estimation time bins with > 10μm/s drift; see also Section 4.9) which appear in DREDge’s and KS’ motion estimates; note that these large jumps are much more frequent in the KS output. Further visualizations of DREDge’s improvements in these NP Ultra recordings appear in Supp. Figs. 15 and 16.

Still, in qualitatively similar recordings with natural drift of a similar magnitude made by the International Brain Lab, DREDge reliably yielded improvements over KS. KS had already been employed by the International Brain Lab (IBL) in its motion estimation and spike sorting pipeline^20^. To perform a large scale comparison between DREDge and IBL’s application of KS on these datasets, we designed a metric of registration quality: taking inspiration from KS’ internal template heuristic, we computed the mean correlation of all time bins of each recording’s spike raster (before or after registration by KS or DREDge) with the raster’s temporal mean (see Section 4.9). Computing this metric on *n* = 47 IBL Neuropixels 1 recordings (Fig. 5.b) revealed that DREDge consistently improved the stability of the data when compared both to no registration and to KS (metric mean differences 0.04 and 0.01 respectively; two-sided paired *t* -test *p* < 10^*-*8^ in both cases). See Supp. Fig. 14 for illustrative examples. Although these improvements in correlation were modest, since the drift itself was modest, in no case did KS score higher on this metric than DREDge, a result which establishes DREDge as a state-of-the-art method in the case of acute mouse Neuropixels recordings.

Further, unlike KS, DREDge was able to track the same imposed zig-zag motion, plus additional probe motion, in recordings made with the Neuropixels Ultra (NP Ultra^34^) probe (Fig. 5.a, iii). This probe features a much smaller recording area than those of Neuropixels 1 or 2 (a vertical extent of 282μm when recording a dense channel neighborhood near the tip, versus 2880μm for NP2 and 3840μm for NP1 in their dense layouts), with the same number of recording channels in a much denser layout (6 columns of 48 electrodes with 6μm vertical and horizontal spacing). In this case, the raster plot of DREDge’s registered spike position revealed stably localized spikes from individual neuronal sources in a recording featuring both artificially imposed and other motion which were both substantial relative to the size of the recording area (Fig. 5.a, iii).

When applying DREDge and KS to *n* = 12 similar NP Ultra datasets, we repeatedly observed such improvements (Fig. 5.c,d). Since these datasets featured motion which was much larger relative to the recording area than in the IBL datasets, accurate motion estimation will have a larger impact on the recording. Indeed, as visualized in the left panel of Fig. 5.c, applying the template correlation metric analysis used above in the IBL Neuropixels study showed that DREDge led to larger improvements than we had observed in the IBL experiment. In the NP Ultra data, DREDge’s mean difference relative to no registration was 0.1 and relative to KS was 0.06; these values were both significantly different from 0 (two-sided paired *t* -test *p* < 0.01 in both cases). To validate the application of this metric as a measure of registration quality in these datasets, we also visualized the raw and motion-corrected spiking activity in 7 of these recordings in Supp. Fig. 15. We also plotted the frame-by-frame correlation to the template in all 12 recordings in Supp. Fig. 16. Further, we observed that DREDge tended to produce motion estimates with fewer physically implausible jump artifacts than KS on these datasets. We quantified this observation using a jump-counting metric which identified the number of frames in which each method estimated motion larger than a physical threshold of 10 μm/s (Fig. 5.d; see Section 4.9); these recordings should feature jumps of this magnitude only very rarely. DREDge’s motion estimate produced fewer such jump artifacts in all NP Ultra recordings studied, with 419 fewer jumps in each recording on average, a significant effect (paired *t* -test *p* < 0.0001).

We hypothesized that DREDge’s improvement in drift tracking over KS in these cases could relate to the NP Ultra probe’s smaller recorded depth relative to the range of drift relative to NP1 and NP2, which would lead to less agreement of individual frames with any global template like that which KS constructs. To test this hypothesis, we spatially subsetted the recording area in the NP1 and NP2 recordings of Fig. 5.a to fit inside the 282μm span of the NP Ultra probe; we found similar improvements in DREDge’s tracking relative to KS in this setting (Supp. Fig. 17). Together, these experiments increased our confidence in DREDge’s improvement in performance relative to KS in NP Ultra data and in general as the amplitude of motion increases relative to the length of the recorded area.

### Estimating motion in chronic mouse recordings

In chronic implantations, experimenters record in multiple sessions separated from each other by days or weeks from a single probe insertion. Within each session, chronic recordings can be more stable than acute recordings, especially when the probe is mounted directly on the skull rather rather than held in place externally; however, across sessions separated by days or weeks, changes arise in the firing pattern of the neuronal population as well as in single unit templates, complicating motion correction across sessions.

Since DREDge’s decentralized framework led to improved robustness to nonstationarities in firing patterns relative to KS in acute probe implantations, we hypothesized that DREDge would be well-suited to the task of registering recordings made across sessions recorded from individual chronic probe implantations. We studied DREDge’s performance on a collection of Neuropixels 1 recordings (*N* = 2, 31 and 57 recording sessions, 1.5 *±* 1.3 and 1.5 *±* 1.1 days between sessions; see also Methods) and four-shank Neuropixels 2 recordings (*N* = 2, 11 and 13 recording sessions, 13.1 *±* 6.0 and 13.5 *±* 11.2 days between selected sessions; see a timeline for one of the implantations in Fig. 6.a). The Neuropixels 2 recordings were made up of simultaneous recordings made on four shanks (jointly inserted and programmable recording arrays separated by 250μm) with 96 recorded channels per shank; we separated the recordings by shank, so that each session yielded four 96-channel recordings. We then took a simple and direct approach to chronic registration with both DREDge and KS, differing from previously-used KS-based pipelines^6^. Rather than co-registering consecutive pairs of recordings, we simply either combined spike position data collected across sessions or concatenated the raw binary data from different sessions and ran DREDge directly; DREDge’s modularity made both workflows straightforward (see Section 4.7 for information about running DREDge and KS on these data).

**Figure 6.**
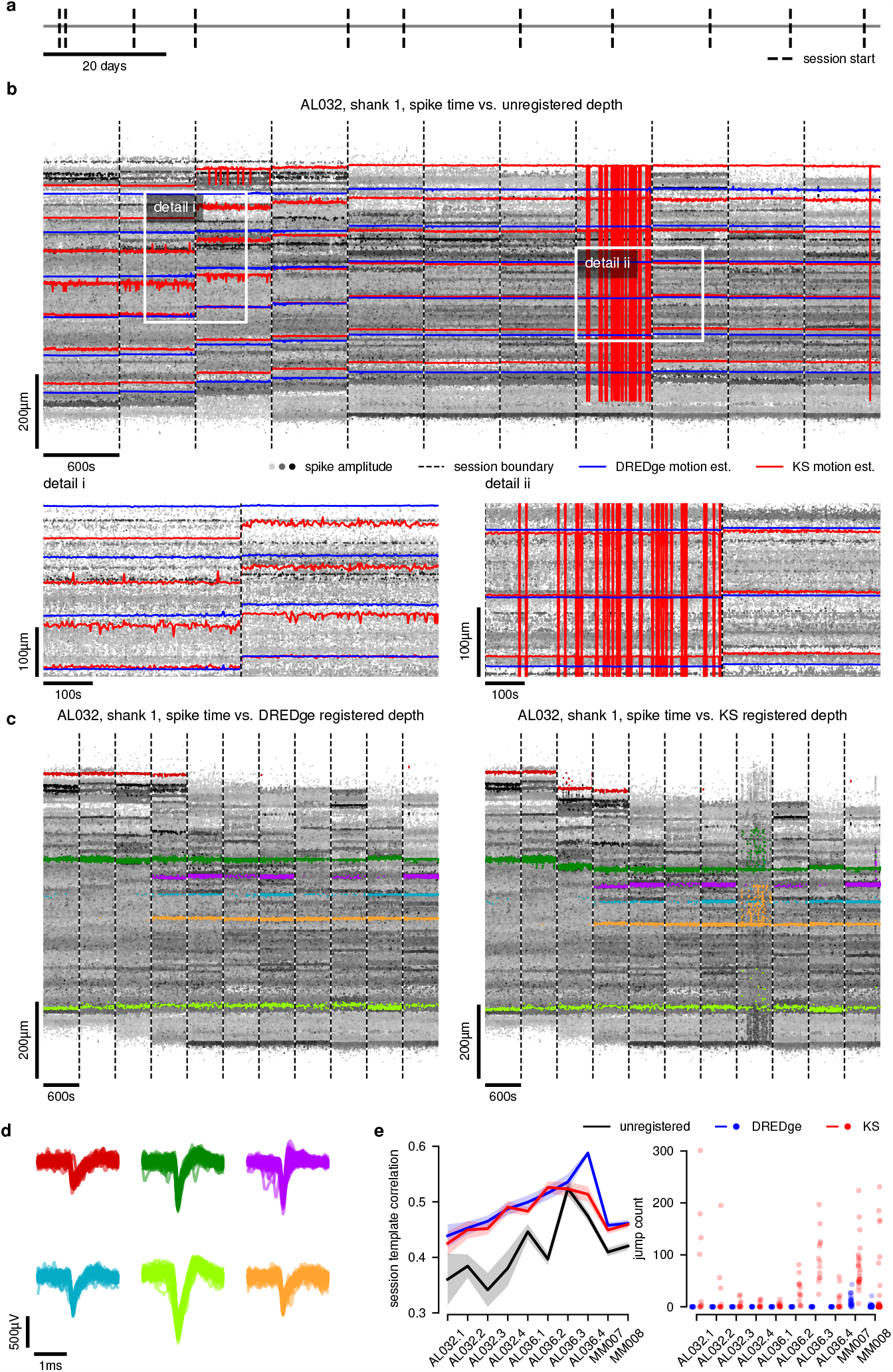
Tracking drift across weeks in chronic recordings. In **a**-**d**, we focused on 13 sessions from one shank of a chronic NP2.4 implantation (AL032, shank 1 of Steinmetz et al. ^6^). **a** Time-line of recordings and inter-session gaps, spanning 130 days. **b** Unregistered spike positions from 10-minute snips of each session plotted over time, with session boundaries indicated as vertical dashed lines; nonrigid DREDge and KS motion estimates appear as blue and red lines, respectively, centered on their nonrigid window center. Detail zooms highlight DREDge’s relative stability in comparison to KS. **c** Registered spike positions over time (DREDge, left; KS, right), with spike clusters which were manually isolated in the DREDge-registered spike positions shown in color; these putative units’ positions under KS’ motion trace are shown on the right. **d** Waveforms extracted on the detection channels for spikes in each cluster reveal well-stereotyped shapes, validating the motion estimate. **e** Comparisons to KS in two chronic NP2.4 implantations (separated into 4 shanks each) and two chronic NP1 implantations. DREDge is on par with or better than KS according to a simple metric of inter-session correlation (left; see Section 4.9; here we show the template correlation’s mean and standard error over sessions in each recording), and both methods improve on no registration. Further, according to a simple metric which counts non-physical jump artifacts (Section 4.9), the DREDge motion estimate is substantially more stable across this collection of chronic datasets (right); here, each dot shows the number of jumps in a single session, and each dataset contains many sessions.

In Fig. 6.a-d, we studied the drift tracking result in recordings from one shank of a Neuropixels 2 recording (AL032 shank 1^6^) in detail, applying DREDge and KS to 13 sessions made across 130 days with inter-session gaps of days or weeks. For an equal comparison, we ran KS on the concatenated binary representation, rather than following the pair-by-pair approach of previous work^6^. We first visualized DREDge’s and KS’ motion estimates over the unregistered spike raster (Fig. 6.b). In detail zooms, DREDge’s improvement in stability relative to KS became apparent, along with substantial differences in the motion estimation results, especially in the early upper portion of detail i.

Although DREDge offered a clear improvement in stability, it was not clear a priori whether the broad trend of motion it detected was more correct accurate than the trend of KS’ motion estimate. To check that this visual improvement corresponded to the real motion of the tissue, we isolated spikes from 6 putative single units by manually thresholding their amplitudes and motion-corrected positions (depth and horizontal position in the probe plane). These clusters are shown over the full set DREDge’s registered spike positions in the left panel of Fig. 6.c, and the corresponding plot for KS appears on the right, showing that spike positions which were stable under DREDge’s motion estimate corresponded to drifting or jumping trajectories under KS. Plots including the horizontal spike positions used to select spikes for these clusters appear in Supp. Fig. 18. We found that waveforms extracted on the maximum-amplitude channel at times corresponding to each of these spikes corresponded to well-stereotyped waveform shapes (Fig. 6.d), suggesting that the spikes did come from drifting single units, each of which were present across several sessions of the chronic recording; this provided evidence that, in this case, DREDge was tracking the probe trajectory more accurately than KS while also improving the stability of the motion estimate.

To test whether such improvements were repeatable, we computed metrics of DREDge’s performance against Kilosort’s on all 10 datasets. As in the previous section, we began by studying the mean template correlation metric (see Section 4.9), which correlates each time bin of the spike raster to the spike raster’s temporal mean and then considers the mean of those correlations. For this analysis, we visualized the spread of the mean template correlation session by session in Fig. 6.e (left); lines indicate the mean over sessions, and confidence bands show standard errors over sessions. Since the drift in these recordings is essentially nonexistent except in the first few sessions, this metric is not sensitive enough to differentiate DREDge and KS, including in cases like the one of Fig. 6.a-d discussed above where the metric values for DREDge and KS are very close; significant improvements in this metric only appear in cases such as AL036, shank 3 (shown in Supp. Fig. 19) which feature relatively large amounts of motion. However, visual inspection of other cases (Supp. Figs. 19 and 20) show that DREDge more accurately tracks what motion is present at the beginning of these recordings. Importantly, DREDge’s motion tracking maintains stability across this set of recordings, especially when compared to KS. We quantified stability using the jump-counting metric of the previous section (Fig. 6.e, right; see Section 4.9). DREDge’s motion estimate always led to fewer jumps, with differences in mean jump count per 10 or 3 minute session segment ranging from 5 to 167, with an average of 47 more jumps per session segment in KS’ motion trace; DREDge had no jumps in 74% of sessions versus KS’ 30%. Taken as a whole, these results introduce DREDge as a robust and simple drift-tracking algorithm for chronic MEA recordings.

## 3 Discussion

We have presented DREDge, a robust decentralized registration algorithm for both spiking and local field potential extracellular electrophysiology data recorded via dense multi-electrode probes. We applied DREDge to recordings made with several different high-density probe types (Neuropixels 1, 2, NHP, and Ultra; Neuroseeker), in multiple species (mouse, rat, macaque, human), and across recording types (AP, LFP, acute, chronic, intraoperative, during electrode insertion), and validated the efficacy of LFP- and AP-based motion tracking directly and in comparison to a previous automated approach (Kilosort 2.5) as well as manual tracking. The decentralized framework leads to natural robustness to changes in the neural populations present in the recording and their firing patterns, which enabled novel applications and improvements over current methods. First, in human intraoperative recordings which featured challenging high-amplitude and fast drift due to breathing and heartbeats along with long-term drift, DREDge’s LFP-based motion tracking enabled automated analyses of evoked local field potentials; this LFP-based tracking also enabled high temporal resolution motion correction of AP data, leading to improvements in single-unit spike sorting. Next, DREDge was able to track motion across many millimeters in recordings made during probe insertion through the relatively large brain of rhesus macaque, revealing variations in the electrophysiological properties of spikes across the depth of the insertion. In acute mouse recordings, DREDge outperformed existing approaches, especially when generalizing to new probe types. Finally, we were able to track motion across days and months in chronic recordings in mice.

DREDge’s code is fully open-source, and its modular implementation makes it easy to integrate into existing pipelines. It is already possible to integrate DREDge into current state-of-the-art spike sorting pipelines, such as Kilosort^27^, by using its motion estimate to drive motion-correction interpolation of the AP band as a preprocessing step via the SpikeInterface framework^19^. Further, DREDge is being integrated into new spike sorting pipelines which use a drift estimate to make their core routines drift-aware rather than relying on interpolation to correct for motion before sorting^35^. DREDge could also be integrated into other key steps in single-unit spike sorting, such as waveform-based quality metrics^53^ which are currently confounded by motion. DREDge’s motion estimation in chronic recordings could also be combined with existing approaches^54^ to enable stable tracking of single units over days and weeks.

DREDge’s core algorithm could also be extended to enable new workflows both in extracellular electrophysiology and in other domains, such as calcium imaging^26^ or cryogenic electron microscopy, where a related approach was already independently developed^55^. Finally, integrating DREDge as part of an online recording system could extend the simultaneous probe localization and electrophysiological feature mapping of our macaque insertion experiment to help experimenters target specific anatomy during recording on the fly, or even to increase the spatial precision of targeting for deep brain stimulation applications.

## Acknowledgements

We would like to thank Yangling Chou, Daniel Soper, Aaron Tripp, Fausto Minidio, Alex Zhang, Alexandra O’Donnell, and Michael Okun for their help in data collection. We would like to especially thank the patients for their willingness to participate in this research. We thank Matteo Carandini, Jennifer Colonell, Olivier Winter, Andrew Zimnik, and the International Brain Lab for helpful discussions and data coordination. We thank Alessio Buccino, Gaelle Chapuis, Margot Elmaleh, Samuel Garcia, Pierre Yger, and also the Simons Collaboration on the Global Brain Spike Sorting Working group for many useful discussions. This research was supported by the ECOR and K24-NS088568 (to SSC) and the Tiny Blue Dot Foundation (to SSC and ACP) and NIH grant U01NS121616 (to ZMW). This research was also supported by the Howard Hughes Medical Institute at Stanford University (to EMT). EMT is supported by the Grossman center and the Brain and Behavior Research Foundation. CW, JB, EV, and LP are funded by Simons Foundation 344 543023, NSF Neuronex Award DBI-1707398 and the Gatsby Charitable Foundation. EV is also supported by K99MH128772. The rat brain in vivo data has been recorded within the Hungarian Brain Research Program Grant (NAP2022-I-2/2022). DM is also supported by the OTKA Hungarian postdoctoral grant (PD143582). The views and conclusions contained in this document are those of the authors and do not represent the official policies, either expressed or implied, of the funding sources. The funders had no role in the study design, data collection, analysis, decision to publish, or preparation of the manuscript.

## 4 Methods

### 4.1 Preprocessing of action potential band data

For input into DREDge, raw electrophysiology data in the action potential band (300-6000Hz) passes through several steps, starting with quality control and filtering, then spike event detection and localization, and finally a rasterization step which leads to a binned spatiotemporal representation of spiking activity. These steps can be thought of as modular components that can be chosen according to user preference so that DREDge’s motion estimation step itself becomes another module in a bigger electrophysiology pipeline.

In the experiments conducted for this work, the initial filtering, detection, and localization steps were chosen to suit each data source. For the International Brain Lab (IBL) mouse recordings^32^, the IBL’s electrophysiology preprocessing pipeline^20^, including highpass filtering, analog-to-digital converter (ADC) offset correction, dead and noisy channel detection, and spatial highpass filtering, was reproduced using modular components available in the SpikeInterface framework^19^. Spike detections and localizations for input into DREDge were computed using the corresponding module from^35^, which relies on the point source model of Boussard et al. ^36^ to localize the spike events relative to the probe. Kilosort-based motion estimates were collected from IBL’s own runs of pyKilosort, a Python port of Kilosort 2.5^6^, and these motion estimates, in turn, used pyKilosort’s detected and localized spike events using raw data preprocessed as for DREDge^20^.

Once a collection of spike times, amplitudes, and localization features has been collected, DREDge processes these into a rasterized representation. Given spatial and temporal bin sizes *h*_d_, *h*_t_ (typically 1 micron and 1 second, respectively) leading to *D* bins along the length of the probe and *T* bins across time, all spikes landing in each spatiotemporal bin are collected. These are reduced into a *D* × *T* matrix, referred to here as the spike raster, by summing log(1 + *x*)-transformed spike amplitudes landing in each bin and transforming again with log(1 + *x*), followed by spatiotemporal smoothing (Gaussian filtering at 1μm and 1s scale). Here, the logarithmic transforms stabilize the representation to the heavy skewness present in the distributions of amplitudes and firing rates observed in natural data; similar transformations are performed by Kilosort 2.5 in its preprocessing before motion estimation. When constructing spike rasters for visualization (or for computing the template correlation metric of Section 4.9 and Figs. 5 and 6), the log transformations are not applied, since they lead to less interpretable units. Instead, the raster consists of the mean amplitudes of all spikes landing in each time and depth bin, filling in empty cells with zeros.

Kilosort 2.5 uses a similar preprocessing, constructing an image of suitably log transformed spike counts binned by their log transformed amplitudes and depths for each time bin, leading to a three-dimensional structure, in contrast to our two-dimensional raster; Kilosort’s full preprocessing is discussed in detail in the supplementary materials of Steinmetz et al. ^6^. A two-dimensional raster decreases the computational burden of pairwise cross-correlation and allows our method to share logic between the spike domain and LFPs, which are naturally easier to represent as images rather than three-dimensional structures.

### 4.2 Preprocessing of local field potential band data

In the local field potential (LFP) band, DREDge is able to operate directly on preprocessed electrophysiology traces, rather than on discrete events detected in this band. Relying on discrete events in the LFP band would be unreliable, since these are typically very sparse, and since fast motion can induce power in similar frequency bands as potential events of interest, confounding their detection in the presence of drift.

For input into DREDge, human LFP recordings^15^ were preprocessed according to the IBL’s electrophysiology preprocessing pipeline for LFP data^20^, including bandpass filtering, ADC offset correction, dead and noisy channel detection, and common referencing, and then downsampled to the target sampling rate for motion estimation (typically 250Hz; the effect of varying this rate was studied in Fig. 2.d-h). These steps were followed by a second spatial derivative along the probe’s vertical axis applied separately in each column, following averaging channels at the same depth. These latter steps sharpen the signal and represent it as a time-varying function over the depth domain, like the *D* × *T* spike raster in the AP band pipeline above, replacing the temporal bin size according to the preprocessed recording’s sampling frequency and the spatial bin size according to the vertical inter-channel spacing. This pipeline was implemented by means of open-source modules available in the SpikeInterface library^19^, allowing end users to substitute it with their own preprocessing.

### 4.3 Displacement and correlation matrices

Both the AP and LFP preprocessing pipelines above result in a time-varying signal represented over the long axis of the probe, which can be captured in a *D*×*T* matrix **R**, whose *D* rows represent depth bins and whose *T* columns represent time bins. Given such input, and in the case where a rigid displacement (i.e., a displacement that does not vary across depth) is being estimated, DREDge starts by calculating normalized cross-correlation^28^ vectors for each pair of time bins **R**_:*t*_ and **R**_:*t*′_, 1 ≤ *t, t*′ ≤ *T*. From each pair, the lag of the maximal cross-correlation and the maximal correlation value itself are used to populate *T* × *T* matrices **D** and **C**, so that **D**_*t*′_ is an estimate of the relative displacement between time bins *t* and *t*′ and **C**_*tt*′_ is the correlation of these time bins at this offset.

To extend to the nonrigid case, DREDge begins by dividing the depth domain into *B* user-configurable soft blocks with Gaussian profiles. For instance, *B* ≈ 10 evenly spaced Gaussian windows with bandwidth (standard deviation) of 500μm are well suited when estimating the nonrigid motions typically present in the IBL Neuropixels data of Fig. 5.b. Then, the normalized cross-correlations above are estimated for each of the *B* windows, substituting the formulas used to compute covariances and variances in the normalized cross-correlation with their weighted versions, where the soft windows are used as weights which decrease the contribution of depth bins far away from their centers to their displacement estimates. The results are then gathered as above into *B* × *T* × *T* arrays **D** and **C**, so that 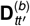 and 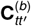 give the displacement and correlation between times *t, t*′ in the *b*th window, *b* = 1, …, *B*.

### 4.4 Robust decentralized registration

In the decentralized framework, the centralization problem (Varol et al. ^30^, equation 1) poses motion estimation as an optimization problem that models the estimated displacements between pairs of times as arising from differences of a true, unknown motion trace **P** across the corresponding time interval. In its basic form, the centralization problem is the minimization problem

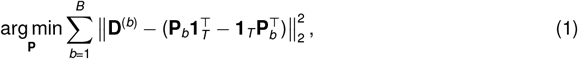

which seeks to find a motion trace **P**_*b*_ ∈ ℝ^*T*^ for each nonrigid block *b* = 1, …, *B* such that the pairwise differences of entries of **P**_*b*_ − **P**_*bt*′_, closely reconstruct the entries 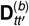 of the block’s *T* × *T* displacement matrix **D**^(*b*)^. In this way, **P** gathers the “decentralized” displacement estimates of **D** into a central motion estimate. When **D**^(*b*)^ is antisymmetric (naturally, the displacement **D**^(*b*)^ between times *t* and *t*′is the opposite of that between *t*′and *t*), the minimum of this basic version of the problem is attained by the row means of the displacement matrix,

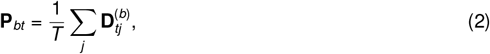

(see Section 7.1), but posing the problem in this decentralized framework enables several key modeling extensions.

Real-world data has several features which must be modeled in order to robustly estimate motion in both the AP and LFP bands. Multiple separate factors may make it impossible to estimate the relative displacement between two time bins by cross-correlation: these include nonstationarities in neural firing patterns, oscillations in the LFP band, changes in the neural population being recorded due to probe motion, and portions of recording with low signal. We implement three strategies to down-weight or exclude such pairs of time bins when estimating **P**. First, such effects often manifest in relatively low maximal correlations 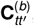, which can be accounted for during inference by ignoring pairs of time bins whose maximal correlation fails to exceed a threshold parameter *θ*_C_ and by weighting the rest of the terms by the corresponding correlations. Next, spatiotemporal regions of the recording with low activity can lead to spurious displacement and correlation estimates; it is beneficial to prevent such regions from affecting the rest of the motion estimate, which is achieved below via the spatiotemporal weights matrices **V**^(*b*)^. Finally, nonstationarities in firing patterns or LFP oscillations (possibly due to probe motion) can occur over long time periods. However, it is possible that time bins across these periods can have superficial similarities, leading to high correlations and spurious displacement estimates. The time horizon parameter *θ*_T_ below sets a limit on the time difference across which pairs of time bins are considered. Finally, the above measures can lead to spatiotemporal regions in which the motion estimate **P** is poorly determined. For instance, in noisy portions of a recording it is possible that all observations have been excluded due to low maximal correlations, leading to an ill-defined estimate of the motion in that region. In such cases, DREDge leverages a spatiotemporal smoothing term to make use of the information from neighboring temporal and spatial bins.

These spatiotemporal censoring, weighting, and smoothing operations are most simply introduced into the decentralized framework by restating it as a Bayesian inverse problem. To that end, we construct a probabilistic model which directly extends the centralization problem and in which **P** is considered a latent parameter to be inferred based on the observations **D**. To introduce the model, we start with the spatiotemporal smoothing prior. We let *R*(**P**) denote the negative log-prior, which penalizes large spatial and temporal derivatives:

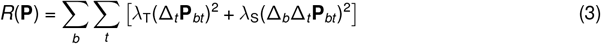

Here, _*t*_ and _*b*_ denote discrete temporal and spatial derivatives (i.e., _*t*_ **P**_*bt*_ = *P*_*b*(*t*+1)_ *-P*_*bt*_ when 1 < *t* < *T*); λ_T_, λ_S_ ≥0 control the relative importance of these terms and are set to 1 in all experiments above.

With this prior in place, we then model the observed displacements **D** as arising from the latent motion trace **P** with normally distributed errors:

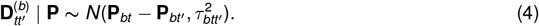

Here, we model the observed displacements 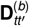 as conditionally independent given the latent displacement **P**. The variance 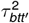, controls the weight of each observation and is given by

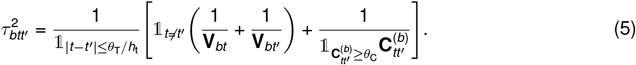

Here, **V**_*bt*_ is chosen to be either 0 or infinity depending on whether there is enough spiking activity in the *b*th window at time *t*, measured by computing the inner product of **R** with the *b*th window at that time and determining whether this value crosses a threshold parameter *θ*_V_. When |*t -t*′| > *θ*_T_ Or 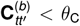, or in the case that **V**_*bt*_ or 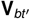 are 0, it is possible that 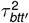, becomes infinite, which is equivalent to ignoring the observation 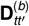. This observation model’s log likelihood is then a weighted version of equation (1).

In this framework, the centralization problem becomes the problem of maximum a posteriori inference of **P**:

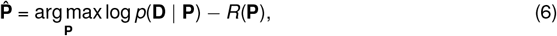

where *p*(**D** | **P**) and *R*(**P**) are the likelihood and negative log-prior above. The likelihood term *p*(**D** | **P**) factorizes over the *B* nonrigid windows, so that without the prior these *B* problems could be solved independently. However, the spatial smoothing of the prior links neighboring spatial windows, so that the *B* problems must be solved simultaneously. Fortunately, since *R*(**P**) only links neighboring nonrigid blocks, the Hessian matrix of the objective in equation (6) has block-tridiagonal structure when viewed as a *B* × *B* matrix of *T* × *T* blocks. Then, the inference problem as a whole reduces to a block-tridiagonal linear solve, which we carry out using a block version of the usual tridiagonal algorithm (Thomas’ algorithm). The time complexity of this operation scales linearly in the number of windows *B* and linearly in *T*, since the time horizon parameter *θ*_T_ above ensures that the blocks in the Hessian matrix are banded matrices with bandwidth less than *θ*_*T*_ ^56^; the dependence on the time horizon scales with 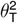.

### 4.5 Online motion tracking

When estimating motion in the LFP band at, for instance, 250Hz, *T* grows very rapidly, so that just a minute of recording would have *T* = 15000. Even with the linear complexity in *T* noted above, This rapid growth in the problem size leads to slow results when running the batch algorithm above in the LFP band. We mitigated these effects by choosing to estimate drift chunk by chunk in an ‘online’ fashion in these cases. In this online method, the preprocessed data **R** is processed in *C* chunks **R**^(*c*)^, *c* = 1, …, *C* of size at most *D* × *T*_0_. *T*_0_ = 2500 is our default and suggested choice for LFP applications, corresponding to 10s chunks of 250Hz-sampled preprocessed LFP data.

We initialize the algorithm by using the batch algorithm of the previous section to find the (possibly nonrigid) displacement estimate **P**^(1)^ in the first block. Then, given the previous chunk’s displacement estimate **P**^(*c*)^, we can find the current chunk’s displacement estimate **P**^(*c*+1)^ by solving a version of equation (6) where we condition on the previous chunk’s estimate **P**^(*c*)^:

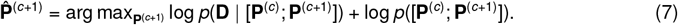

Here, [·; ·] is the operation which concatenates along the time axis (columns). Proceeding through the recording chunk by chunk, we can recover the full displacement estimate by concatenating those in each chunk. Since the sizes of the chunks’ sub-problems are bounded, this method will scale *linearly* in the total length of the recording.

### 4.6 Motion correction after DREDge

After estimating motion using DREDge, downstream applications will need to use this estimate to correct for motion artifacts in their data before further processing. In the LFP band, motion correction is carried out by interpolating the recording to infer its values at new, time-varying electrode positions chosen to move inversely to the motion estimate. Since LFP signals tend to be smooth in space, interpolation should not lead to much aliasing; however, features in spatial frequency bands which exceed the Nyquist rate corresponding to the probe’s electrode spacing may lead to distortion. Links to the Python and MATLAB code used to carry out this interpolation are below in Section 6.

In the AP band, a similar interpolation can be carried out using the SpikeInterface framework^19^. Alternatively, motion correction can be applied directly to the estimated positions of spikes extracted from uncorrected data^35^. In the motion-corrected or registered spike rasters which appear in many of the figures above and below, the corrected depth position of a spike at time *t* and depth *z* is computed by subtracting the estimated displacement at time *t* and depth *z* from *z*, where this displacement is estimated by bilinear interpolation between the displacement estimates at neighboring time and nonrigid depth bin centers.

### 4.7 Tracking drift in chronic recordings

When tracking drift in chronic recordings with DREDge, we followed two approaches. The first and simplest approach was to directly concatenate the raw data binary files and input them directly into the preprocessing and motion estimation pipelines described above; this approach was used for the chronic NP1 data. We also followed this approach in all cases when registering chronic recordings with Kilosort 2.5. For the chronic NP2 data, we ran preprocessing and extracted spike locations separately in each session. For input into DREDge, we then combined the detected spikes across sessions by offsetting the spike times in each session by the sum of the previous sessions’ durations. These combined spike events were then used to create the spike raster used for motion estimation with DREDge. These two approaches should yield similar results, and they were chosen in each case for methodological convenience. Apart from this difference, motion estimation with DREDge and KS were conducted in the same manner as the other analyses of this paper.

### 4.8 Tracking fast motion from spikes using clustering and splines

In cases where LFP signals are not available or where they do not contain features which are useful for motion tracking but spike data is plentiful, it may be necessary to correct for motion which is too fast to be modeled by DREDge’s or KS’ spike-based motion tracking, whose temporal resolution is limited to bins of length on the order of one or more seconds. In Fig. 2.b and Supp. Fig. 3, we introduced a method which uses spike data to correct for fast motion after initial coarse registration with DREDge. To do so, we used HDBSCAN^57^ to cluster high amplitude spikes by their registered location and amplitude features. Next, we obtained a time-series of spikes’ centered positions by subtracting the cluster’s mean registered depth from all spikes’ registered depths and then combining all of the spikes together into one point cloud. We then removed outliers (points more than 5 standard deviations from the mean in each cluster) and fit a smoothing spline to model the moving position of this point cloud as a function of time at sub-second temporal resolution. The number of knots of the fitted splines is equal to 2.5 times the number of seconds. These steps are detailed in Supp. Fig. 3. Note that this approach did not lead to improved registration accuracy in all cases; it is most useful in cases where there is rigid sub-second motion as well as sufficient density of high-amplitude spikes to allow for good spline estimates of the sub-second motion. In these cases (as in Supp. Fig. 3), this approach can significantly reduce within-cluster spike variability.

### 4.9 Spike registration quality metrics

To directly and quantitatively compare motion correction results before downstream processing such as spike sorting, we introduced two simple metrics. First, we developed a metric for registration quality of spiking data inspired by the template heuristic internally used by Kilosort’s motion estimation algorithm, which we referred to as the template correlation. To compute this metric for a given set of registered or unregistered spike locations, we first transform these into the two-dimensional spike raster described above in Section 4.1: spikes are binned into spatiotemporal time bins (1s and 1μm), and the mean amplitude of spikes in the bin is assigned to the corresponding position in the spike raster, leading to a *D* × *T* matrix with rows corresponding to the *D* depth bins and columns corresponding to the *T* time bins. Spatiotemporal bins which lie outside the extent of the probe after motion correction are masked. Next, we take the (masked) mean over time of this raster, leading to a template vector with *D* entries. Since areas outside the probe are ignored in this mean, it will not be contaminated by low- or no-activity bins. Finally, we compute Pearson’s *r* between each frame of the raster and this template, again ignoring masked spatiotemporal bins to avoid computing correlations of the template with empty space. This leads to *T* correlation values which can be used as a frame-wise measure of registration quality, as in Supp. Figs. 16, 19 and 20. Alternatively, the mean of these correlations can be presented for as a summary of registration quality for an entire recording, as shown in Fig. 5.b,c and Fig. 6.e.

In Fig. 6.e and Supp. Figs. 19 and 20, we also show a simple measure of the stability of motion estimation, which we refer to as the jump count. This metric directly captures the number of likely non-physical jumps in the estimated motion trace, by counting the number of registration time bins in which the motion estimate’s velocity exceeds 10μm/s relative to the previous bin.

#### Extracellular waveform feature extraction

In the analysis of Fig. 4, waveform features were computed from unsorted spikes detected by the initial detection step of Boussard et al. ^35^. We used the neural net described by Lee et al. ^52^ to denoise the detected waveforms on multiple electrodes. For single-channel features, we used the extracellular waveforms from the channels with the highest peak-to-peak (PTP) amplitude. Multi-channel waveforms were then extracted on the 40 channels closest to this maximum amplitude channel.

For single-channel waveforms, we computed three features: peak amplitude, peak-to-peak duration, and peak-to-trough ratio. Peak amplitude was the maximum point of the absolute waveform. Peak-to-trough duration was defined as the time difference between the maximum point and the minimum point of the waveform. The peak-to-trough ratio was defined as the logarithm of the absolute amplitude of the maximum point divided by the absolute amplitude of the minimum point.

For multi-channel waveforms, we computed three features: spatial spread of the spike across the probe, and the inverse of propagation velocity above and below the channel with maximum amplitude. The spatial spreads of the multi-channel waveforms were quantified using an amplitude-weighted sum of distances to the channel with maximum amplitude. If *a*_*i*_ denotes the PTP amplitude on channel *i* and *d*_*i*_ denotes the distance of this channel to the maximum amplitude channel, the spatial spread of each spike was computed as:

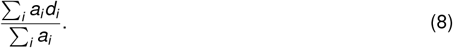

The inverse velocities were defined the same way as^7^ with the addition of a zero intercept constraints in the linear regression.

#### Brain anatomy estimation and alignment in the non-human primate recording

In the brain anatomy estimation of Fig. 4, since the resolution of MRI is poor for deep structures, the region boundaries for the monkey recording were identified by an expert from listening to the change in firing pattern during recording. The depth of the region boundaries corresponds to the actual recorded drive motion.

Due to the difficulty of penetrating the dura, the estimate of anatomical depth at the start of insertion is uncertain, so that the relative offset of DREDge’s motion estimate and the insertion drive’s measured distance is not known a priori. To align the anatomy with the computed feature map, we looked to match our observed electrophysiological features with the expert’s annotations in an easily identifiable landmark, namely the boundary between the white matter and the striatum. ‘Positive spiking’ units whose spikes contain large positive amplitudes before hyperpolarization are usually associated with dendrites and axons^58;59^. Thus, the white matter can be characterized by a high rate of such positive-going spikes, which is distinct from the striatum. We thus matched DREDge’s tracked depth with the drive motion and region boundaries by aligning the transition from positive to negative spikes to the boundary between the white matter and striatum. We used this offset as a reference to align the rest of the regions.

#### Setting parameters for DREDge and Kilosort

Due to the considerable variation in the types of drift observed across probe types, species, and importantly the methods used for probe mounting and implantation, it can be necessary to adjust the parameters of motion estimation algorithms (both DREDge and Kilosort 2.5). DREDge’s default parameters, discussed below, were determined in the large-scale International Brain Lab experiment whose results are shown in Fig. 5.b, and should apply well to recordings which are similar: i.e., stable Neuropixels recordings which feature mildly nonrigid motion on the order of 100μm. These recordings were made in head-fixed mice with an externally mounted probe, and thus feature some slight motion of the brain relative to the head; recordings made in different configurations, such as the human data of Fig. 2, where head-fixing and other brain stabilization methods cannot be used, or the chronic Neuropixels data of Fig. 6, where the probe is head-mounted, can present other drift scenarios that may require parameter adjustments. In this paper, we set parameters uniformly in all comparisons to Kilosort, in the sense that for each such experiment (i.e., set of recordings; for instance, the IBL experiment of Fig. 5.a, or the Neuropixels Ultra experiment of Fig. 5.a,c,d and Supp. Fig. 15), we used a fixed set of parameters for both DREDge and Kilosort across all datasets in each experiment; we did, however, tune the parameters of both DREDge and Kilosort for each experiment. In this section and Supp. Table 1, we present and discuss the set of parameters used for both DREDge and Kilosort in the experiments of this paper, and offer some suggestions about how DREDge’s parameters might generally be adjusted.

The most often adjusted parameters in DREDge are those which control the nonrigid windows. These windows have Gaussian profiles and divide the recording into a series of soft blocks, and they are parameterized by the distance between window centers (`win_step_um`, in μm) and the width of the windows (`win_scale_um`, in μm, which controls the standard deviations or bandwidths of the Gaussian bumps). In cases where the motion is known to be rigid (i.e., not to vary along the depth of the probe), the windowing can be turned off by setting the parameter `rigid=True`. Otherwise, these parameters may be tuned to match the amount of nonrigidity (i.e., the amount of variation in the motion along the depth of the probe) in the recording: more nonrigid motion will require more (i.e., more closely spaced or equivalently smaller `win_step_um`) windows. More nonrigidity may also require smaller window bandwidths (`win_scale_um`), since windows which are larger than the scale at which the motion varies as a function of depth may cover a varying motion profile. There is a tradeoff here, since setting `win_scale_um` to a small number will reduce the number of spikes or the amount of LFP signal falling into each window, which can reduce the stability and accuracy of the motion estimate in that window. Finally, the margin between the window centers and the edge of the probe is controlled by `win_margin_um`, in μm. To gain intuition about how to set these parameters and those discussed below, we encourage users to visualize the spike raster of Section 4.1; DREDge bundles functions for making these plots, which are in general very informative visualizations that can help users build intuition about not just the amount of drift in their recordings, but also the recordings’ quantity and amplitude of spikes and possible artifacts. Another parameter which can be helpful to adjust in some cases is the maximal offset used when computing cross-correlations (`max_disp_um`, in μm). This parameter limits the maximum spatial lag out to which cross-correlations are computed, and can be thought of as a “search radius” when comparing pairs of time bins. DREDge automatically sets this parameter to a quarter of the nonrigid spatial window size, but users can adjust this based on their own understanding of the amount of drift which is possible between time bins separated by less than the time horizon (*θ*_T_ above); such an understanding can be gained roughly by looking at spike raster visualizations. The time horizon itself was set to the fixed value of 1000s in all experiments here, except for the NHP insertion experiment of Fig. 4 where it was set to 100s; this setting allowed us to use our prior knowledge that the neuronal population was turning over rapidly during the insertion, avoiding spurious matches.

For completeness, we will briefly discuss other DREDge parameters of note which were not changed throughout this work. First, in all spike-based experiments shown here, the spatial and temporal bin sizes for spike raster computation in DREDge were set to 1μm and 1s, so that DREDge’s spike-based motion estimation always produced motion estimates with 1s temporal resolution; these basic parameters were chosen using simulation experiments (not shown). Second, the correlation threshold (*θ*_C_ above) was set to 0.1 in AP applications and 0.8 in LFP applications. Finally, the chunk size for online LFP registration was set to 10 seconds (or 2500 samples at 250Hz); since this chunk size is small, the time horizon parameter is not relevant in the LFP application.

Kilosort’s motion estimation algorithm was discussed in detail in the supplementary material of Steinmetz et al. ^6^ ; here we elaborate the discussion of certain parameters which were part of our tuning, based in part on our reading of Kilosort 2.5’s Matlab code (available at https://github.com/MouseLand/Kilosort/tree/v2.5/). Kilosort 2.5 exposes one parameter to control the registration, `nBlocks`, which controls the number of nonrigid blocks (rectangular windows rather than Gaussian profiles); the number of blocks used is 2·`nBlocks`*-*1 (see line 58 of align block2.m). When constructing its three-dimensional spike histogram, Kilosort uses a spatial bin size of 5μm and a temporal bin size controlled by the algorithm’s global batch size (expressed in samples), which leads to approximately 2.18*s* temporal bins in data sampled at 30kHz, although this will vary with the sampling rate; we did not find improvements in some exploratory experiments when tuning the spatial bin size and did not attempt to adjust the temporal bin size. The most important parameters which we adjusted in our experiments were Kilosort 2.5’s two search radius parameters (like our `max_disp_um` above), which are not exposed programmatically. The first of these, which we refer to as `nBinsReg1` and is in units of spatial bins, sets the maximal search radius of template cross-correlations during an initial rigid registration pass, before the recording has been divided into spatial blocks; this parameter is set to 15 bins (or 75μm) by default. The second, `nBinsReg2`, also in units of spatial bins, controls the search radius of template cross-correlations performed in the nonrigid pass, after the recording is divided into blocks, and is set to 5 bins or 25μm by default. Although these defaults are well suited to data with fairly small drift dominated by a rigid component, we found it essential to adjust them in recordings with larger drift or nonrigid drift whose overall amplitude was larger than 25μm. Our Kilosort 2.5 fork with modifications to expose these parameters is available at https://github.com/cwindolf/Kilosort/tree/modded-v2.5.

A table showing the parameters used in each experiment for both DREDge and Kilosort 2.5 appears in Supp. Table 1.

### 4.10 Datasets

#### Human brain activity in vivo data

Human brain activity was recorded in vivo in the course of clinically relevant neurosurgical intervention at both Massachusetts General Hospital (MGH) and the University of California San Francisco (UCSF) with most of the data and methods presented here included in previous publications^15;16^. In brief, in both data sets, all patients voluntarily participated after informed consent according to guidelines as monitored by the Massachusetts General Brigham (previously Partners) Institutional Review Board (IRB) Massachusetts General Hospital (MGH), and the UCSF Institutional Review Board. In all cases, participants were informed that participation in the experiment would not alter their clinical treatment in any way and that they could withdraw at any time without jeopardizing their clinical care. Participants were not compensated monetarily for participating. Recordings in the operating room were acquired with participants who were already scheduled for a craniotomy for concurrent clinical intraoperative neurophysiological monitoring or testing for mapping motor, language, and sensory regions and removal of tissue as a result of tumor or epilepsy or undergo intra-operative neurophysiology as part of their planned deep brain stimulator (DBS) placement^15;16;60;61;62^. Participants were either under general anesthesia or under monitored anesthesia care (awake or asleep) during the recordings according to clinical need (e.g. intraoperative stimulation mapping procedures or DBS surgeries). At MGH, participants also consented to the video recording of the surgical procedure as long as the video did not indicate the identity of the patient or staff. This video was used to confirm that the manual tracking could match the movement of the brain relative to the electrode. We performed tissue-level tracking of the video recordings to compare to the LFP-tracked motion tracking.

With both MGH and UCSF data collection sites, Neuropixels probes (NP v 1.0-S, IMEC) include an electrode shank (width: 70μm, length: 10 mm, thickness: 100μm) of 960 total sites laid out in a checkerboard pattern with contacts at 18 μm site to site distances (16 μm (column), 20 μm (row);^3^) with some probes with sharpened tips. The Neuropixels probes (NP v 1.0, version S, IMEC) were connected to a 3B2 IMEC headstage connected via a multiplexed cable to a PXIe acquisition module card (IMEC), installed into a PXIe Chassis (PXIe-1071 chassis, National Instruments)^15;16^. For the Neuropixels 1.0 probes as used in human studies, the linear dynamic range of the Neuropixels amplifier is 10 mVpp. This range is digitized using a 10 bits Analog to Digital conversion^63^.

At both collection sites, the Neuropixels probes were generally attached to a stable frame attached to the bed or frame around the skull or a stable arm with the probe being lowered to be inserted into the brain in sterile conditions. As such, this meant that, following exposure of the brain through a craniotomy, the brain tissue could move independently of the stably held Neuropixels probe. At UCSF, the Neuropixels probe was secured to the metal cap dovetail probe mount (IMEC, Leuven, Belgium). The probe mount was then attached to either an Elekta microdrive (Elekta, Stockholm, Sweden) or Narishige (Tokyo) micromanipulator (MM-3 or M-3333). Then, the manipulator/microdrive was either secured to the Mayfield skull clamp using a 3-joint mounting arm (Noga NF9038CA) and Nano clamp (Manfrotto 386BC-1, Cassola, Italy) assembly attached to the primary articulating arm and C-clamp of the Integra Brain Retractor System A2012 (Integra, Princeton, NJ)^16^. At MGH, the probe was either secured using SteriStrips (3M™ Steri-Strip™ Reinforced Adhesive Skin Closures) to a sterile syringe which was held by a 3-axis micromanipulator built for Utah array placement (Blackrock Neurotech, Salt Lake City, UT) or to cannulae placed in a NeuroFortis Neuro Omega manipulator (AlphaOmega Engineering, Nazareth, Israel) held by the ROSA ONE® Brain (Zimmer Biomet) arm^15^. At UCSF, probes, headstages, interface cables, Narishige micromanipulators, screwdrivers, and probe mount with metal cap dovetail were all separately sterilized according to standard protocols of ethylene oxide sterilization, while the Elekta device was sterilized using Sterrad. At MGH, the probe was sterilized with Ethylene Oxide (BioSeal) and used with the sterile Medtronic needle electrodes while the handling of the connections and recording equipment was wrapped in a sterile plastic bag and sealed using TegaDerm (3M) to keep the field sterile.

Ground and reference connections were kept separate in human brain recordings at both sites^15 16^. At MGH, recordings were referenced to sterile ground, and recording reference needle electrodes (Medtronic) connected (via safety connectors separately soldered to the separate ground and reference leads) were placed in nearby muscle tissue (often scalp) as deemed safe by the neurosurgical team. At UCSF, two 27G subdermal needle electrodes (Ambu, Columbia, MD) were placed in the skin were soldered separately to the probe flex-interconnect to serve as ground and reference using lead-free solder and two strands of twisted 36 AWG copper wire.

Data acquisition was performed using open-source acquisition software to record the neural data which include SpikeGLX (http://billkarsh.github.io/SpikeGLX/) and OpenEphys (Siegle et al. ^64^, https://open-ephys.org/gui). Since Neuropixels 1 probes enable 384 recording channels to address 960 electrodes across the probe shank, two different acquisition maps were used. At MGH, both one map (short column map) targeting the lower portion of the probe (the most distal channels) and a second map (‘long column’ map) recording two rows of contacts along the entire length of the electrode were used in different cases. The data collected at UCSF all included two rows of contacts along the entire length of the electrode.

For the sake of timing and correlating task activity with the neural activity, TTL triggers via a parallel port produced either during a task via MATLAB or custom code from a separate computer were sent to both the National Instruments and IMEC recording systems, via a parallel port system. This TTL output sent synchronization triggers via the SMA input to the IMEC PXIe acquisition module card to allow for added synchronizing triggers which were also recorded on an additional breakout analog and digital input/output board (BNC-2110, National Instruments) connected via a PXIe board (PXIe-6341 module, National Instruments)^15^.

For the simple visual task, stimuli were presented on an LCD computer monitor (58×30 cm, ASUS) placed in front of the participant and with the use of the Psychophysics toolbox^65^. The monitor distance from the subject was adjusted based on clinical considerations and the patient’s comfort and was placed 0.25 m away from the participant. The participant was asked to perform 100 trials of two different tasks, each distinguished by a certain visual stimulus. In the Square Task, each trial begins with the display of a red fixation cross for 0.5-4 sec on a grey background, before the appearance of a single black or white square with dimensions 5.5 cm × 5.5 cm (resulting in a visual display between 5.72º by 5.72º of the visual field) on a grey background, presented for 2-4 seconds with the duration jittered randomly. Each trial was composed of a fixation cross followed by either a black or white square and every trial was immediately after one another. The choice of black or white squares per trial was randomly selected from sequences of black or white designations pulled from a maximum-length sequence (m-sequence) distribution^66;67;68;69^. The participant was asked to fixate on the central red cross and count how many black or white squares were shown to improve engagement.

For a subset of the data (N=3), we used previously analyzed and manually tracked motion from the LFP to compare to the DREDge motion tracks^15^. Briefly, the steps involve extracting the LFP from the binary files into local field potential (LFP, ¡500 Hz filtered data, sampled at 2500 Hz) SpikeGLX using MATLAB and available preprocessing code. Focusing on non-noisy time ranges, we capture the displacement in the movement bands by importing the LFP voltage as an .stl file from MATLAB into Blender (https://www.blender.org/). Using the surface voltage and the Grease Pencil feature, we traced the shifting band of negatively deflecting LFP throughout the recording^15^. The motion traces were imported into MATLAB and compared with the LFP signal. This tracked motion information was upsampled to 2500 Hz to the LFP (interp1, ‘makima’).

The evoked potentials were averaged relative the image onset (2 seconds before and four seconds after image presentation). When analyzing spectral domains, we performed wavelet transforms to calculate the Morelet wavelet coefficient amplitude, the equivalent of power, to examine the amplitude of each frequency band from 0.5 to 200Hz. We subdivided the bands into delta (0-4Hz), theta (4-8Hz), beta (15-30Hz), gamma (30-55Hz), and high gamma (65-100Hz; Oostenveld et al. ^70^).

We tested comparisons across conditions with the Kruskal–Wallis test for non-equivalence of multiple medians to determine statistically separable groups or Wilcoxon rank sum test (two-sided) for pairwise comparisons between individual medians.

#### Mouse brain activity in vivo data

Extracellular recordings in mouse were obtained from multiple sources. For the quantitative comparison in Fig. 5.b, we relied on datasets recorded by laboratories participating in the International Brain Lab’s reproducible electrophysiology experiment^32^. The experiment recorded from 140 mice across 7 labs, and we processed recordings which passed the raw data quality control protocols described in that work (Table 1), which included target thresholds on the number of channels in the target region validated by histology, behavioral criteria, overall singleunit yield criteria, and limits on recording noise level. These SpikeGLX recordings were loaded via SpikeInterface and preprocessed according to the IBL’s standard preprocessing procedure^20^, including highpass filtering, demultiplexer phase shift correction, stripe artifact removal via a spatial highpass filter, and channel-wise standardization. This preprocessing pipeline was implemented via modules from SpikeInterface on all mouse recordings except for those from IBL, which were preprocessed using IBL’s own code available at https://github.com/int-brain-lab/ibl-neuropixel. These pipelines yielded similar results. These preprocessed recordings were then input into the initial spike detection, denoising, and localization pipeline of Boussard et al. ^35^ to extract point-source model localization features^36^ from that pipeline’s denoised and collision-cleaned waveforms. For the comparison to Kilosort, we used the IBL’s own runs of pyKilosort, a Python port of Kilosort 2.5, which were documented in more detail by IBL et al. ^20^ and which used the same preprocessing pipeline.

In Fig. 5.a, we included two acute recordings with imposed zig-zag motion from the work of Steinmetz et al. ^6^, described in more detail there. During these recordings (both included under dataset1 in the corresponding link in Data Availability below), one of which was performed using a Neuropixels 1.0 probe and the other with a Neuropixels 2.0 probe, 10 cycles of vertical triangle-wave drift with 50μm amplitude and 100s period were imposed via an electronic micromanipulator.

The chronic four-shank Neuropixels 2 recordings used in Fig. 6 and Supp. Fig. 19 were also previously presented by Steinmetz et al. ^6^. We studied two chronic implantations (AL032 and AL036) in detail, selecting 11 recordings separated by 13.1*±*6.0 days from AL032 and 13 recordings separated by 13.5 *±* 11.2 days from AL036.

The chronic Neuropixels 1 implantations recorded at UCLA were performed in compliance with the Institutional Animal Care and Use Committee. Two C57Bl6/J male mice (10-12 weeks of age) were used in experiments. Surgeries were performed under isofluorane anaesthesia (3% induced, 1.5-2% maintained). Headbar implantation and Neuropixels implantation were performed within the same surgery. First, the dorsal surface of the skull was cleared of skin and periosteum. A thin layer of cyanoacrylate (VetBond, World Precision Instruments) was applied to the edges of skull and allowed to dry. The skull was then scored with a scalpel to ensure optimal adhesion. After ensuring the skull was properly aligned within the stereotax, craniotomy locations were marked by making a small etch in the skull with a dental drill. A titanium headbar was then affixed to the back of the skull with a small amount of glue (Zap-a-gap). The headbar and skull were then covered with Metabond, taking care to avoid covering the marked craniotomy locations. After the Metabond was dry, the craniotomies for the probes and grounding screw were drilled. Once exposed, the brain was covered with Dura-Gel (Cambridge Neurotech). The implant was held using a custom plastic holder and positioned using Neurostar stereotax. After positioning the shanks at the surface of the brain, avoiding blood vessels, probes were inserted at slow speed (5 μm/s). Once the desired depth was reached, an additional layer of Kwik-Sil was applied over the craniotomy. The probe was then fixed to the skull with Metabond.

The Neuropixels Ultra data explored in Fig. 5 and Supp. Fig. 15 were reported in Ye et al. ^34^, and feature a very dense electrode layout, with 384 sites arranged in a 64 × 6 grid with 6μm vertical and horizontal channel spacing. Here, we focused on recordings with zig-zag motion imposed by a similar methodology as discussed above; more details are available in the reference.

#### Rat brain activity in vivo data

The rat recordings of Supp. Fig. 5 were made with the Neuroseeker probe, a 128-site high density probe, at 20 kHz with 16 bit resolution and with the rat under ketamine/xylazine anaesthesia^71;40^. These recordings are wideband (0.1-7500 Hz), so that LFP and AP were obtained by lowpass and highpass filtering.

#### Non-human primate brain activity in vivo data

The methods are described in detail elsewhere^33^, but, in brief, the Non-human primate recordings used the Neuropixels 1.0-NHP probe manufactured in two variants: 1) 45 mm long x 125 μm wide x 90 μm thick, featuring 4416 electrodes comprising 11.5 banks of 384 channels each; and 2) 25 mm long, 125 μm wide, and 60 μm thick, featuring 2496 electrodes comprising 6.5 banks of 384 channels with two aligned vertical columns. Probe tips were sharpened to a 25°angle using the Narishige EG-402 micropipette beveler. Neural recordings were referenced to either: 1) the large electrical reference point on the tip of the electrode, 2) an external electrical reference wire placed within the recording chamber, or 3) a stainless steel guide tube cannula. Electrical signals are digitized and recorded separately for the action potential (AP) band (10 bits, 30 kHz, 5.7 μV mean input-referred noise) and local field potential (LFP) band (10 bits, 2.5 kHz). Data collection was performed using SpikeGLX software. Recording sites are programmatically selectable with some constraints on site selection.

Multiple designs were used to allow for the lowering of the Neuropixels 1.0-NHP probes into the brain^33^. When using a non-penetrating guide tube, the dura was typically penetrated with a tungsten electrode prior to using a Neuropixels probe to create a small perforation in the dura to ease insertion. When inserting electrodes to deep targets (> 20mm), the alignment between the drive axis and the probe shank is essential for enabling safe insertion, as misalignment can cause the probe to break. For this application, we developed several approaches to maintain precise alignment of the probe and drive axis. The choice of appropriate insertion method depended on the mechanical constraints introduced by the recording chamber design, the depth of recording targets, the number of simultaneous probes required, and the choice of penetrating or non-penetrating guide tube. The interaction of these constraints and a more thorough discussion of insertion approaches is provided on the Neuropixels users wiki^33^. Open-source designs for mechanical mounting components for Neuropixels-1.0-NHP to drives from Narishige, NAN, and other systems are available in a public repository: https://github.com/etrautmann/Neuropixels-NHP-hardware.

The recording used in Supp. Fig. 3 was made in an anesthetized paralyzed preparation, described in detail previously^72^. We induced anesthesia with an intramuscular injection of ketamine HCI (10 mg/kg) and maintained the animal with isoflurane anesthesia during catheterization of saphenous veins and endotracheal intubation. Throughout the experiment, we maintained anesthesia with an infusion of 6 and 15 μg/kg/h sufentanil citrate and neuromuscular blockade with 0.1 mg/kg/h vecuronium bromide to limit eye movements. We opened a craniotomy and durotomy to insert a Neuropixels array^3^ or 2-shank 128-channel silicon laminar arrays from the NeuroNex Technology Hub^73^. The sites were sealed with agar, and petroleum jelly was routinely applied to prevent the agar from drying and maintain the cortex’s health. We generated and controlled stimuli with an Apple Mac Pro computer. We presented stimuli on a CRT monitor (HP1190) running at a resolution of 1280 × 960 pixels (64 pixels per degree) and 120 Hz. Most stimuli were binary or ternary noise patterns presented at a rate of 40 Hz.

## 5 Data availability

Human data is available for download at Dryad (https://doi.org/10.5061/dryad.d2547d840) and DANDI (https://dandiarchive.org/dandiset/000397) from Massachusetts General Hospital^15^ and at Dryad (https://doi.org/10.7272/Q6ST7N3B) from the University of California San Francisco^16^.

International Brain Lab data for the reproducible electrophysiology experiment is publicly available and can be downloaded by following the instructions at https://int-brain-lab.github.io/iblenv/notebooks_external/data_release_repro_ephys.html using the tag 2022_Q2_IBL_et_al_RepeatedSite. The NP1 and NP2 imposed motion datasets here (dataset1) can be downloaded at Figshare https://figshare.com/articles/dataset/_Imposed_motion_datasets_from_Steinmetz_et_al_Science_2021/14024495?file=26476589.

## 6 Code availability

DREDge is available to run on AP data via the SpikeInterface library, and on both AP and LFP data by open-source Python code hosted at the GitHub repository https://github.com/evarol/dredge/. DREDge is implemented in Python, and it relies on PyTorch’s convolution routines to implement GPU-accelerated normalized cross-correlations^74^, on SciPy for its bundled linear system solvers and interpolation routines^75^, and on SpikeInterface^19^ for its electrophysiology data readers and preprocessing routines, some of which were implemented as part of this work.

Code for running Kilosort 2.5 with an extended set of adjustable parameters is available at https://github.com/cwindolf/Kilosort/tree/modded-v2.5.

Code for the analyses of human data described in this paper has been made available at https://github.com/Center-For-Neurotechnology/HumanNeuropixelsPipeline (currently without a license), which includes links to other useful repositories not maintained by authors of this paper, with the exceptions of https://github.com/evarol/dredge (available under the MIT license) and https://github.com/williamunoz/InterpolationAfterDREDge) (available under the MIT license). Local field potential motion corrected interpolation required the removal of low-frequency peaks in the signal, a step utilizing Zapline-plus (https://github.com/MariusKlug/zapline-plus). For all the Neuropixels data, open source acquisition software was used to acquire the neural data which include SpikeGLX Release v20201103-phase30 (http://billkarsh.github.io/SpikeGLX/) and OpenEphys (https://open-ephys.org/gui). Single unit sorting was performed using Kilosort (https://github.com/MouseLand/Kilosort) as well as Phy2 (https://github.com/cortex-lab/phy). Custom Matlab (version R2021a) and Python code in combination with open source code from the Field-trip toolbox (http://www.fieldtriptoolbox.org/, Oostenveld et al. ^70^) was used for the majority of the analyses. Some code involving manual alignment is available on GitHub (https://github.com/Center-For-Neurotechnology/CorticalNeuropixelProcessingPipeline). The burst suppression ratio (BSR) was computed using an automated method (https://github.com/drasros/bs_detector_icueeg). Psychtoolbox-3 (http://psychtoolbox.org/) with io64 parallel port drivers and MATLAB functions were used to drive TTL trigger pulses for alignment as well as run the visual task.

## References

[1] Urs Frey, Jan Sedivy, Flavio Heer, Rene Pedron, Marco Ballini, Jan Mueller, Douglas Bakkum, Sadik Hafizovic, Francesca D Faraci, Frauke Greve, Kay-Uwe Kirstein, and Andreas Hierle-mann. Switch-Matrix-Based High-Density microelectrode array in CMOS technology. IEEE J. Solid-State Circuits, 45(2):467–482, February 2010.

[2] Bogdan C Raducanu, Refet F Yazicioglu, Carolina M Lopez, Marco Ballini, Jan Putzeys, Shi-wei Wang, Alexandru Andrei, Veronique Rochus, Marleen Welkenhuysen, Nick van Helleputte, Silke Musa, Robert Puers, Fabian Kloosterman, Chris van Hoof, Richárd Fiáth, István Ulbert, and Srinjoy Mitra. Time multiplexed active neural probe with 1356 parallel recording sites. Sensors, 17(10), October 2017.

[3] James J Jun, Nicholas A Steinmetz, Joshua H Siegle, Daniel J Denman, Marius Bauza, Brian Barbarits, Albert K Lee, Costas A Anastassiou, Alexandru Andrei, Çağatay Aydın, et al. Fully integrated silicon probes for high-density recording of neural activity. Nature, 551(7679):232–236, 2017.

[4] Richárd Fiáth, Bogdan Cristian Raducanu, Silke Musa, Alexandru Andrei, Carolina Mora Lopez, Marleen Welkenhuysen, Patrick Ruther, Arno Aarts, and István Ulbert. Fine-scale mapping of cortical laminar activity during sleep slow oscillations using high-density linear silicon probes. J. Neurosci. Methods, 316:58–70, March 2019.

[5] Gian Nicola Angotzi, Fabio Boi, Aziliz Lecomte, Ermanno Miele, Mario Malerba, Stefano Zucca, Antonino Casile, and Luca Berdondini. SiNAPS: An implantable active pixel sensor CMOS-probe for simultaneous large-scale neural recordings. Biosens. Bioelectron., 126:355–364, February 2019.

[6] Nicholas A. Steinmetz, Cagatay Aydin, Anna Lebedeva, Michael Okun, Marius Pachitariu, Marius Bauza, Maxime Beau, Jai Bhagat, Claudia Böhm, Martijn Broux, Susu Chen, Jennifer Colonell, Richard J. Gardner, Bill Karsh, Dimitar Kostadinov, Carolina Mora-Lopez, Junchol Park, Jan Putzeys, Britton Sauerbrei, Rik J. J. van Daal, Abraham Z. Vollan, Marleen Welkenhuysen, Zhiwen Ye, Joshua Dudman, Barundeb Dutta, Adam W. Hantman, Kenneth D. Harris, Albert K. Lee, Edvard I. Moser, John O’Keefe, Alfonso Renart, Karel Svoboda, Michael Häusser, Sebastian Haesler, Matteo Carandini, and Timothy D. Harris. Neuropixels 2.0: A miniaturized high-density probe for stable, long-term brain recordings. bioRxiv, 2020. doi: 10.1101/2020.10.27.358291. URL https://www.biorxiv.org/content/early/2020/10/28/2020.10.27.358291.

[7] Xiaoxuan Jia, Joshua H Siegle, Corbett Bennett, Samuel D Gale, Daniel J Denman, Christof Koch, and Shawn R Olsen. High-density extracellular probes reveal dendritic backpropagation and facilitate neuron classification. Journal of neurophysiology, 121(5):1831–1847, 2019.

[8] Ethan B Trepka, Shude Zhu, Ruobing Xia, Xiaomo Chen, and Tirin Moore. Functional interactions among neurons within single columns of macaque V1. Elife, 11, November 2022.

[9] Janis Karan Hesse and Doris Y Tsao. A new no-report paradigm reveals that face cells encode both consciously perceived and suppressed stimuli. Elife, 9, November 2020.

[10] Eric M Trautmann, Sergey D Stavisky, Subhaneil Lahiri, Katherine C Ames, Matthew T Kaufman, Daniel J O’Shea, Saurabh Vyas, Xulu Sun, Stephen I Ryu, Surya Ganguli, and Krishna V Shenoy. Accurate estimation of neural population dynamics without spike sorting. Neuron, 103 (2):292–308.e4, July 2019.

[11] Hidehiko K Inagaki, Susu Chen, Margreet C Ridder, Pankaj Sah, Nuo Li, Zidan Yang, Hana Hasanbegovic, Zhenyu Gao, Charles R Gerfen, and Karel Svoboda. A midbrain-thalamus-cortex circuit reorganizes cortical dynamics to initiate movement. Cell, 185(6):1065–1081.e23, March 2022.

[12] Selmaan N Chettih, Emily L Mackevicius, Stephanie Hale, and Dmitriy Aronov. Barcoding of episodic memories in the hippocampus of a food-caching bird. bioRxiv, July 2023.

[13] Nicholas A Steinmetz, Peter Zatka-Haas, Matteo Carandini, and Kenneth D Harris. Distributed coding of choice, action and engagement across the mouse brain. Nature, 576(7786):266–273, December 2019.

[14] Aditi Pophale, Kazumichi Shimizu, Tomoyuki Mano, Teresa L Iglesias, Kerry Martin, Makoto Hiroi, Keishu Asada, Paulette García Andaluz, Thi Thu Van Dinh, Leenoy Meshulam, and Sam Reiter. Wake-like skin patterning and neural activity during octopus sleep. Nature, 619(7968): 129–134, July 2023.

[15] Angelique C. Paulk, Yoav Kfir, Arjun R. Khanna, Martina L. Mustroph, Eric M. Trautmann, Dan J. Soper, Sergey D. Stavisky, Marleen Welkenhuysen, Barundeb Dutta, Krishna V. Shenoy, Leigh R. Hochberg, R. Mark Richardson, Ziv M. Williams, and Sydney S. Cash. Large-scale neural recordings with single neuron resolution using neuropixels probes in human cortex. Nature Neuroscience, 25(2):252–263, January 2022. doi: 10.1038/s41593-021-00997-0. URL 10.1038/s41593-021-00997-0.

[16] Jason E Chung, Kristin K Sellers, Matthew K Leonard, Laura Gwilliams, Duo Xu, Maximilian E Dougherty, Viktor Kharazia, Sean L Metzger, Marleen Welkenhuysen, Barundeb Dutta, et al. High-density single-unit human cortical recordings using the Neuropixels probe. Neuron, 2022.

[17] Nicholas A Steinmetz, Christof Koch, Kenneth D Harris, and Matteo Carandini. Challenges and opportunities for large-scale electrophysiology with neuropixels probes. Curr. Opin. Neurobiol., 50:92–100, June 2018.

[18] Marius Pachitariu, Nicholas Steinmetz, Shabnam Kadir, Matteo Carandini, and Kenneth D Harris. Kilosort: realtime spike-sorting for extracellular electrophysiology with hundreds of channels. bioRxiv, page 061481, June 2016.

[19] Alessio P Buccino, Cole L Hurwitz, Samuel Garcia, Jeremy Magland, Joshua H Siegle, Roger Hurwitz, and Matthias H Hennig. Spikeinterface, a unified framework for spike sorting. eLife, 9:e61834, nov 2020. ISSN 2050-084X. doi: 10.7554/eLife.61834. URL 10.7554/eLife.61834.

[20] IBL, Kush Banga, Julien Boussard, Gaë lle A Chapuis, Mayo Faulkner, Kenneth D Harris, Julia M Huntenberg, Cole Hurwitz, Hyun Dong Lee, Liam Paninski, Cyrille Rossant, Noam Roth, Nicholas A Steinmetz, Charlie Windolf, and Olivier Winter. Spike sorting pipeline for the International Brain Laboratory. Technical report, International Brain Laboratory, 5 2022. URL https://figshare.com/articles/online_resource/Spike_sorting_pipeline_for_the_International_Brain_Laboratory/19705522.

[21] John Ashburner. A fast diffeomorphic image registration algorithm. Neuroimage, 38(1):95–113, 2007.

[22] Aristeidis Sotiras, Christos Davatzikos, and Nikos Paragios. Deformable medical image registration: A survey. IEEE transactions on medical imaging, 32(7):1153–1190, 2013.

[23] Ignacio Arganda-Carreras, Carlos O. S. Sorzano, Roberto Marabini, José María Carazo, Carlos Ortiz-de Solorzano, and Jan Kybic. Consistent and elastic registration of histological sections using vector-spline regularization. In Proceedings of the Second ECCV International Conference on Computer Vision Approaches to Medical Image Analysis, CVAMIA’06, page 85–95, Berlin, Heidelberg, 2006. Springer-Verlag. ISBN 3540462570. doi: 10.1007/118897628. URL 10.1007/11889762_8.

[24] B Avants, C Epstein, M Grossman, and J Gee. Symmetric diffeomorphic image registration with cross-correlation: Evaluating automated labeling of elderly and neurodegenerative brain. Medical Image Analysis, 12(1):26–41, February 2008. doi: 10.1016/j.media.2007.06.004. URL 10.1016/j.media.2007.06.004.

[25] Alexander Dubbs, James Guevara, and Rafael Yuste. moco: Fast motion correction for calcium imaging. Frontiers in neuroinformatics, 10:6, 2016.

[26] Eftychios A. Pnevmatikakis and Andrea Giovannucci. Normcorre: An online algorithm for piece-wise rigid motion correction of calcium imaging data. Journal of Neuroscience Methods, 291: 83–94, 2017. ISSN 0165-0270. doi: 10.1016/j.jneumeth.2017.07.031. URL https://www.sciencedirect.com/science/article/pii/S0165027017302753.

[27] Marius Pachitariu, Shashwat Sridhar, and Carsen Stringer. Solving the spike sorting problem with Kilosort. bioRxiv, 2023. doi: 10.1101/2023.01.07.523036. URL https://www.biorxiv.org/content/early/2023/01/07/2023.01.07.523036.

[28] JP Lewis. Fast Normalized Cross-Correlation. Technical report, Industrial Light & Magic, 1995. URL http://scribblethink.org/Work/nvisionInterface/nip.pdf.

[29] Samuel Garcia, Charlie Windolf, Julien Boussard, Benjamin Dichter, Alessio P Buccino, and Pierre Yger. A modular approach to handle in-vivo drift correction for high-density extracellular recordings. bioRxiv, page 2023.06.29.546882, June 2023.

[30] Erdem Varol, Julien Boussard, Nishchal Dethe, Olivier Winter, Anne Urai, The International Brain Laboratory, Anne Churchland, Nick Steinmetz, and Liam Paninski. Decentralized motion inference and registration of neuropixel data. In ICASSP 2021 - 2021 IEEE International Conference on Acoustics, Speech and Signal Processing (ICASSP), pages 1085–1089, 2021. doi: 10.1109/ICASSP39728.2021.9414145.

[31] Charlie Windolf, Angelique C Paulk, Yoav Kfir, Eric Trautmann, Domokos Meszéna, William Muñoz, Irene Caprara, Mohsen Jamali, Julien Boussard, Ziv M Williams, et al. Robust online multiband drift estimation in electrophysiology data. In ICASSP 2023-2023 IEEE International Conference on Acoustics, Speech and Signal Processing (ICASSP), pages 1–5. IEEE, 2023.

[32] IBL, Kush Banga, Julius Benson, Niccolò Bonacchi, Sebastian A Bruijns, Rob Campbell, Gaë lle A Chapuis, Anne K Churchland, M Felicia Davatolhagh, Hyun Dong Lee, Mayo Faulkner, Fei Hu, Julia Hunterberg, Anup Khanal, Christopher Krasniak, Guido T Meijer, Nathaniel J Miska, Zeinab Mohammadi, Jean-Paul Noel, Liam Paninski, Alejandro Pan-Vazquez, Noam Roth, Michael Schartner, Karolina Socha, Nicholas A Steinmetz, Karel Svoboda, Marsa Taheri, Anne E Urai, Miles Wells, Steven J West, Matthew R Whiteway, Olivier Winter, and Ilana B Witten. Reproducibility of in-vivo electrophysiological measurements in mice. bioRxiv, 2022. doi: 10.1101/2022.05.09.491042. URL https://www.biorxiv.org/content/early/2022/05/12/2022.05.09.491042.

[33] Eric M Trautmann, Janis K Hesse, Gabriel M Stine, Ruobing Xia, Shude Zhu, Daniel J O’Shea, Bill Karsh, Jennifer Colonell, Frank F Lanfranchi, Saurabh Vyas, et al. Large-scale high-density brain-wide neural recording in nonhuman primates. bioRxiv, pages 2023–02, 2023.

[34] Zhiwen Ye, Andrew M Shelton, Jordan R Shaker, Julien M Boussard, Jennifer Colonell, Sahar Minavi, Susu Chen, Charlie Windolf, Cole Hurwitz, Tomoyuki Namima, Frederico Pedraja, Shahaf Weiss, Bogdan Raducanu, Torbjørn Ness, Gaute T Einevoll, Gilles Laurent, Nathaniel B Sawtell, Wyeth Bair, Anitha Pasupathy, Carolina Mora-Lopez, Barun Dutta, Liam Paninski, Joshua H Siegle, Christof Koch, Shawn R Olsen, Timothy D Harris, and Nicholas A Steinmetz. Ultra-high density electrodes improve detection, yield, and cell type specificity of brain recordings. bioRxiv, 2023. doi: 10.1101/2023.08.23.554527. URL https://www.biorxiv.org/content/early/2023/08/24/2023.08.23.554527.

[35] Julien Boussard, Charlie Windolf, Cole Hurwitz, Hyun Dong Lee, Han Yu, Olivier Winter, and Liam Paninski. Dartsort: A modular drift tracking spike sorter for high-density multi-electrode probes. bioRxiv, 2023. doi: 10.1101/2023.08.11.553023. URL https://www.biorxiv.org/content/early/2023/08/14/2023.08.11.553023.

[36] Julien Boussard, Erdem Varol, Hyun Dong Lee, Nishchal Dethe, and Liam Paninski. Three-dimensional spike localization and improved motion correction for Neuropixels recordings. In M. Ranzato, A. Beygelzimer, Y. Dauphin, P.S. Liang, and J. Wortman Vaughan, editors, Advances in Neural Information Processing Systems, volume 34, pages 22095–22105. Curran Associates, Inc., 2021. URL https://proceedings.neurips.cc/paper_files/paper/2021/file/b950ea26ca12daae142bd74dba4427c8-Paper.pdf.

[37] Ferenc Mechler, Jonathan D. Victor, Ifije Ohiorhenuan, Anita M. Schmid, and Qin Hu. Three-dimensional localization of neurons in cortical tetrode recordings. Journal of Neurophysiology, 106(2):828–848, August 2011. doi: 10.1152/jn.00515.2010. URL 10.1152/jn.00515.2010.

[38] Cole Hurwitz, Kai Xu, Akash Srivastava, Alessio Buccino, and Matthias Hennig. Scalable spike source localization in extracellular recordings using amortized variational inference. In H. Wallach, H. Larochelle, A. Beygelzimer, F. d’Alché-Buc, E. Fox, and R. Garnett, editors, Advances in Neural Information Processing Systems, volume 32. Curran Associates, Inc., 2019. URL https://proceedings.neurips.cc/paper_files/paper/2019/file/f12f2b34a0c3174269c19e21c07dee68-Paper.pdf.

[39] Alessio P Buccino, Michael Kordovan, Torbjørn V Ness, Benjamin Merkt, Philipp D Häfliger, Marianne Fyhn, Gert Cauwenberghs, Stefan Rotter, and Gaute T Einevoll. Combining biophysical modeling and deep learning for multielectrode array neuron localization and classification. J. Neurophysiol., 120(3):1212–1232, September 2018.

[40] Csaba Horváth, Lili Fanni Tóth, István Ulbert, and Richárd Fiáth. Dataset of cortical activity recorded with high spatial resolution from anesthetized rats. Scientific Data, 8(1), July 2021. doi: 10.1038/s41597-021-00970-3. URL 10.1038/s41597-021-00970-3.

[41] I. Tal and M. Abeles. Cleaning MEG artifacts using external cues. Journal of Neuroscience Methods, 217(1-2):31–38, July 2013. doi: 10.1016/j.jneumeth.2013.04.002. URL 10.1016/j.jneumeth.2013.04.002.

[42] Marius Klug and Niels A. Kloosterman. Zapline-plus: A zapline extension for automatic and adaptive removal of frequency-specific noise artifacts in M/EEG. Human Brain Mapping, 43(9): 2743–2758, March 2022. doi: 10.1002/hbm.25832. URL 10.1002/hbm.25832.

[43] Alain de Cheveigné. ZapLine: A simple and effective method to remove power line artifacts. NeuroImage, 207:116356, February 2020. doi: 10.1016/j.neuroimage.2019.116356. URL 10.1016/j.neuroimage.2019.116356.

[44] István Ulbert, Eric Halgren, Gary Heit, and George Karmos. Multiple microelectrode-recording system for human intracortical applications. Journal of Neuroscience Methods, 106(1):69–79, mar 2001. doi: 10.1016/s0165-0270(01)00330-2. URL 10.1016%2Fs0165-0270%2801%2900330-2.

[45] Richárd Csercsa, Balázs Dombovári, Dániel Fabó, Lucia Wittner, Loránd Erőss, László Entz, András Sólyom, György Rásonyi, Anna Szu? cs, Anna Kelemen, Rita Jakus, Vera Juhos, László Grand, Andor Magony, Péter Halász, Tamás F. Freund, Zsófia Maglóczky, Sydney S. Cash, László Papp, György Karmos, Eric Halgren, and István Ulbert. Laminar analysis of slow wave activity in humans. Brain, 133(9):2814–2829, jul 2010. doi: 10.1093/brain/awq169. URL 10.1093%2Fbrain%2Fawq169.

[46] Patrick Baudena, Eric Halgren, Gary Heit, and Jeffrey M. Clarke. Intracerebral potentials to rare target and distractor auditory and visual stimuli. III. frontal cortex. Electroencephalography and Clinical Neurophysiology, 94(4):251–264, April 1995. doi: 10.1016/0013-4694(95)98476-o. URL 10.1016/0013-4694(95)98476-o.

[47] Daniel Fabo, Virag Bokodi, Johanna-Petra Szabó, Emilia Tóth, Pariya Salami, Corey J. Keller, Boglárka Hajnal, Thomas Thesen, Orrin Devinsky, Werner Doyle, Ashesh Mehta, Joseph Madsen, Emad Eskandar, Lorand Erőss, István Ulbert, Eric Halgren, and Sydney S. Cash. The role of superficial and deep layers in the generation of high frequency oscillations and interictal epileptiform discharges in the human cortex. Scientific Reports, 13(1), June 2023. doi: 10.1038/s41598-022-22497-2. URL 10.1038/s41598-022-22497-2.

[48] Laura D. Lewis, ShiNung Ching, Veronica S. Weiner, Robert A. Peterfreund, Emad N. Eskandar, Sydney S. Cash, Emery N. Brown, and Patrick L. Purdon. Local cortical dynamics of burst suppression in the anaesthetized brain. Brain, 136(9):2727–2737, July 2013. doi: 10.1093/brain/awt174. URL 10.1093/brain/awt174.

[49] M. Brandon Westover, Mouhsin M. Shafi, ShiNung Ching, Jessica J. Chemali, Patrick L. Purdon, Sydney S. Cash, and Emery N. Brown. Real-time segmentation of burst suppression patterns in critical care EEG monitoring. Journal of Neuroscience Methods, 219(1):131–141, September 2013. doi: 10.1016/j.jneumeth.2013.07.003. URL 10.1016/j.jneumeth.2013.07.003.

[50] Pariya Salami, Mia Borzello, Mark A. Kramer, M. Brandon Westover, and Sydney S. Cash. Quantifying seizure termination patterns reveals limited pathways to seizure end. Neurobiology of Disease, 165:105645, April 2022. doi: 10.1016/j.nbd.2022.105645. URL 10.1016/j.nbd.2022.105645.

[51] Yina Wei, Anirban Nandi, Xiaoxuan Jia, Joshua H Siegle, Daniel Denman, Soo Yeun Lee, Anatoly Buchin, Werner Van Geit, Clayton P Mosher, Shawn Olsen, et al. Associations between in vitro, in vivo and in silico cell classes in mouse primary visual cortex. Nature Communications, 14(1):2344, 2023.

[52] JinHyung Lee, Catalin Mitelut, Hooshmand Shokri, Ian Kinsella, Nishchal Dethe, Shenghao Wu, Kevin Li, Eduardo Blancas Reyes, Denis Turcu, Eleanor Batty, et al. Yass: Yet another spike sorter applied to large-scale multi-electrode array recordings in primate retina. BioRxiv, pages 2020–03, 2020.

[53] D. N. Hill, S. B. Mehta, and D. Kleinfeld. Quality metrics to accompany spike sorting of extracellular signals. Journal of Neuroscience, 31(24):8699–8705, June 2011. doi: 10.1523/jneurosci.0971-11.2011. URL 10.1523/jneurosci.0971-11.2011.

[54] Augustine Xiaoran Yuan, Jennifer Colonell, Anna Lebedeva, Adam Charles, and Timothy Harris. Multi-day neuron tracking in high density electrophysiology recordings using EMD. bioRxiv, August 2023. doi: 10.1101/2023.08.03.551724. URL https://www.biorxiv.org/content/10.1101/2023.08.03.551724.

[55] Benjamin H. Savitzky, Ismail El Baggari, Colin B. Clement, Emily Waite, Berit H. Goodge, David J. Baek, John P. Sheckelton, Christopher Pasco, Hari Nair, Nathaniel J. Schreiber, Jason Hoffman, Alemayehu S. Admasu, Jaewook Kim, Sang-Wook Cheong, Anand Bhattacharya, Darrell G. Schlom, Tyrel M. McQueen, Robert Hovden, and Lena F. Kourkoutis. Image registration of low signal-to-noise cryo-stem data. Ultramicroscopy, 191:56–65, 2018. ISSN 0304-3991. doi: 10.1016/j.ultramic.2018.04.008. URL https://www.sciencedirect.com/science/article/pii/S0304399117304369.

[56] Liam Paninski, Yashar Ahmadian, Daniel Gil Ferreira, Shinsuke Koyama, Kamiar Rahnama Rad, Michael Vidne, Joshua Vogelstein, and Wei Wu. A new look at state-space models for neural data. Journal of Computational Neuroscience, 29(1-2):107–126, August 2009. doi: 10.1007/s10827-009-0179-x. URL 10.1007/s10827-009-0179-x.

[57] Leland McInnes, John Healy, and Steve Astels. hdbscan: Hierarchical density based clustering. Journal of Open Source Software, 2(11):205, 2017. doi: 10.21105/joss.00205. URL 10.21105/joss.00205.

[58] Carl Gold, Cyrille C Girardin, Kevan AC Martin, and Christof Koch. High-amplitude positive spikes recorded extracellularly in cat visual cortex. Journal of neurophysiology, 102(6):3340–3351, 2009.

[59] Jeremy M Barry. Axonal activity in vivo: technical considerations and implications for the exploration of neural circuits in freely moving animals. Frontiers in neuroscience, 9:153, 2015.

[60] Angelique C Paulk, Jimmy C Yang, Daniel R Cleary, Daniel J Soper, Mila Halgren, Alexandra R O’Donnell, Sang Heon Lee, Mehran Ganji, Yun Goo Ro, Hongseok Oh, Lorraine Hossain, Jihwan Lee, Youngbin Tchoe, Nicholas Rogers, Kivilcim Kiliç, Sang Baek Ryu, Seung Woo Lee, John Hermiz, Vikash Gilja, István Ulbert, Daniel Fabó, Thomas Thesen, Werner K Doyle, Orrin Devinsky, Joseph R Madsen, Donald L Schomer, Emad N Eskandar, Jong Woo Lee, Douglas Maus, Anna Devor, Shelley I Fried, Pamela S Jones, Brian V Nahed, Sharona Ben-Haim, Sarah K Bick, Robert Mark Richardson, Ahmed M Raslan, Dominic A Siler, Daniel P Cahill, Ziv M Williams, G Rees Cosgrove, Shadi A Dayeh, and Sydney S Cash. Microscale physiological events on the human cortical surface. Cerebral Cortex, 31(8):3678–3700, March 2021. doi: 10.1093/cercor/bhab040. URL 10.1093/cercor/bhab040.

[61] Mohsen Jamali, Ben Grannan, Keren Haroush, Ziev B. Moses, Emad N. Eskandar, Todd Herrington, Shaun Patel, and Ziv M. Williams. Dorsolateral prefrontal neurons mediate subjective decisions and their variation in humans. Nature Neuroscience, 22(6):1010–1020, April 2019. doi: 10.1038/s41593-019-0378-3. URL 10.1038/s41593-019-0378-3.

[62] Jimmy C. Yang, Angelique C. Paulk, Pariya Salami, Sang Heon Lee, Mehran Ganji, Daniel J. Soper, Daniel Cleary, Mirela Simon, Douglas Maus, Jong Woo Lee, Brian V. Nahed, Pamela S. Jones, Daniel P. Cahill, Garth Rees Cosgrove, Catherine J. Chu, Ziv Williams, Eric Halgren, Shadi Dayeh, and Sydney S. Cash. Microscale dynamics of electrophysiological markers of epilepsy. Clinical Neurophysiology, 132(11):2916–2931, November 2021. doi: 10.1016/j.clinph.2021.06.024. URL 10.1016/j.clinph.2021.06.024.

[63] Carolina Mora Lopez, Jan Putzeys, Bogdan Cristian Raducanu, Marco Ballini, Shiwei Wang, Alexandru Andrei, Veronique Rochus, Roeland Vandebriel, Simone Severi, Chris Van Hoof, Silke Musa, Nick Van Helleputte, Refet Firat Yazicioglu, and Srinjoy Mitra. A neural probe with up to 966 electrodes and up to 384 configurable channels in 0.13 μm SOI CMOS. IEEE Transactions on Biomedical Circuits and Systems, 11(3):510–522, June 2017. doi: 10.1109/tbcas.2016.2646901. URL 10.1109/tbcas.2016.2646901.

[64] Joshua H Siegle, Aarón Cuevas López, Yogi A Patel, Kirill Abramov, Shay Ohayon, and Jakob Voigts. Open ephys: an open-source, plugin-based platform for multichannel electrophysiology. Journal of Neural Engineering, 14(4):045003, jun 2017. doi: 10.1088/1741-2552/aa5eea. URL 10.1088%2F1741-2552%2Faa5eea.

[65] David H. Brainard. The psychophysics toolbox. Spatial Vision, 10(4):433–436, 1997. doi: 10.1163/156856897x00357. URL 10.1163/156856897x00357.

[66] R. C. Reid, J. D. Victor, and R. M. Shapley. The use of m-sequences in the analysis of visual neurons: Linear receptive field properties. Visual Neuroscience, 14(6):1015–1027, November 1997. doi: 10.1017/s0952523800011743. URL 10.1017/s0952523800011743.

[67] Erich E. Sutter. Imaging visual function with the multifocal m-sequence technique. Vision Research, 41(10-11):1241–1255, May 2001. doi: 10.1016/s0042-6989(01)00078-5. URL 10.1016/s0042-6989(01)00078-5.

[68] Giedrius T. Buračas and Geoffrey M. Boynton. Efficient design of event-related fMRI experiments using m-sequences. NeuroImage, 16(3):801–813, July 2002. doi: 10.1006/nimg.2002.1116. URL 10.1006/nimg.2002.1116.

[69] Hitoshi Tabuchi, Tsuranu Yokoyama, Masahiro Shimogawara, Kunihiko Shiraki, Eiichiro Nagasaka, and Tokuhiko Miki. Study of the visual evoked magnetic field with the m-sequence technique. Invest. Ophthalmol. Vis. Sci., 43(6):2045–2054, June 2002.

[70] Robert Oostenveld, Pascal Fries, Eric Maris, and Jan-Mathijs Schoffelen. FieldTrip: Open source software for advanced analysis of MEG, EEG, and invasive electrophysiological data. Computational Intelligence and Neuroscience, 2011:1–9, 2011. doi: 10.1155/2011/156869. URL 10.1155/2011/156869.

[71] Richárd Fiáth, Bogdan Cristian Raducanu, Silke Musa, Alexandru Andrei, Carolina Mora Lopez, Chris van Hoof, Patrick Ruther, Arno Aarts, Domonkos Horváth, and István Ulbert. A silicon-based neural probe with densely-packed low-impedance titanium nitride microelectrodes for ultrahigh-resolution in vivo recordings. Biosens. Bioelectron., 106:86–92, May 2018.

[72] J. Anthony Movshon, Lynne Kiorpes, Michael J. Hawken, and James R. Cavanaugh. Functional maturation of the macaque’s lateral geniculate nucleus. The Journal of Neuroscience, 25(10): 2712–2722, mar 2005. doi: 10.1523/jneurosci.2356-04.2005. URL 10.1523%2Fjneurosci.2356-04.2005.

[73] Long Yang, Kwang Lee, Jomar Villagracia, and Sotiris C Masmanidis. Open source sili-con microprobes for high throughput neural recording. Journal of Neural Engineering, 17(1): 016036, jan 2020. doi: 10.1088/1741-2552/ab581a. URL 10.1088%2F1741-2552%2Fab581a.

[74] Adam Paszke, Sam Gross, Francisco Massa, Adam Lerer, James Bradbury, Gregory Chanan, Trevor Killeen, Zeming Lin, Natalia Gimelshein, Luca Antiga, Alban Desmaison, Andreas Kopf, Edward Yang, Zachary DeVito, Martin Raison, Alykhan Tejani, Sasank Chilamkurthy, Benoit Steiner, Lu Fang, Junjie Bai, and Soumith Chintala. Pytorch: An imperative style, high-performance deep learning library. In H. Wallach, H. Larochelle, A. Beygelzimer, F. d’Alché-Buc, E. Fox, and R. Garnett, editors, Advances in Neural Information Processing Systems 32, pages 8024–8035. Curran Associates, Inc., 2019. URL http://papers.neurips.cc/paper/9015-pytorch-an-imperative-style-high-performance-deep-learning-library.pdf.

[75] Pauli Virtanen, Ralf Gommers, Travis E. Oliphant, Matt Haberland, Tyler Reddy, David Cournapeau, Evgeni Burovski, Pearu Peterson, Warren Weckesser, Jonathan Bright, Stéfan J. van der Walt, Matthew Brett, Joshua Wilson, K. Jarrod Millman, Nikolay Mayorov, Andrew R. J. Nelson, Eric Jones, Robert Kern, Eric Larson, C J Carey, İlhan Polat, Yu Feng, Eric W. Moore, Jake VanderPlas, Denis Laxalde, Josef Perktold, Robert Cimrman, Ian Henriksen, E. A. Quintero, Charles R. Harris, Anne M. Archibald, Antônio H. Ribeiro, Fabian Pedregosa, Paul van Mulbregt, and SciPy 1.0 Contributors. SciPy 1.0: Fundamental Algorithms for Scientific Computing in Python. Nature Methods, 17:261–272, 2020. doi: 10.1038/s41592-019-0686-2.

