## Supplementary Information for "DREDge: robust motion correction for high-density extracellular recordings across species"

### 7 Supplementary Information

#### 7.1 Proof of equation (2)

Since the objective in equation (1) decomposes over nonrigid blocks $b = 1, \ldots, B$, it suffices to solve the rigid problem

$$\arg\min_{\mathbf{p} \in \mathbb{R}^T} L(\mathbf{P}); \qquad L(\mathbf{p}) = \frac{1}{4} \left\| \mathbf{D} - (\mathbf{p}\mathbf{1}_T^\top - \mathbf{1}_T \mathbf{p}^\top) \right\|_2^2, \tag{9}$$

where $\mathbf{D} \in \mathbb{R}^{T \times T}$ is an antisymmetric matrix. After simplifying by using this antisymmetry, the gradient of $L$ is given by

$$\nabla_{\mathbf{p}} L(\mathbf{p}) = T\mathbf{p} - (\mathbf{p}^\top \mathbf{1}_T)\mathbf{1}_T - \mathbf{D}\mathbf{1}_T. \tag{10}$$

Let $\mathbf{q} = \frac{1}{T}\mathbf{D}\mathbf{1}_T$ be the row means of $\mathbf{D}$, and note that since $\mathbf{D}$ is antisymmetric, $\mathbf{q}$ sums to 0. Plugging $\mathbf{q}$ for $\mathbf{p}$ above leads to $\nabla_{\mathbf{p}} L(\mathbf{p})|_{\mathbf{p}=\mathbf{q}} = 0$, so that these row means cause the gradient to vanish.

Differentiating again, the Hessian is

$$\mathbf{H}_L(\mathbf{p})_{tt'} = T\mathbf{I}_T - \mathbf{1}_T \mathbf{1}_T^\top \tag{11}$$

which is symmetric and diagonally dominant and therefore positive semidefinite, but not positive definite since $\mathbf{1}_T$ is in the null space.

Thus $L$ is (not strictly) convex and $\mathbf{q}$ is a minimizer of $L$; the rest of the minimizers are of the form $\mathbf{q} + c$ for $c \in \mathbb{R}$.

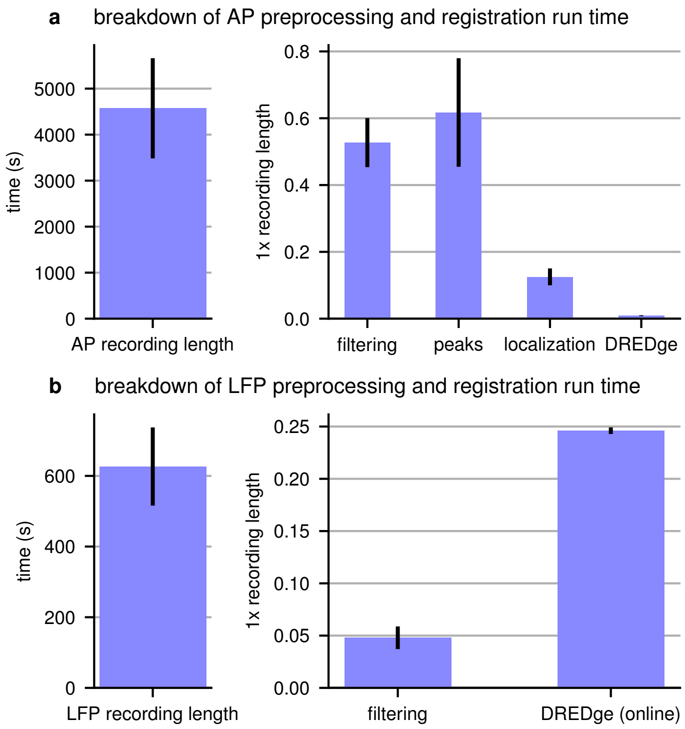


Supplementary Figure 1: **Measuring DREDge's run time in AP and LFP. a**, run time breakdown in acute recordings made by the IBL[32]. These $n = 8$ recordings are over an hour on average (left, $76 \pm 18$ min), and were preprocessed according to Section 4.1. Here, *filtering* indicates the raw data pipeline of IBL et al.[20] (including temporal highpass filtering, dead channel detection and interpolation, spatial highpass filtering, and standardization); *peaks* and *localization* are the peak detection, denoising, and localization pipeline of[35] and Boussard et al.[36]. DREDge's runtime comes in at a small fraction of both real recording time and the overall pipeline's run time. **b**, run time breakdown in the tip-layout recordings of Paulk et al.[15]. These $n = 7$ recordings were around 10min long ($10.4 \pm 1.8$min, left), and were preprocessed according to Section 4.2, leading to filtered preprocessed recordings sampled at 250Hz. Online nonrigid motion estimation at 250Hz temporal resolution ran at roughly one quarter of real time (right). These benchmarks were conducted on academic cluster hardware: **a**, Intel Skylake 6148 processors and NVIDIA V100 GPU; **b**, Intel Xeon Gold 6126 processors and NVIDIA V100 GPU. Bar heights are means and intervals are standard deviations.

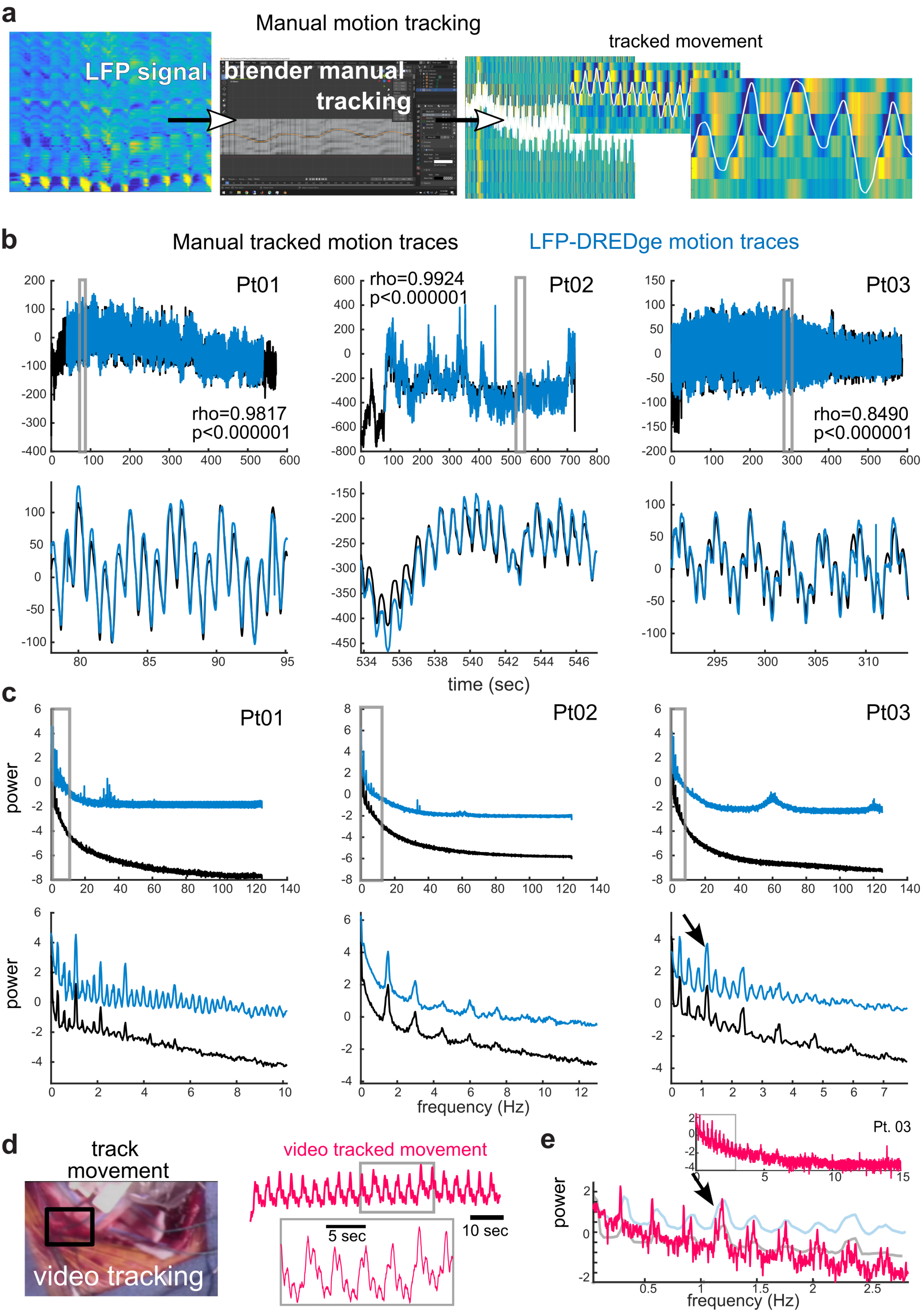
a
Manual motion tracking
LFP signal
blender manual tracking
tracked movement
b
Manual tracked motion traces
LFP-DREDge motion traces
Pt01
rho=0.9817
p<0.000001
Pt02
rho=0.9924
p<0.000001
Pt03
rho=0.8490
p<0.000001
time (sec)
c
Pt01
Pt02
Pt03
power
frequency (Hz)
d
track movement
video tracking
video tracked movement
5 sec
10 sec
e
Pt. 03
power
frequency (Hz)

Supplementary Figure 2: **Motion tracking comparisons.** **a**, Manual tracking method using tracing in Blender to track voltage changes in the LFP through time. **b**, Motion traces tracked manually (black lines) versus those used with DREDge (blue) in three data sets. Top row: full traces of the data through time. Bottom row: zoomed-in views from the grey-outlined boxes shown in the top row. **c**, Power spectra of the motion traces tracked manually (black lines) versus those used with DREDge (blue) in three data sets. Top row: full power spectra of the motion track through time. Bottom row: zoomed-in views from the grey-outlined boxes shown in the top row. **d**, Motion tracked using video of the brain movements during a Neuropixels recording and an open craniotomy. Left: video of the intraoperative recording and the pumping evident in the CSF surrounding the electrode was tracked through time. Right: Simultaneous traces of the video-tracked movements and the LFP-tracked movements in the same patient (Pt. 03) at two different scales. **e**, Power spectrum of the motion tracked using video of the brain movements during a Neuropixels recording and an open craniotomy. Top spectrum: zoomed-out view. Bottom power spectrum plot: zoomed-in view of the video motion tracked overlaid on the same spectra of the DREDge and manual motion tracked for the same case (Pt03).

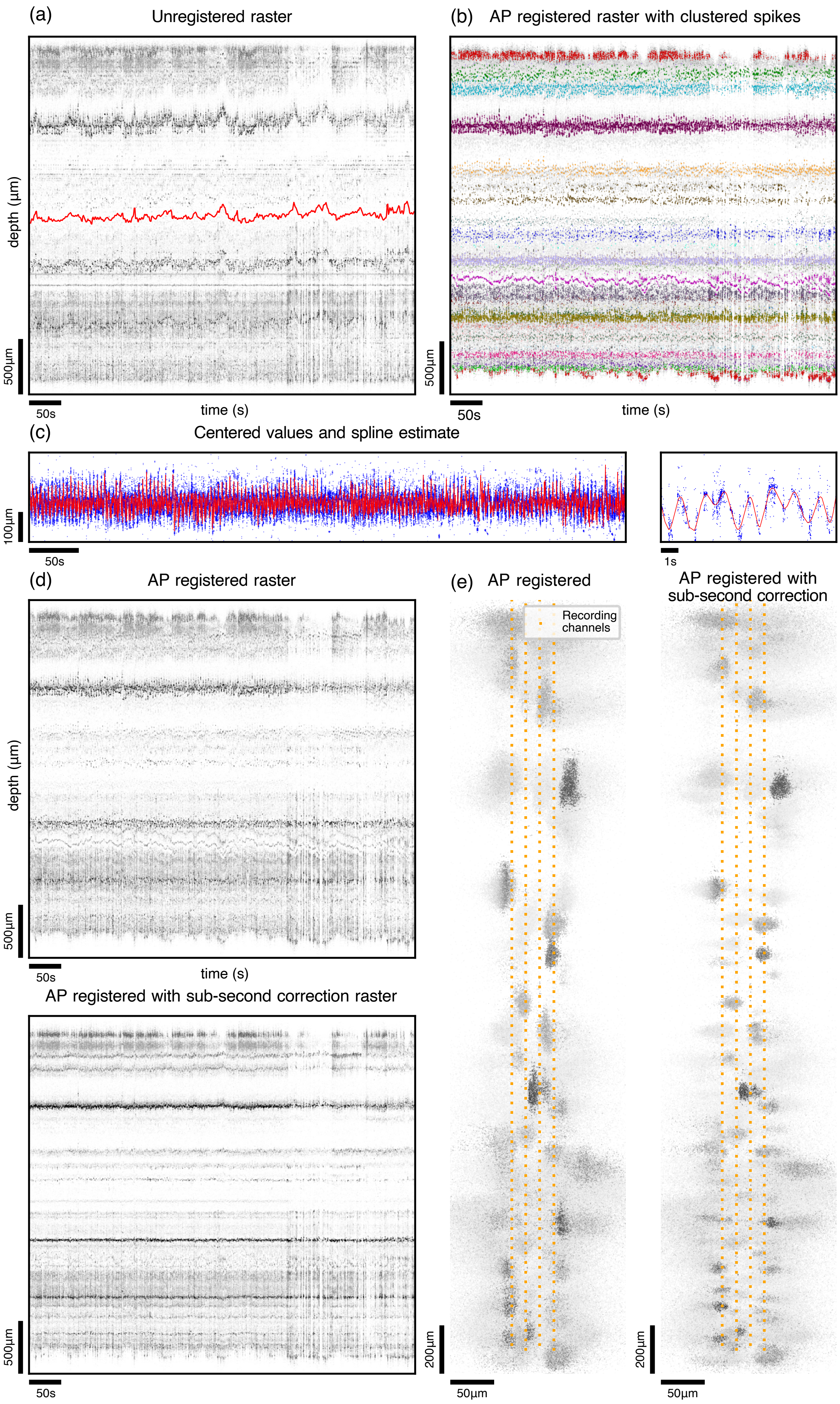
(a)
Unregistered raster
depth (μm)
500μm
50s
time (s)
(b)
AP registered raster with clustered spikes
500μm
50s
time (s)
(c)
Centered values and spline estimate
100μm
50s
1s
(d)
AP registered raster
depth (μm)
500μm
50s
time (s)
AP registered with sub-second correction raster
500μm
50s
(e)
AP registered
AP registered with sub-second correction
Recording channels
200μm
50μm
200μm
50μm

1451

Supplementary Figure 3: **Spline registration** (a) Spike raster before registration, with AP DREDge displacement estimate overlaid in red. (b) Spike raster after registration (grey). Colored spikes represent high amplitude spikes that were clustered by HDBSCAN using their registered positions and amplitudes. (c) Left, centered positions of the clustered spikes (blue) with the spline fit, that captures the sub-second displacement, overlaid in red. Right, zoom of this time-series between times 100 and 110 seconds. (d) Top, AP registered spike raster. Bottom, AP registered spike raster corrected using the spline estimate of sub-second displacement. (e) Localizations of detected spikes after AP DREDge registration (left), and spline correction (right). Both high and low amplitude clusters are now much sharper.

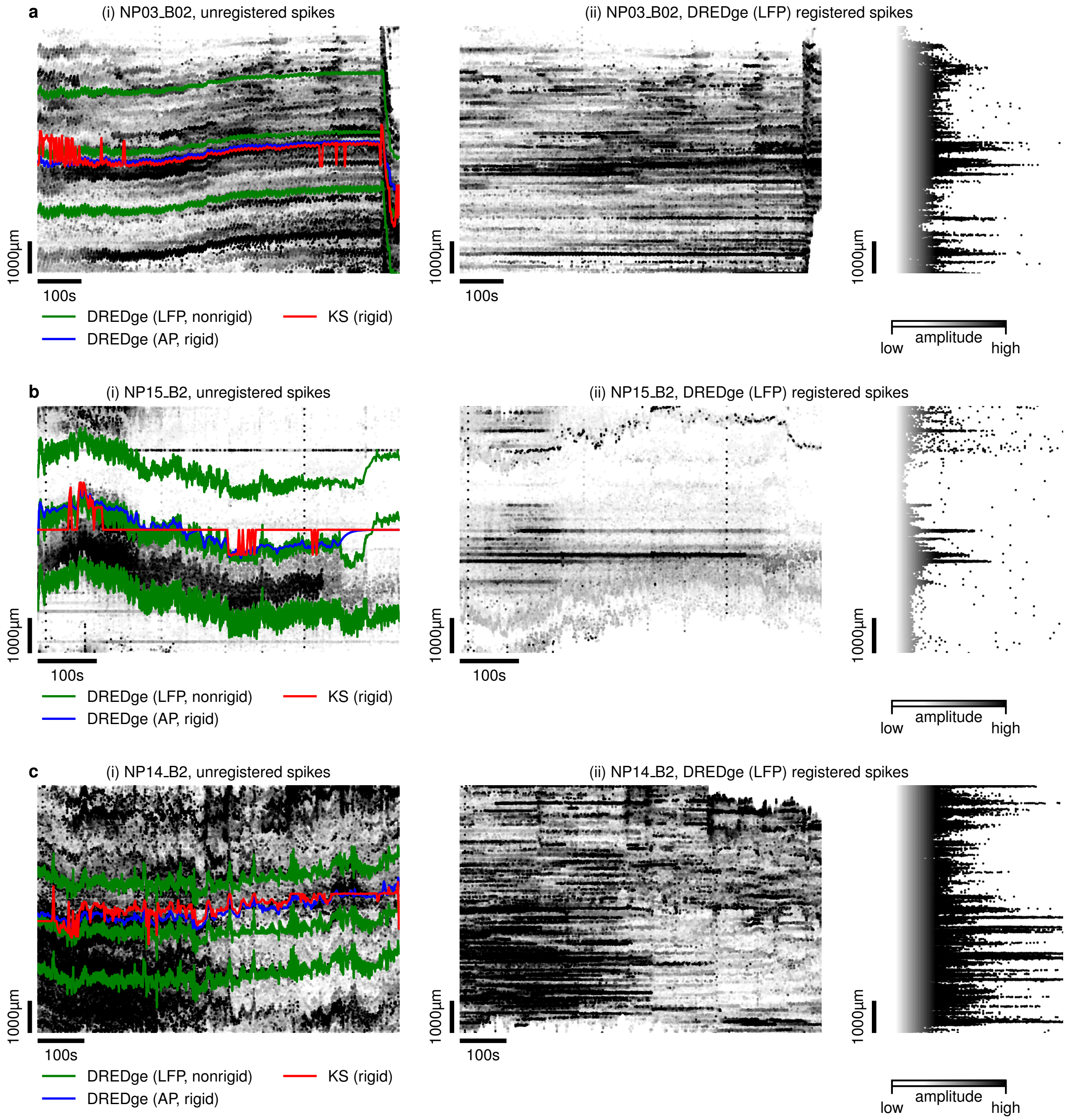


Supplementary Figure 4: **Additional cases where LFP-based motion tracking discovers signal in noisy spiking data.** In addition to the dataset shown in Fig. 2.c, here we tracked motion in three more human intraoperative datasets from Chung et al.[16], recorded with long channel configurations. In these cases, AP-based motion estimation with DREDge and KS struggled (i; blue and red), but nonrigid LFP-based motion tracking was able to follow extensive motion (i, green), leading to stabilized unit traces in plots of spike depth against time and amplitude (ii).

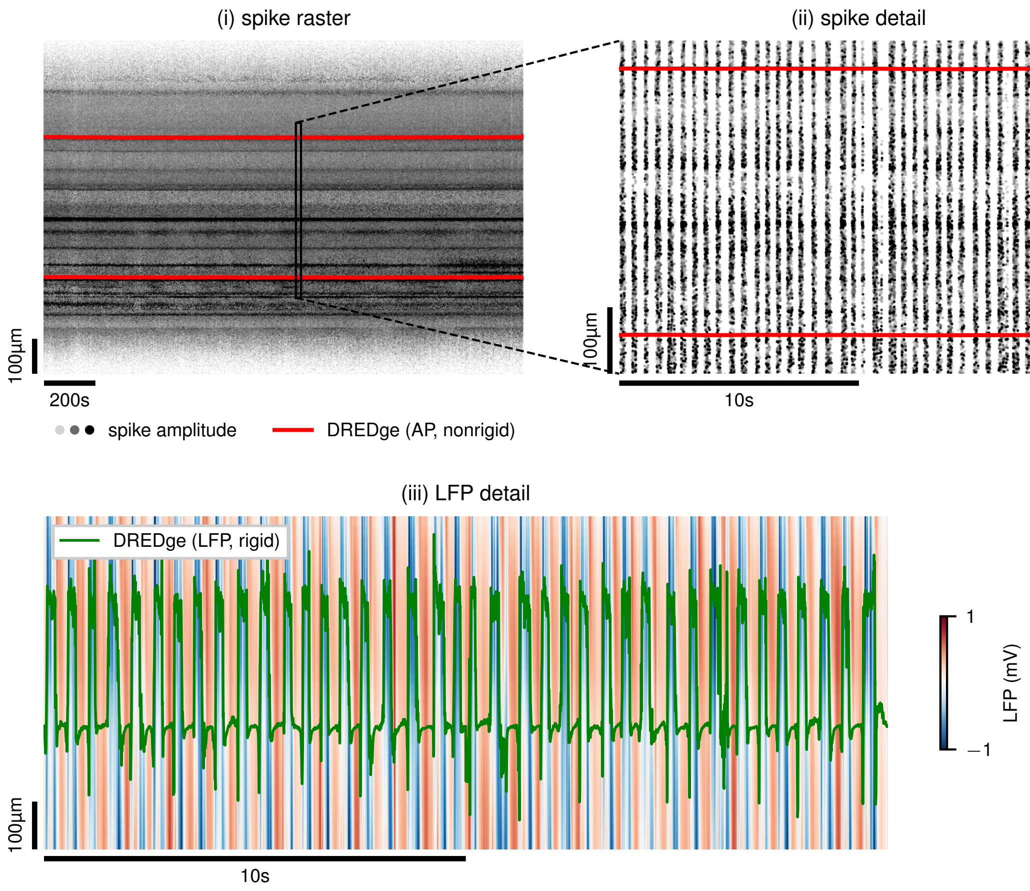


1453 Supplementary Figure 5: **Relying on spike-based motion estimation when LFP is dominated by other signals.** In high-resolution recordings in ketamine/xylazine-anaesthetized rat[40], slow-wave activity composed of alternating up/down states can be observed across the length of the probe in the AP and LFP bands (visible as vertical stripes of oscillating spiking and LFP activity in ii and iii). At 1Hz temporal resolution, DREDge's AP-based motion tracking smooths over the much faster up/down oscillations (i and ii). In the LFP band, however, these oscillations dominate the signal to the extent that DREDge's high-resolution motion tracking also oscillates along the up/down transitions (iii).

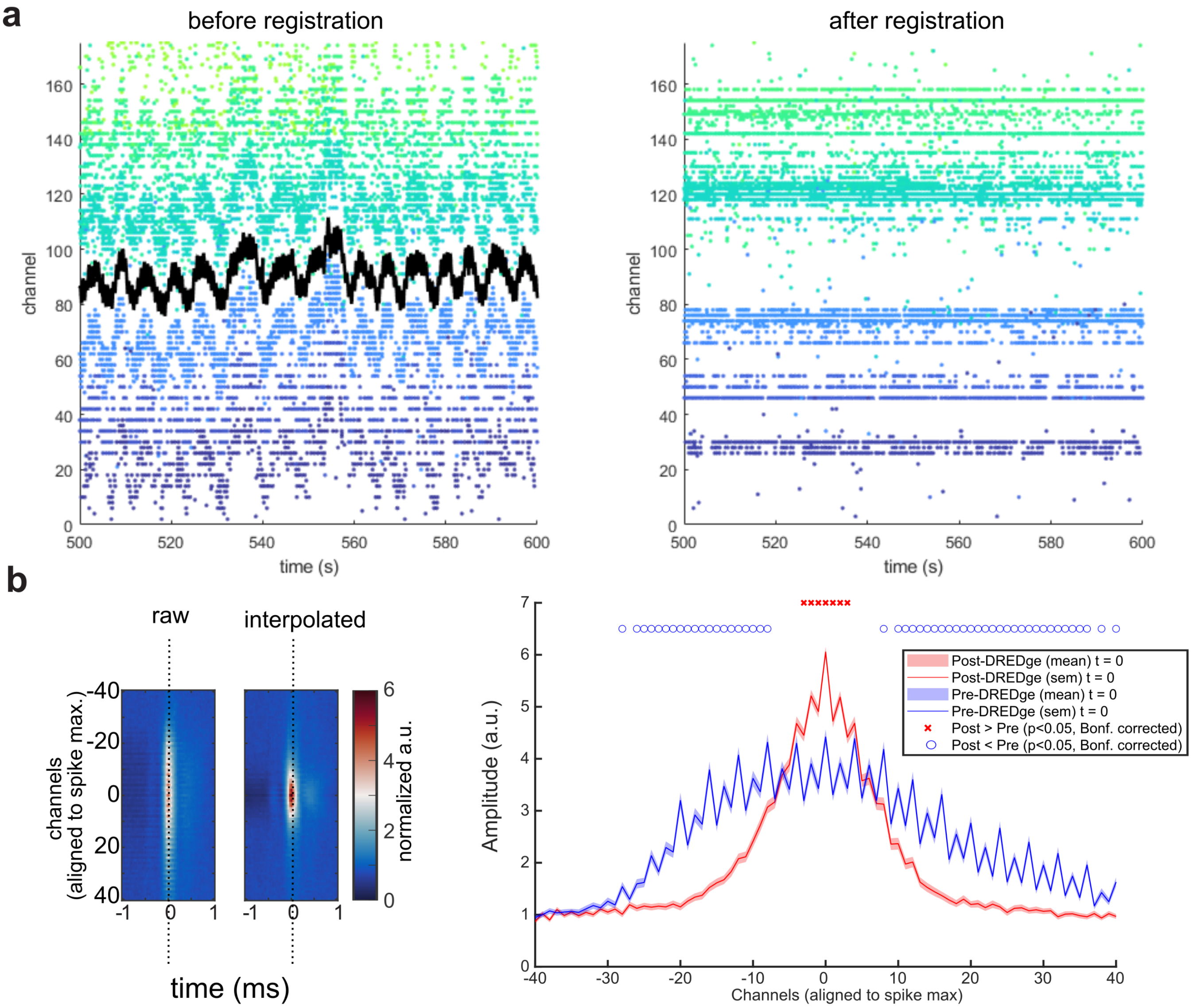


Supplementary Figure 6: **Clustering before and after corrections. a**, A subset of spike detections and sorted units (with different single unit clusters color coded as dots) across channels before (top) and after (bottom) registration with an optimized DREDge motion estimate (black line). Note the emergence of aligned spikes on the bottom panel. **b**, Average spatial distribution of spike clusters when non-interpolated (left) and 250 Hz-interpolated (middle) from Fig. 2.g, where a single slice taken from the middle time range (dotted lines in the left) is then compared across clusters for the amplitude relative to the spatial spread (right plot). The average amplitudes and standard error of the mean (SEM) curves of the spike amplitudes for the raw data versus the motion corrected data set were compared using a two-sided two sample *t*-test at each distance from center, Bonferroni corrected with a threshold of $p < 0.05$. Red 'x' markers indicate the distance from the peak center waveform where the motion corrected spike waveforms are higher than the raw sampled waveforms. Blue 'o' markers indicate the distance from the peak center waveform where the raw spike waveforms are higher than the motion corrected sampled waveforms.

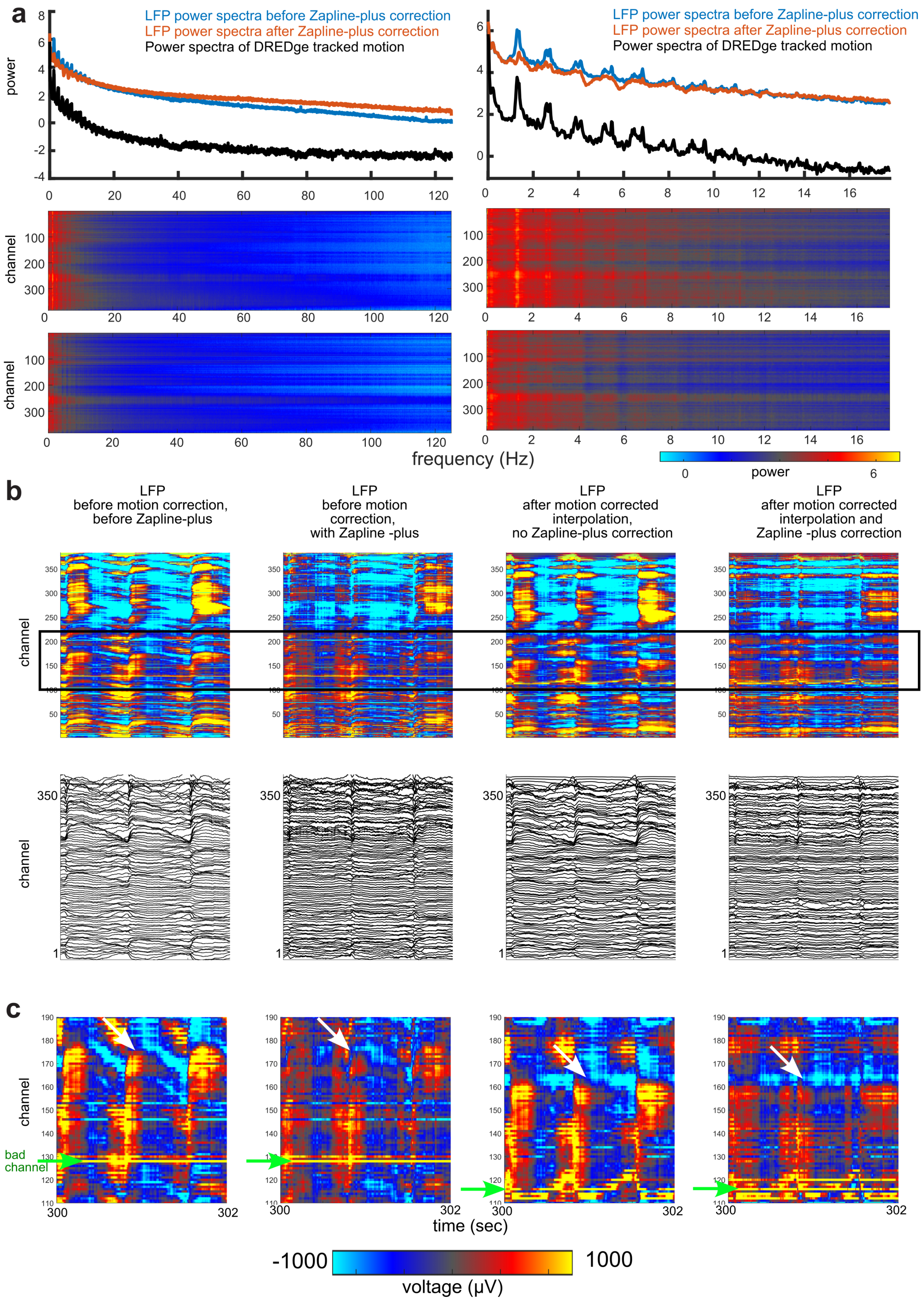

a
LFP power spectra before Zapline-plus correction
LFP power spectra after Zapline-plus correction
Power spectra of DREDge tracked motion
power
channel
frequency (Hz)
power
b
LFP before motion correction, before Zapline-plus
LFP before motion correction, with Zapline -plus
LFP after motion corrected interpolation, no Zapline-plus correction
LFP after motion corrected interpolation and Zapline -plus correction
channel
c
channel
bad channel
time (sec)
-1000
1000
voltage (μV)


1455

Supplementary Figure 7: **Motion correction interpolation of LFP in human brain activity.** **a**, Zapline-plus correction of the spectral features after motion correction. **b**, Steps involving taking the raw spontaneous LFP (left), raw data with Zapline-plus correction of low-frequency peaks (second column), motion-correction interpolation of LFP without Zapline-plus correction (third column), motion-correction interpolation of LFP with Zapline-plus correction (fourth column). **c**, Zoomed in view of the raw spontaneous LFP (left), raw data with Zapline-plus correction of low-frequency peaks (second column), motion-correction interpolation of LFP without Zapline-plus correction (third column), motion-correction interpolation of LFP with Zapline-plus correction (fourth column). The view comes from the box highlighted in b. Green arrows indicate a bad channel (yellow band) which then is shifted up and down following motion-correction interpolation. White arrows indicate an LFP band of activity which is then flattened following the corrections.

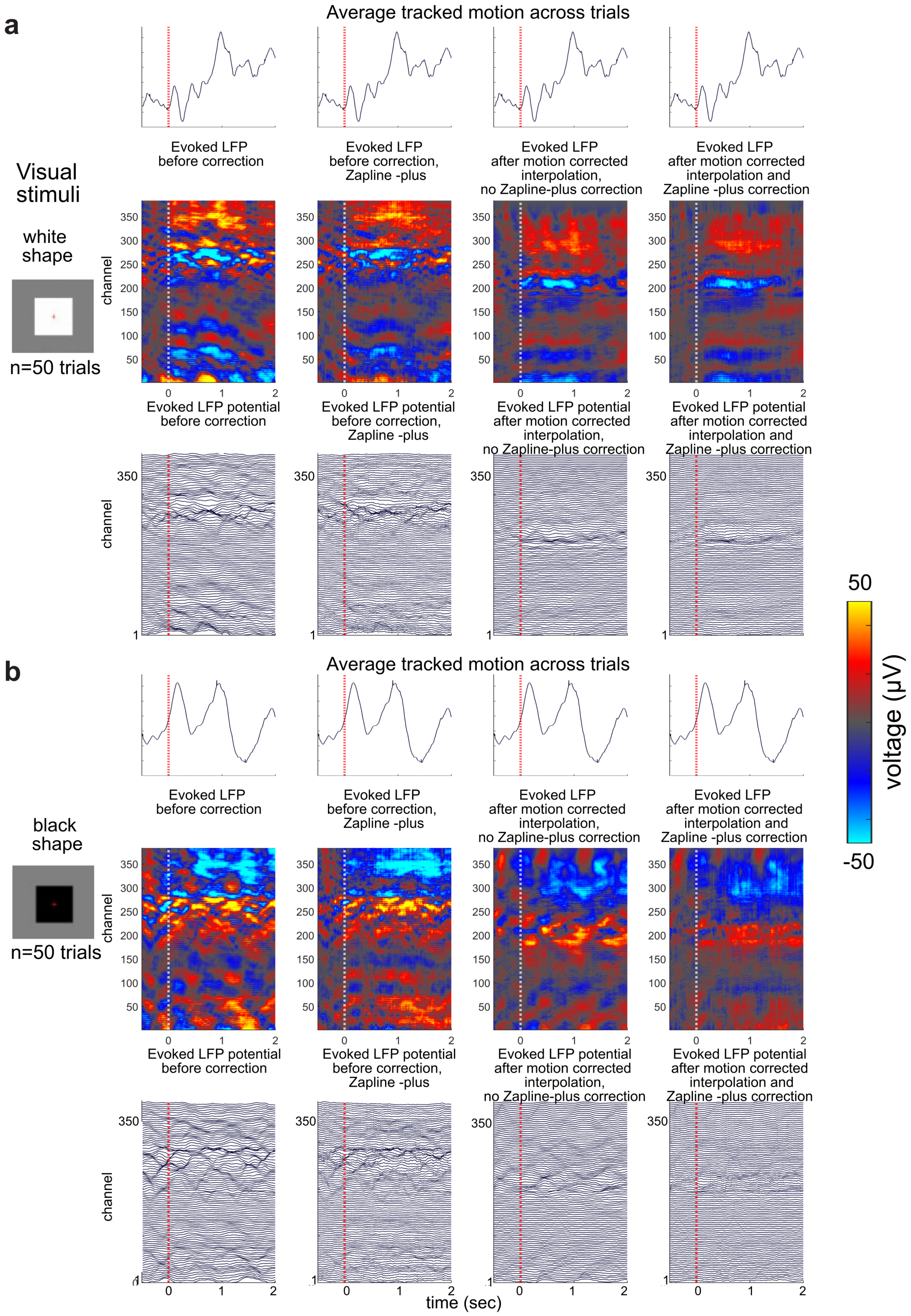
a
Average tracked motion across trials
Visual stimuli
white shape
n=50 trials
Evoked LFP before correction
Evoked LFP before correction, Zapline -plus
Evoked LFP after motion corrected interpolation, no Zapline-plus correction
Evoked LFP after motion corrected interpolation and Zapline -plus correction
channel
Evoked LFP potential before correction
Evoked LFP potential before correction, Zapline -plus
Evoked LFP potential after motion corrected interpolation, no Zapline-plus correction
Evoked LFP potential after motion corrected interpolation and Zapline -plus correction
b
black shape
voltage (µV)
50
-50
time (sec)

1456

Supplementary Figure 8: **Effects of motion correction interpolation and Zapline-plus applied to LFP on average visually evoked potentials in human brain activity. a**, Responses to the onset of white shapes (n=50 trials) in raw spontaneous LFP (left), raw data with Zapline-plus correction of low-frequency peaks (second column), motion-correction interpolation of LFP without Zapline-plus correction (third column), motion-correction interpolation of LFP with Zapline-plus correction (fourth column). **b**, Responses to the onset of black shapes (n=50 trials) in raw spontaneous LFP (left), raw data with Zapline-plus correction of low-frequency peaks (second column), motion-correction interpolation of LFP without Zapline-plus correction (third column), motion-correction interpolation of LFP with Zapline-plus correction (fourth column).

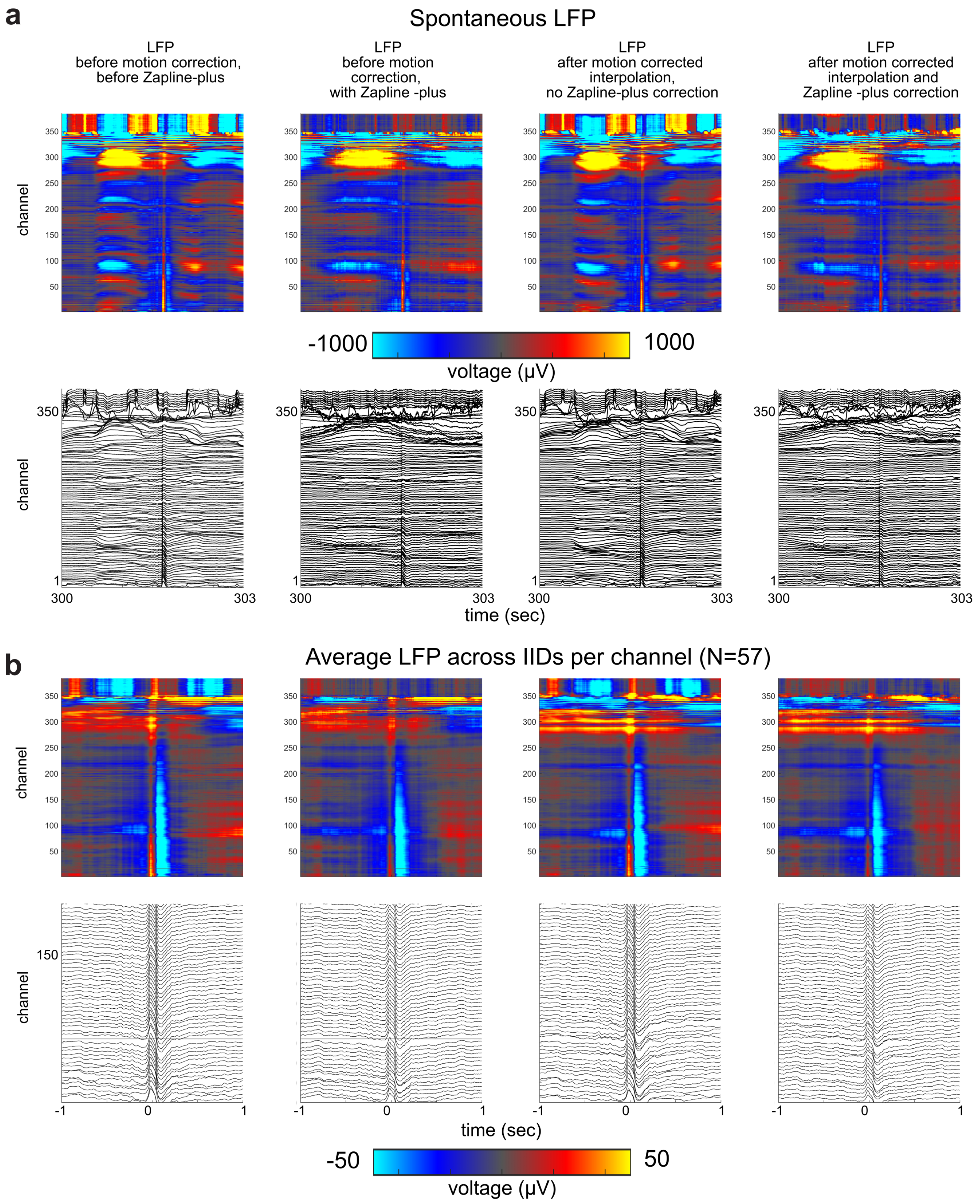


1457

Supplementary Figure 9: **Effects of interpolation and Zapline-plus steps on interictal epileptiform discharge (IID) waveforms.** **a**, Raw spontaneous IID events in the LFP (left), raw data with Zapline-plus correction of low-frequency peaks (second column), motion-correction interpolation of LFP without Zapline-plus correction (third column), motion-correction interpolation of LFP with Zapline-plus correction (fourth column), (N=1, Pt03). **b**, Mean detected IID events (n=57) in the LFP (left), raw data with Zapline-plus correction of low-frequency peaks (second column), motion-correction interpolation of LFP without Zapline-plus correction (third column), motion-correction interpolation of LFP with Zapline-plus correction (fourth column), (N=1, Pt03).

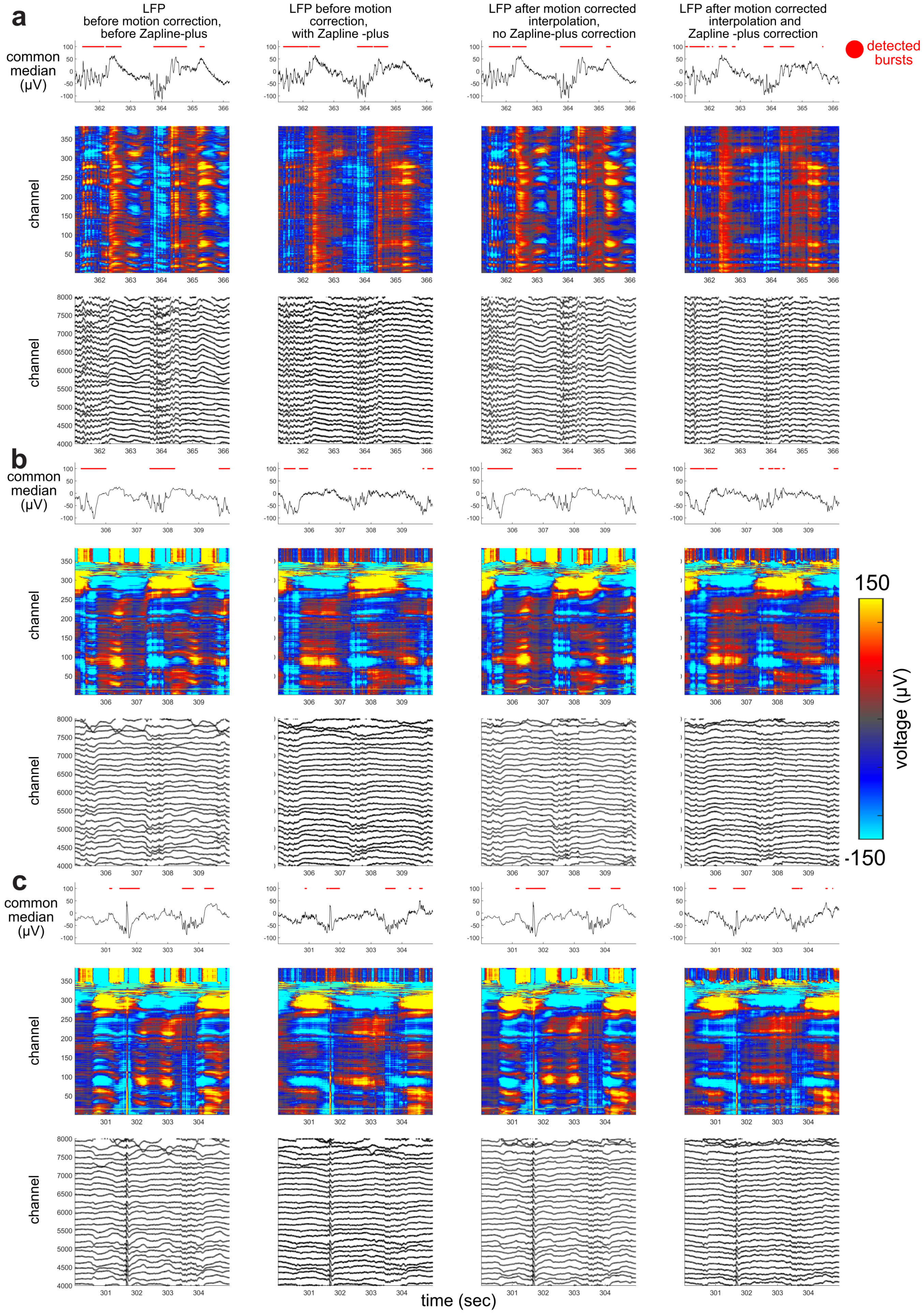
a
b
c
LFP
before motion correction,
before Zapline-plus
LFP before motion
correction,
with Zapline -plus
LFP after motion corrected
interpolation,
no Zapline-plus correction
LFP after motion corrected
interpolation and
Zapline -plus correction
detected
bursts
common
median
(µV)
channel
150
-150
voltage (µV)
time (sec)

Supplementary Figure 10: **Effects of interpolation and Zapline-plus steps on detected burst suppression signals during general anesthesia. a**, Raw spontaneous detected bursts (red dots) in the common median LFP (top) relative to the activity across channels (bottom two rows) for the raw LFP (left), raw data with Zapline-plus correction of low-frequency peaks (second column), motion-correction interpolation of LFP without Zapline-plus correction (third column), motion-correction interpolation of LFP with Zapline-plus correction (fourth column), (N=1, Pt01). **b**, Raw spontaneous detected bursts (red dots) in the common median LFP (top) relative to the activity across channels (bottom two rows) for the raw LFP (left), raw data with Zapline-plus correction of low-frequency peaks (second column), motion-correction interpolation of LFP without Zapline-plus correction (third column), motion-correction interpolation of LFP with Zapline-plus correction (fourth column), (N=1, Pt03). **b**, Raw spontaneous detected bursts (red dots) in the common median LFP (top) relative to the activity across channels (bottom two rows) for the raw LFP (left), raw data with Zapline-plus correction of low-frequency peaks (second column), motion-correction interpolation of LFP without Zapline-plus correction (third column), motion-correction interpolation of LFP with Zapline-plus correction (fourth column), (N=1, Pt03). This example includes a detected epileptiform IID.

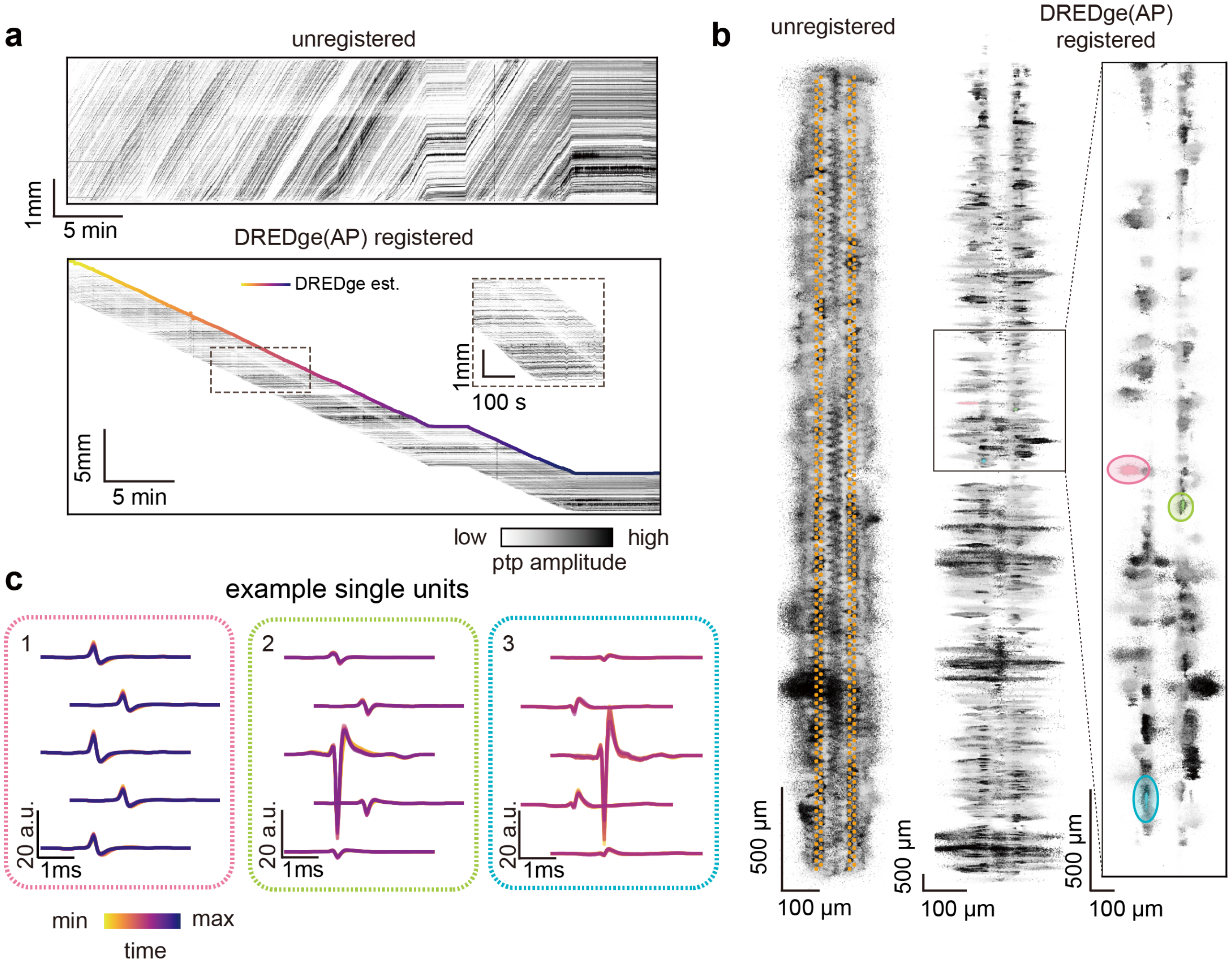


Supplementary Figure 11: **DREDge motion tracking for monkey 2. a** Spike raster before registration. (upper, scale bars: 1mm vertical, 5 min horizontal) and after registration (bottom, scale bars: 5000 µm (vertical), 5 min (horizontal).), overlaid by the DREDge estimated motion trace. **b** Localizations of detected spikes before (left) and after (right) drift correction according to DREDge motion estimation.(scale bars: 500µm vertical, 100µm horizontal) **c** Time binned (bin size = 15 s) templates with DREDge tracked motion correction for three example units ($N_1$ = 1935, $N_2$ = 611, $N_3$ = 639). Unit 2 and 3 lost track in the middle due to recording quality. (scale bars: 1ms, 20)

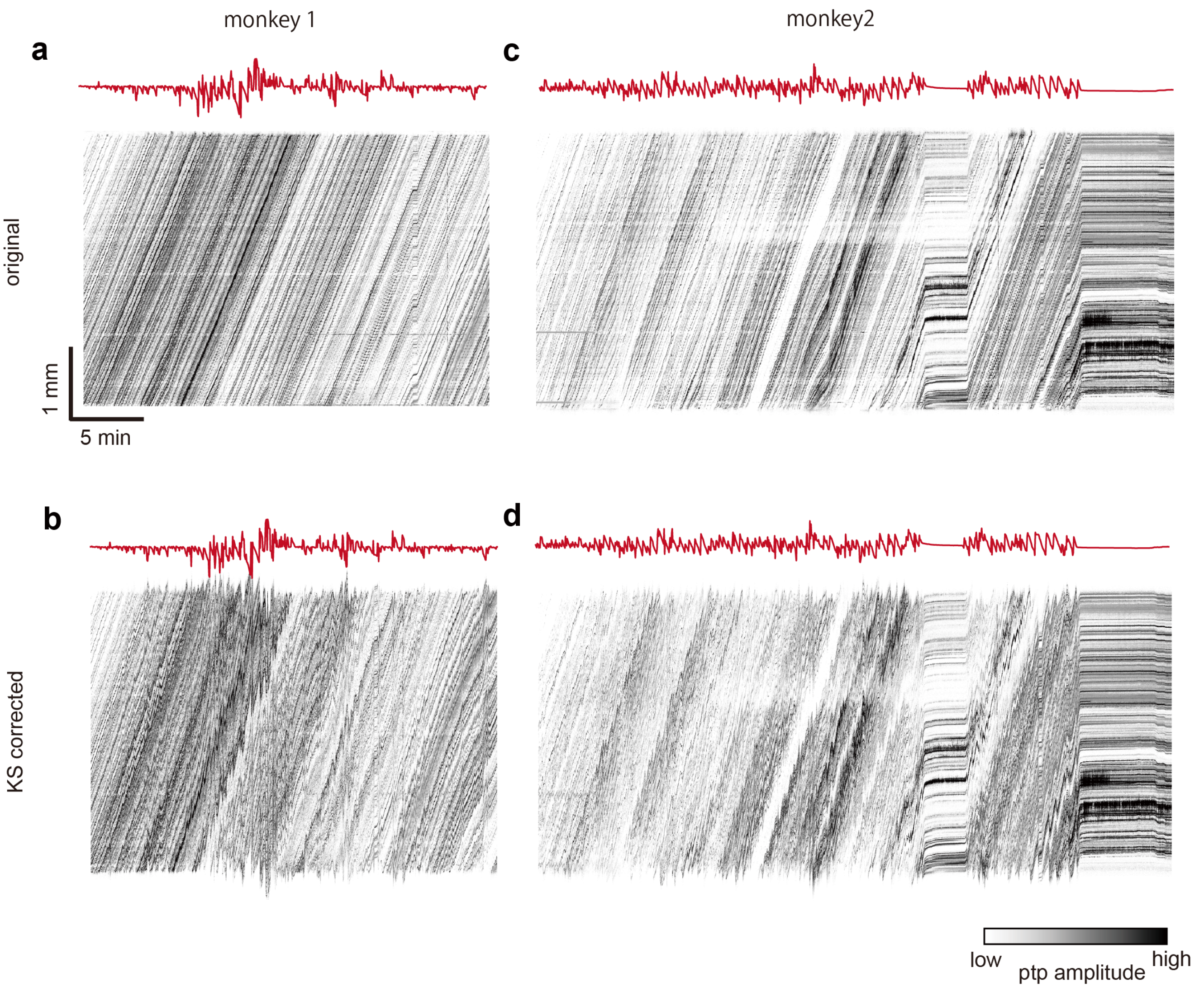


1460

Supplementary Figure 12: **Kilosort 2.5 failed to track long-range drift.** Spike rasters before (upper) and after (below) drift correction from Kilosort 2.5 motion estimation. The Kilosort 2.5 motion estimates are shown as red lines.

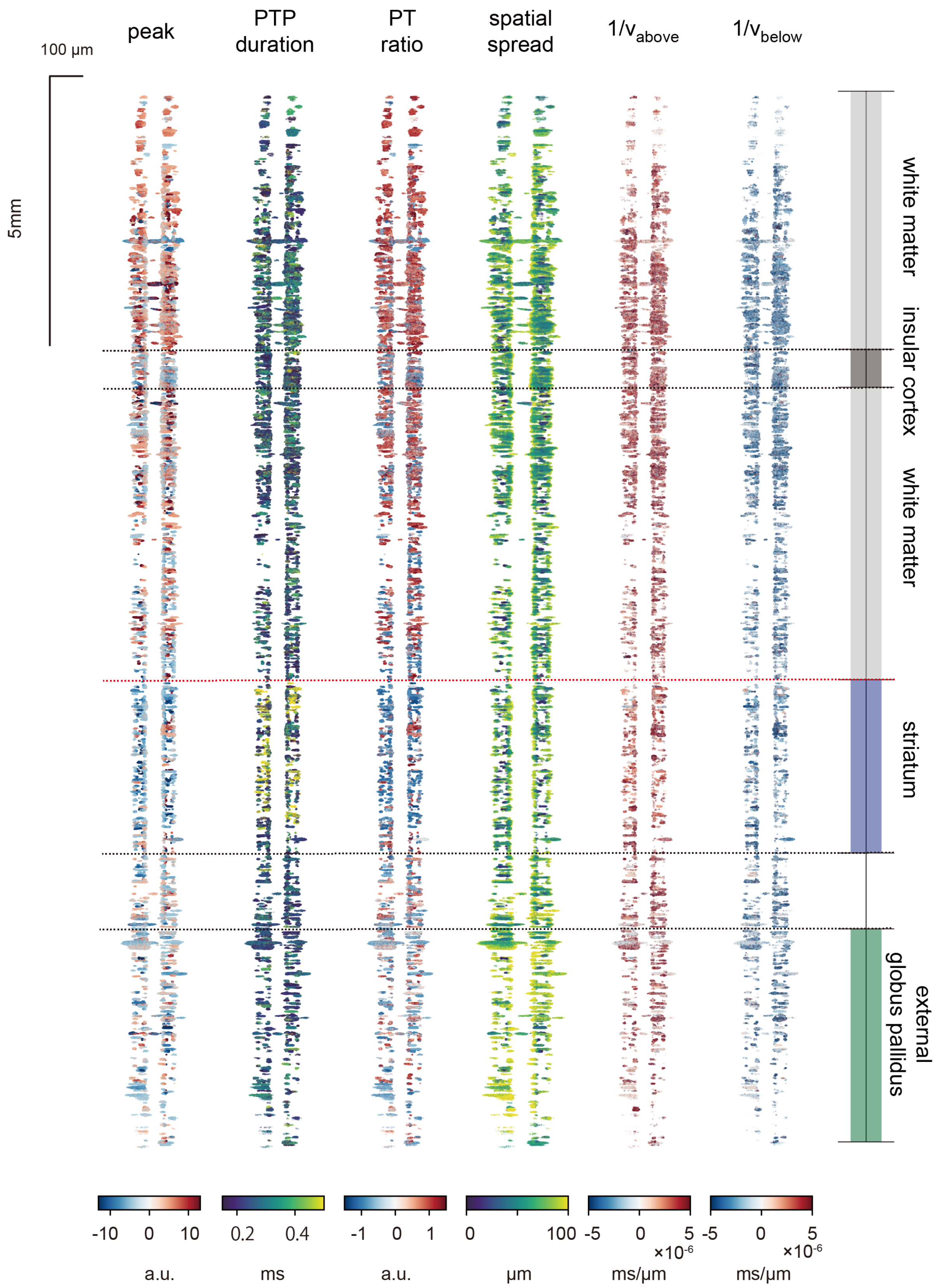


1461

Supplementary Figure 13: **Spike feature scatter plots.** Spatial scatter plots of the spike features of Fig. 4.f, visualizing the individual spikes whose features were averaged to create that panel.

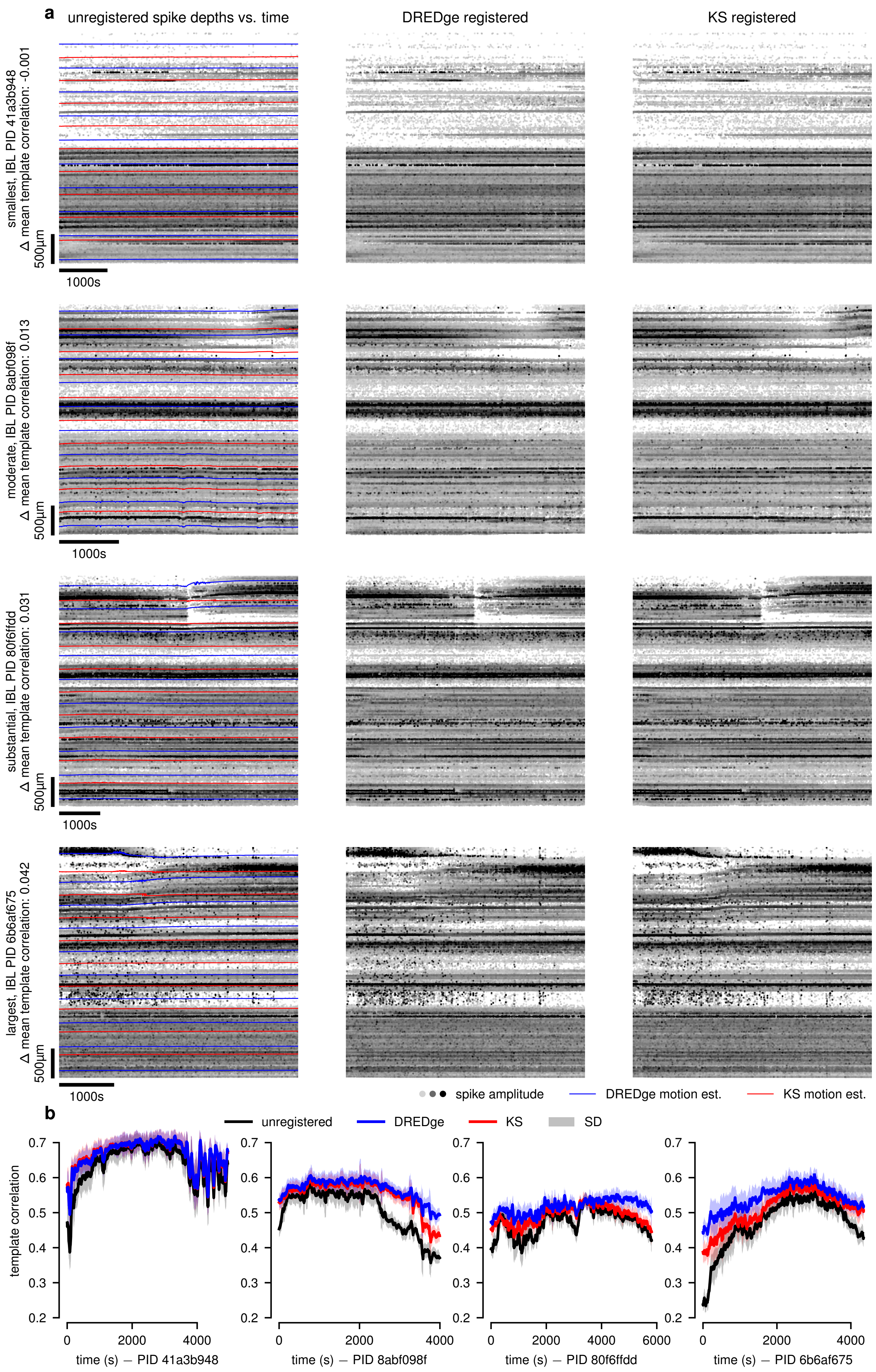
a
unregistered spike depths vs. time
DREDge registered
KS registered
smallest, IBL PID 41a3b948
Δ mean template correlation: -0.001
moderate, IBL PID 8abf098f
Δ mean template correlation: 0.013
substantial, IBL PID 80f6ffdd
Δ mean template correlation: 0.031
largest, IBL PID 6b6af675
Δ mean template correlation: 0.042
500μm
1000s
spike amplitude
DREDge motion est.
KS motion est.
b
unregistered
DREDge
KS
SD
template correlation
time (s) — PID 41a3b948
time (s) — PID 8abf098f
time (s) — PID 80f6ffdd
time (s) — PID 6b6af675

1462

Supplementary Figure 14: **Improved tracking in International Brain Lab data.** Here, we show example comparisons of DREDge motion estimation vs IBL's PyKilosort motion estimation[20] on publicly available IBL datasets[32]. **a** Each row corresponds to a single IBL probe insertion, and from left to right the columns show unregistered spike positions over time with motion traces overlaid (PyKS, red; DREDge, blue), DREDge corrected spike positions over time, and PyKS corrected spike positions over time. In the text on the left hand side, numerical values for the metric shown in Fig. 5.b ($\Delta$ mean template correlation, see also Section 4.9) are supplied, and these increase from the top row to the bottom row, showing visually what kind of relative improvements of DREDge over PyKS the metric quantifies. **b** Frame-by-frame template correlation traces for each registration of the four recordings in **a**. Lines show running means of template correlations of each frame and its 20 neighbors to either side, and the confidence band shows the standard deviation of these correlations.

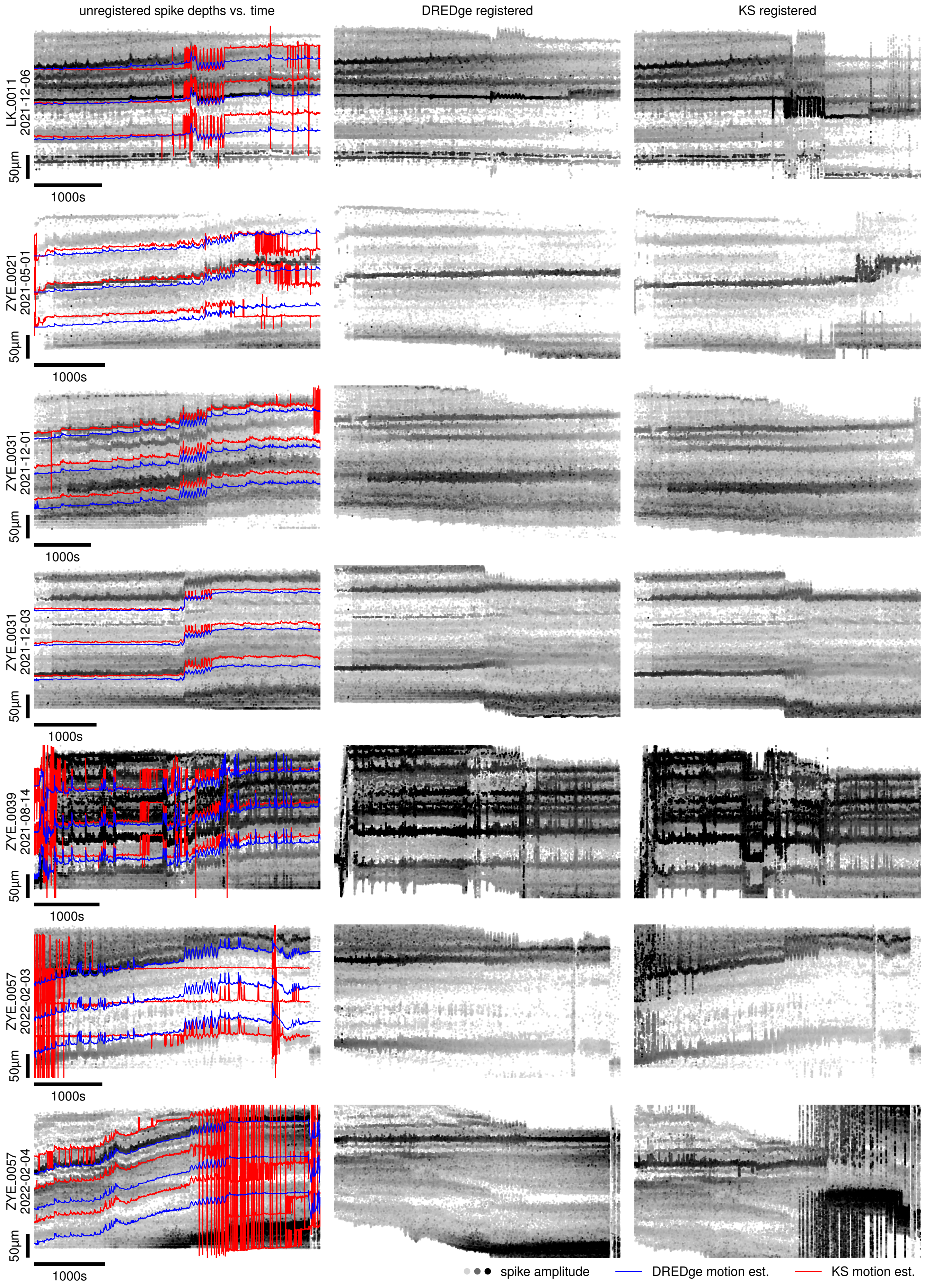

unregistered spike depths vs. time
DREDge registered
KS registered
LK_0011 2021-12-06
ZYE_0021 2021-05-01
ZYE_0031 2021-12-01
ZYE_0031 2021-12-03
ZYE_0039 2021-08-14
ZYE_0057 2022-02-03
ZYE_0057 2022-02-04
50µm
1000s
spike amplitude
DREDge motion est.
KS motion est.


1463

Supplementary Figure 15: **Improved tracking in Neuropixels Ultra probes.** In a subset of 7 of 12 (for space reasons) recordings with the Neuropixels Ultra probe[34], DREDge provides increased stability and fidelity in motion tracking in visual comparisons against Kilosort, as in Fig. 5.a.iii. Each row shows spikes from one NP Ultra dataset, each of which features both natural and imposed zig-zag motion, where the zig-zag motion typically begins around halfway (∼2000s) into the recording. Left column, unregistered spike positions plotted over time, with motion traces from DREDge and KS overlaid. Middle and right columns, spike positions offset by the DREDge and Kilosort motion estimates over time. KS' motion estimates (left column, red) often feature jump artifacts, which affect the quality of its registered raster plot (right column) relative to DREDge's more stable motion traces (left column, blue). Template correlation traces (see Section 4.9 for these motion estimates appear among those in Supp. Fig. 16.

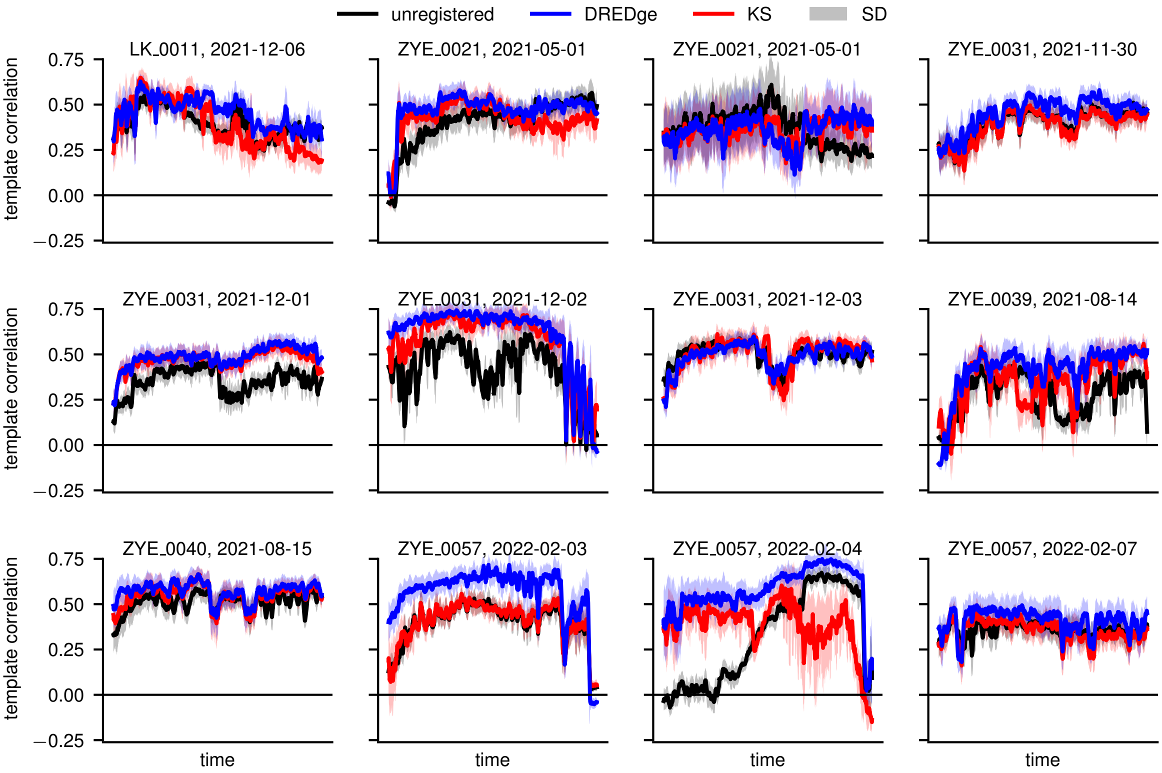


Supplementary Figure 16: **Measuring improved tracking in Neuropixels Ultra probes.** These panels accompany Supp. Fig. 15 by visualizing the frame-by-frame template correlation metric (see Section 4.9) corresponding to each registration of the 7 datasets of that figure along with the rest of the collection of NP Ultra zig-zag imposed motion recordings. Here, we plot the mean and standard deviation of the template correlation of each frame and its 20 neighbors to either side.

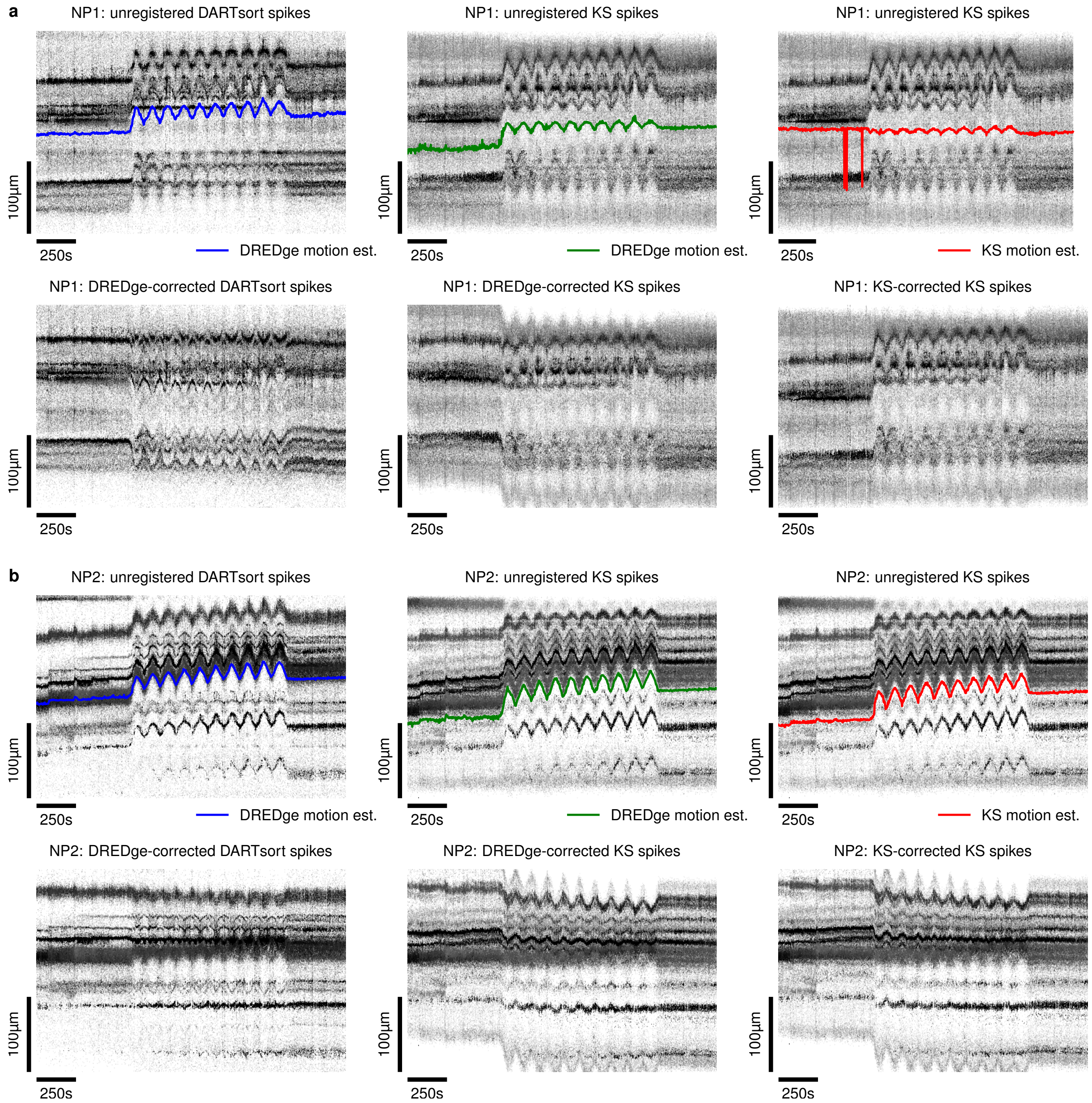


Supplementary Figure 17: **Improved tracking in smaller probes.** We trimmed the imposed motion recordings of Fig. 5.a to a set of channels with height less than that of the ultrahigh-density probe (282μm; Fig. 5.a, iii); **a** shows the trimmed NP1 recording, NP2 in **b**. We compared rigid motion estimation with DREDge and Kilosort 2.5. **a**,**b**: top left, unregistered spike localizations from the initial detection pipeline of[35], with DREDge motion estimate; bottom left, same spikes as above, with positions corrected according to the DREDge motion trace. Top middle, spike localizations from KS' initial detection with DREDge motion estimate; bottom middle shows DREDge-corrected positions for these spikes. Top right, spike localizations from KS with KS motion trace; bottom right shows KS' motion-corrected spike positions. **a**, DREDge's performance is better when using DARTsort's spikes than KS' spikes, and KS' motion tracking largely fails in this trimmed NP1 recording when compared to DREDge's tracking. **b**, KS' performance is on par with DREDge's when using KS' spikes, but DREDge improves when inputting DARTsort's spikes, which visually appear to be more robust to boundary effects.

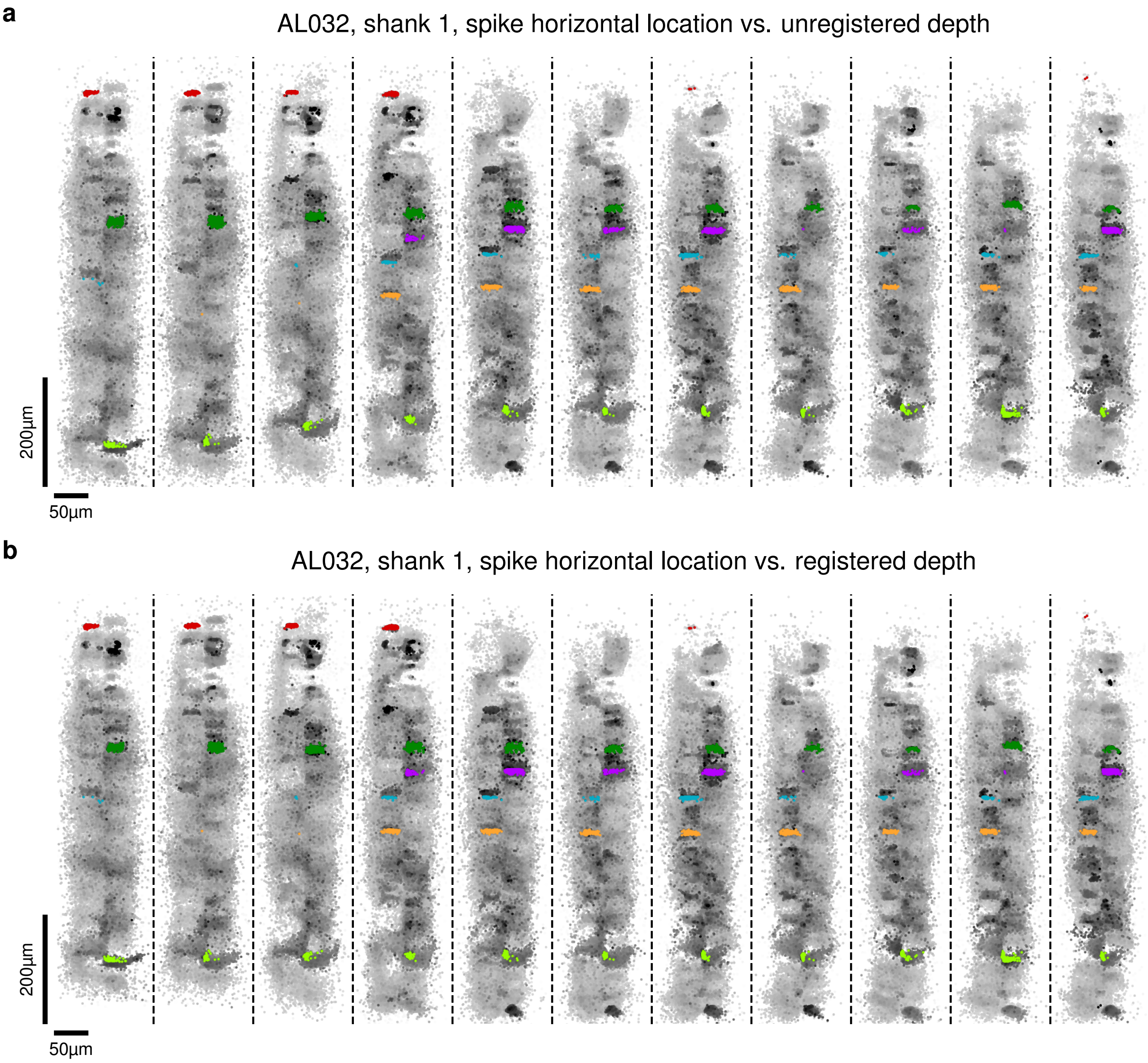


Supplementary Figure 18: **Horizontal and vertical spike positions used in manual clustering.** **a** Unregistered spike depths versus horizontal positions; **b** registered depths versus horizontal positions. Darker spikes have higher amplitudes. Colored spikes shown in both panels correspond to the manual clustering described in the text.

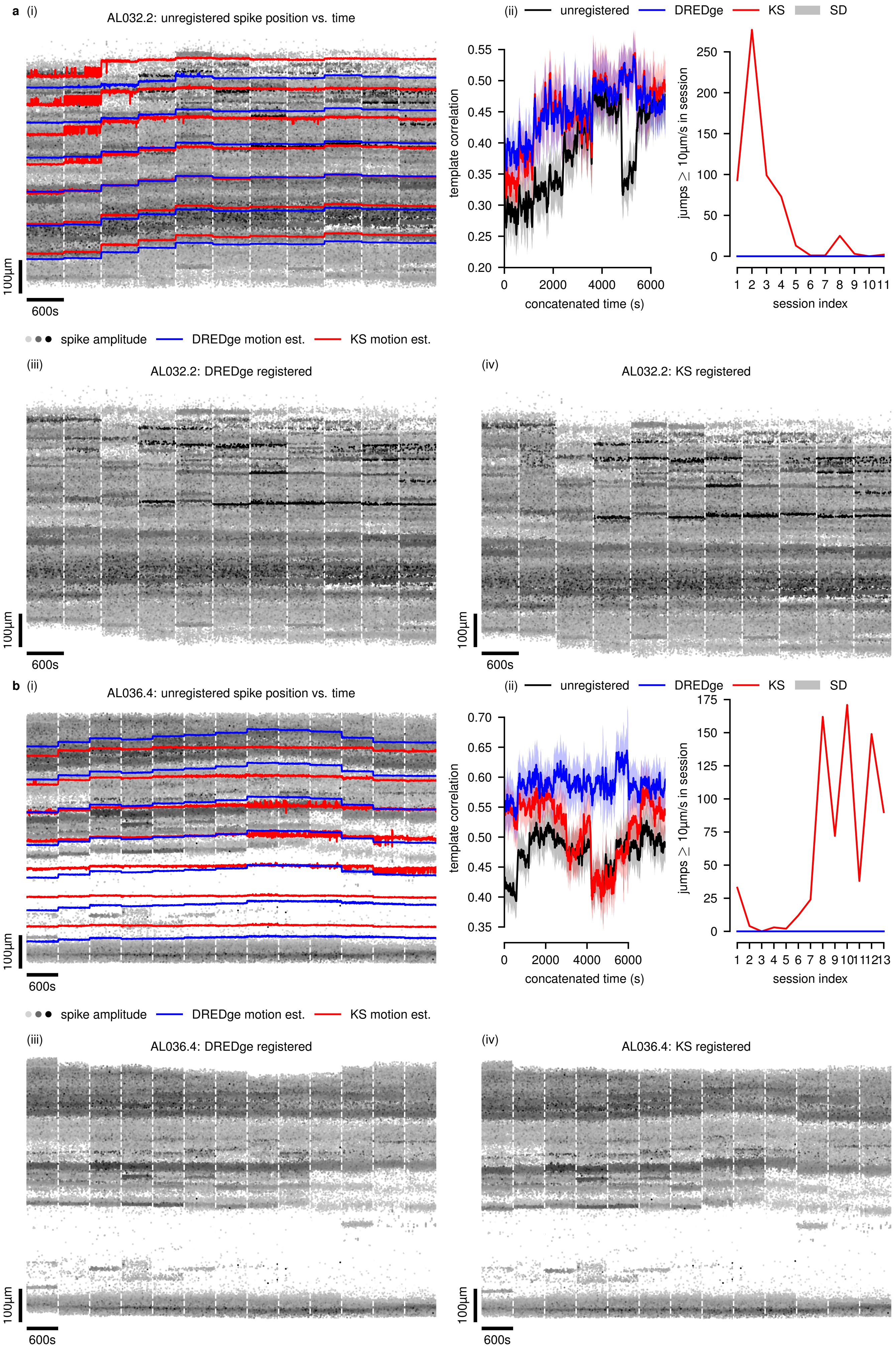
a (i)
AL032.2: unregistered spike position vs. time
100µm
600s
(ii)
unregistered
DREDge
KS
SD
template correlation
concatenated time (s)
jumps ≥ 10µm/s in session
session index
spike amplitude
DREDge motion est.
KS motion est.
(iii)
AL032.2: DREDge registered
(iv)
AL032.2: KS registered
b (i)
AL036.4: unregistered spike position vs. time
(iii)
AL036.4: DREDge registered
(iv)
AL036.4: KS registered

Supplementary Figure 19: **Chronic drift tracking in four-shank NP2 recordings.** Here, we visualized the results of drift tracking over days and weeks in data extracted from individual shanks in two chronic NP2 implantations: **a**, AL032 shank 2; **b**, AL036 shank 4[6]. In **a**,**b**, (i) shows the unregistered spike positions over time with estimated motion overlaid (DREDge, blue; KS2.5, red). In (ii), we present two metrics of registration quality: left, the correlation of each frame to the global template (see Section 4.9), visualized by running means and standard deviations of each frame with its 20 neighbors on either side; right, the number of registration time bins in which the estimated motion exceeded 10μm/s in each session. Respectively, (iii) and (iv) show the registered spike locations over time from DREDge and KS2.5.

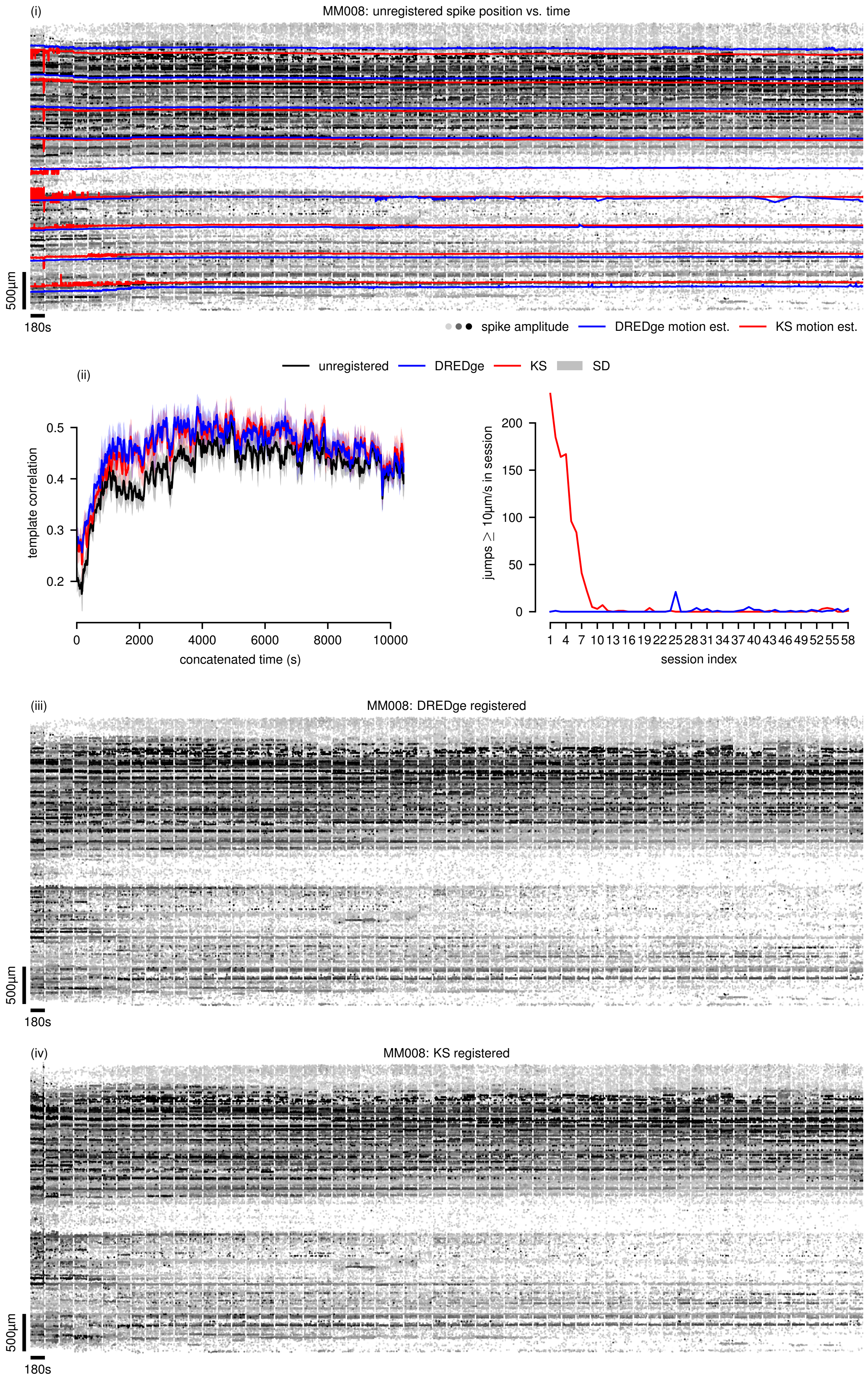
(i)
MM008: unregistered spike position vs. time
500μm
180s
spike amplitude
DREDge motion est.
KS motion est.
unregistered
DREDge
KS
SD
(ii)
template correlation
0.5
0.4
0.3
0.2
0
2000
4000
6000
8000
10000
concatenated time (s)
jumps ≥ 10μm/s in session
200
150
100
50
0
1 4 7 10 13 16 19 22 25 28 31 34 37 40 43 46 49 52 55 58
session index
(iii)
MM008: DREDge registered
500μm
180s
(iv)
MM008: KS registered
500μm
180s

Supplementary Figure 20: **Chronic drift tracking in an NP1 recording.** Here, we visualized the results of drift tracking over days and weeks in data extracted from a chronic NP1 implantation. (i) shows the unregistered spike positions over time with estimated motion overlaid (DREDge, blue; KS2.5, red). In (ii), we present two metrics of registration quality: left, the correlation of each frame to the global template (see Section 4.9), visualized by running means and standard deviations of each frame with its 20 neighbors on either side; right, the number of registration time bins in which the estimated motion exceeded 10µm/s in each session. Respectively, (iii) and (iv) show the registered spike locations over time from DREDge and KS2.5.

| | DREDge parameters | | | | | Kilosort 2.5 parameters | | |
|---|---|---|---|---|---|---|---|---|
| Experiment | `win_step_um` | `win_scale_um` | `win_margin_um` | `max_disp_um` | `max_dt_s` | `nblocks` | `nBinsReg1` | `nBinsReg2` |
| IBL | 400 | 450 | 0 | auto | 1000 | **5** | 15 | 5 |
| NHP | rigid | rigid | rigid | **1000** | **100** | **1** | 15 | 5 |
| chronic NP1 | 400 | 450 | 0 | auto | 1000 | **4** | 15 | 5 |
| chronic NP2.4 | **100** | 450 | **-150** | auto | 1000 | **4** | 15 | 5 |
| NP Ultra | **75** | **250** | **150** | auto | 1000 | **2** | 15 | **15** |
| human LFP | **800** | **1500** | **-750** | **500** | 1000 | N/A | N/A | N/A |

Supplementary Table 1: **Parameters listing for DREDge and Kilosort 2.5.** Here, bold indicates a non-default parameter choice. The experiment abbreviations in the left column refer to: IBL, the IBL datasets experiment of Fig. 5.b; NHP, the NHP experiments of Fig. 4 and Supp. Figs. 11 to 13; chronic NP1, the recordings MM007 and MM008 referred to in Fig. 6 and Supp. Fig. 20; chronic NP2.4, the four-shank NP2 recordings of Fig. 6 and Supp. Fig. 19; NP Ultra, the Neuropixels Ultra recordings of Fig. 5 and Supp. Fig. 15; human LFP, the human LFP recordings of Fig. 2.c, Supp. Fig. 4. In the IBL experiment, KS2.5 registration was carried out via its Python port pyKilosort, which uses the same algorithm. In the human LFP experiment, spike-based registration was not feasible, so KS2.5 parameters are not listed. In the `max_disp_um` column, "auto" indicates the default choice of `win_scale_um`/4. A negative choice for `win_margin_um` indicates the internal margin by which window centers are offset from the edges of the probe. In the NHP row, "rigid" is written to indicate parameters which are only relevant to nonrigid registration.
